# *In Silico* Molecular-Based Rationale for SARS-CoV-2 Spike Circulating Mutations Able to Escape Bamlanivimab and Etesevimab Monoclonal Antibodies

**DOI:** 10.1101/2021.05.18.444605

**Authors:** Erik Laurini, Domenico Marson, Suzana Aulic, Alice Fermeglia, Sabrina Pricl

## Abstract

The purpose of this work was to provide an *in silico* molecular rationale of the role eventually played by currently circulating S-RBD_CoV-2_ mutations in evading the immune surveillance effects elicited by the two Eli Lilly LY-CoV555/bamlanivimab and LY-CoV016/etesevimab monoclonal antibodies. The main findings from this study and shows that, compared to the wild-type SARS-CoV-2 spike protein, mutations E484A/G/K/Q/R/V, Q493K/L/R, S494A/P/R, L452R and F490S are predicted to be markedly resistant to neutralization by LY-CoV555, while mutations K417E/N/T, D420A/G/N, N460I/K/S/T, T415P, and Y489C/S are predicted to confer LY-CoV016 escaping advantage to the viral protein. A challenge of our global *in silico* results against the relevant experimental data resulted in an overall 90% agreement. This achievement not only constitutes a further, robust validation of our computer-based approach but also yields a molecular-based rationale for all relative experimental findings, and leads us to conclude that the current circulating SARS-CoV-2 and all possible emergent variants carrying these mutations in the spike protein can present new challenges for mAb-based therapies and ultimately threaten the fully-protective efficacy of currently available vaccines.

---

The 2019 Coronavirus disease (COVID-19)^1, 2^ elicited by the novel severe acute respiratory syndrome coronavirus 2 (SARS-CoV-2)^3^ has caused more than 157 million of confirmed infections globally, and caused more than 3.3 million deaths so far.^4^ This pandemic has also forced much of the world to enter an unprecedented sort of stand-by condition, with exceptional life-threating situations and unparalleled damage to the global economy. The ability of science and technology to deliver an effective, global solution to COVID-19 will be critical to restoring some semblance of normalcy, and the scientific community has responded commendably to this vital call. In particular, incomparable efforts have been and still are currently focused on the development of effective measures to further limit the spreading of SARS-CoV-2 infection and to treat already affected individuals. To date, drug development is under way; however, no proven effective therapies for this virus currently exist,^5^ while drugs that target the dysregulated cytokine responses (aka cytokine storms) characteristic of COVID-19 are available,^6^ although their clinical benefit is still a matter of debate.^7^ Meanwhile, different mRNA- or virus-based vaccines have received approval (and more are under clinical trial) and have so far provided effective and efficient protection against the disease,^8, 9^ making vaccination the key weapon in fighting the COVID-19 pandemic. Another promising approach is the isolation of SARS-CoV-2 neutralizing monoclonal antibodies (mAbs).^10, 11^ mAbs are immuno-therapeutics, which could i) potentially deliver immediate benefit in COVID-19 treatment, ii) act as passive prophylaxis until vaccines become globally available, and iii) serve as alternative therapeutic strategies in those populations where vaccines have been found to be less protective.^11, 12^ The recent findings that ansuvimab (mAb114) is a safe and effective treatment for symptomatic infection with Ebola virus is a notable example of the successful use of mAb therapy during an outbreak of infectious disease.^13^

Ab-based therapeutics directed against SARS-CoV-2 still present lights and shadows.^14^ Preclinical data and phase-III clinical studies indicate that mAbs could be effectively deployed for prevention or treatment during the viral symptoms phase of the disease.^15^ Cocktail formulations of two or more mAbs are preferred over single Ab preparations because these combinations may result in increased antiviral efficacy and viral escape prevention.^16-18^ However, Ab cocktails are complicate formulations,^19, 20^ and such approach likely involves increased production costs and quantities at a time when the supply chain is being pressured into meeting the high demand for COVID-19 therapeutics, vaccines, and therapeutic agents in general.

The multi-domain SARS-CoV-2 surface spike (S) protein^21-23^ - a trimeric class I fusion protein that mediates viral entry - is the focus of the current Ab discovery efforts. The S protein is composed of two subunits: S1, containing a receptor-binding domain (S-RBD_CoV-2_) that recognizes and binds the human receptor, the angiotensin-converting enzyme 2 (ACE2),^24-27^ and S2, which mediates viral cell membrane fusion by forming a six-helical bundle *via* the two-heptad repeat domain. Viral entry is initiated by the upward shift of the spike RBD at the protein’s apex which, in turn, promotes ACE2 binding (Figure 1, top panels). In addition, viral cell entry involves the S-protein priming operated by the cellular transmembrane serine protease 2 (TMPRSS2),^28^ along with other proteases,^29^ the removal of subunit S1, and the conformational reorganization of subunit S2; all these processes contribute to viral fusion with the cell and transfers of genetic material following receptor involvement.

**Figure 1.**
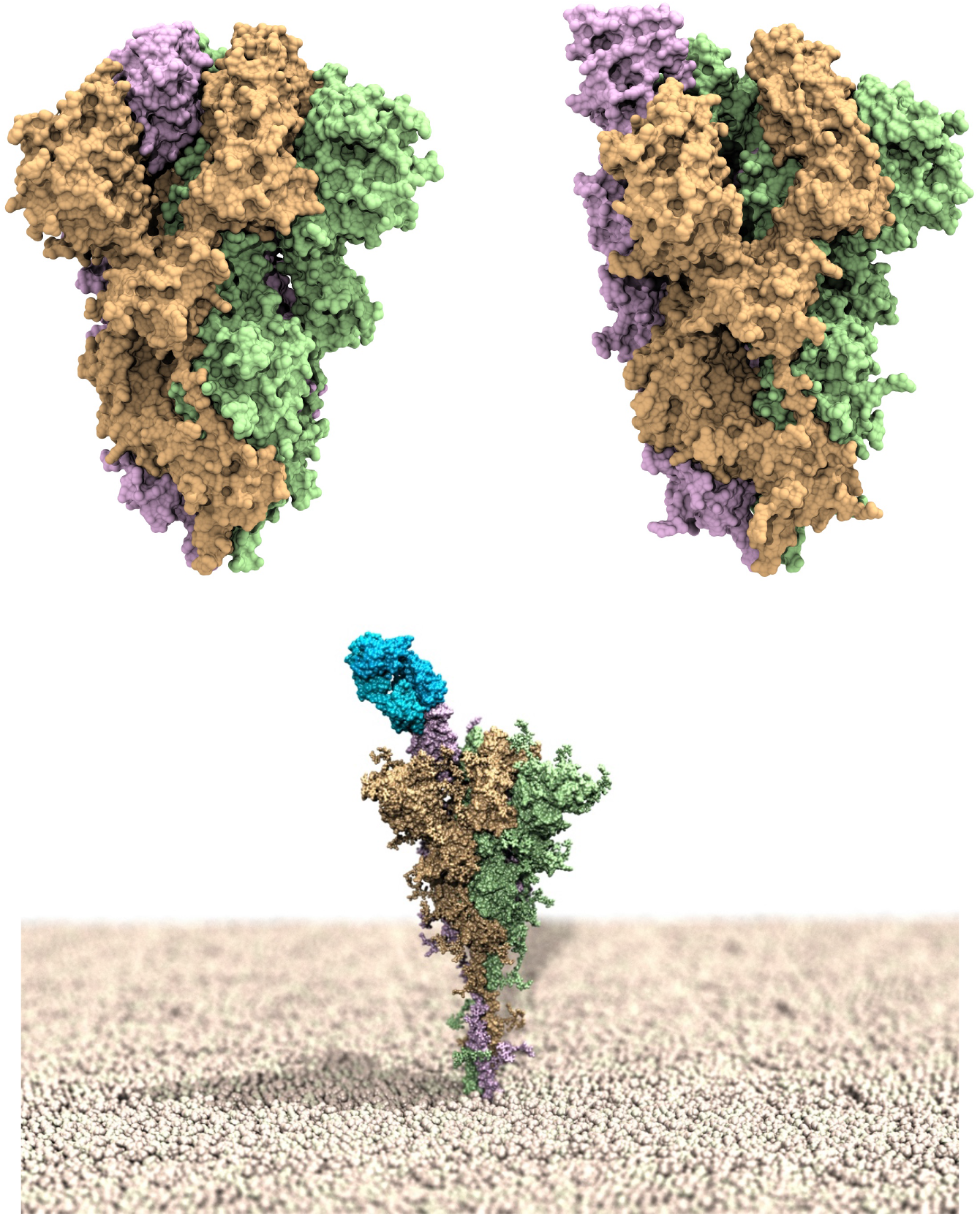
Upper panel: models of the SARS-CoV-2 spike homotrimeric protein in the down (left) and up (right) conformations. The three spike protomers are highlighted by their light green, tan and light purple van der Waals surfaces, respectively. Bottom panel: computer rendering of the full-length SARS-CoV-2 homotrimer embedded in a membrane model (polar heads in light tan spheres), showing one protomer in the up position and in complex with the LY-CoV555 (bamlanivimab) monoclonal antibody (light blue van der Waals surface).

Due to the critical nature of the viral S-RBD_CoV-2_ interaction with ACE2, Abs that bind this domain and interfere with ACE2 attachment can have potent neutralizing activity.^30-37^ An S-RBD_CoV-2_ specific mAb (LY-CoV555 or bamlanivimab) was discovered that can bind the RBD in both (up/active) (bottom panel in Figure 1) and (down/resting) conformations, and was reported to display high *in vitro* and *in vivo* protection potency, thereby supporting its development as a therapeutic for the treatment and prevention of COVID-19.^38^ In the same context, another mAb isolated from a COVID-19 convalescent patient – LY-CoV016 or etesevimab – was soon after reported.^39^ This mAb also showed specific SARS-CoV-2 neutralization activity by recognizing another epitope on the S-RBD_CoV-2_, and was found effective in vivo in both prophylactic and therapeutic settings.^39^ These two mAbs from Eli Lilly presently under clinical evaluation for the treatment and prevention of COVID-19, both alone and in cocktail formulations.^40-42^ Contextually, in the United States the Food and Drug Administration (FDA) has already granted Emergency Use Authorization (EUA) of the combined bamlanivimab/etesevimab cocktail as anti-SARS-CoV-2 monoclonal antibody therapeutic for the treatment of COVID-19, while the European Medicine Agency (EMA) also recently concluded that these two mAbs can be used together to treat confirmed COVID-19 in patients who do not require supplemental oxygen and who are at high risk of their COVID-19 disease becoming severe. EMA also analyzed the use of LY-CoV555 alone and concluded that, despite uncertainties around the benefits of monotherapy, it could be considered a treatment option.

However, viruses that encode their genome in RNA (*e*.*g*., SARS-CoV-2, the Human Immunodeficiency Virus (HIV) and the Influenza A Virus (IAV)), are prone to acquire mutations in time, mainly because of three factors. The first, and likely the most probable source of mutations consists in copying errors as viruses replicate inside host cells.^43^ Interestingly, however, this mechanism may be less relevant for SARS-CoV-2 with respect to other RNA viruses, since coronavirus polymerases – *i*.*e*., those enzymes the play vital role in viral genome replication and transcription – are endowed with a proofreading mechanism that corrects potentially fatal mistakes.^44^ Viral genomic variability may also originate from the recombination of two viral lineages coinfecting the same host.^45^ As a third factor, mutations can be induced by the host cell RNA-editing systems, which form part of host natural immunity.^46, 47^ A further element of complexity is reported in the recent work by Di Giorgio *et al*.,^47^ according to which both the adenosine deaminases acting on RNA (ADAR) and the apolipoprotein B mRNA editing catalytic polypeptide-like (APOBEC) families of proteins are involved in coronavirus genome editing - a process that may change the fate of both virus and patient. Whatever the case, the lesson learned from RNA virus genetics and epidemiology is that mutations are an inevitable consequence of being a virus.^48^ Yet, we also know that those mutations that adversely impact any of the vital steps of virus function are swiftly eliminated by natural selection. On the contrary, neutral variations and especially those mutations that endow the virus with a competitive advantage can reach high frequencies.

Thus far, a steadily increasing number of SARS-CoV-2 genetic variants have been emerging and circulating all over the world since the beginning of the COVID-19 pandemic. Although all proteins encoded in the SARS-CoV-2 RNA are continuously reported and catalogued in the plethora of databased and networks dedicated to COVID-19 genomic surveillance,^49^ the spike protein is the one more often and constantly reported mutated in the viral genomes sequenced worldwide.^50^ In this respect, both the United States Center for Disease Control and Prevention (CDC) and the equivalent European agency (ECDC), in collaboration with the World Health Organization (WHO) and several governmental authorities and working groups have developed a SARS-CoV-2 variant classification schema that groups all major viral variants into main groups: variants of high consequence (VOHC, CDC), variants of concern (VOC), variants of interest (VOI) and variants under monitoring (VUM, ECDC), depending on their associated degree of impact on transmissibility, severity and/or reduced neutralization by antibody/efficacy of Ab treatments (of note, European and US classification may not fully coincide since the importance of variants may differ by location). At present no SARS-CoV-2 circulating variants have been classified as VOHC, while a substantial number them have been marked as VOC and VOI/VUM by CDC^51^ and ECDC,^52^ and all of them are characterized by the presence of at least one spike mutation of interest (MOI)^52^ (see the Conclusion section for a more detailed discussion on this subject). The ability of SARS-CoV-2 mAbs to select any of these variants that is apparently fit and that naturally occurs even at low frequencies in circulating viral populations suggests that the therapeutic use of single mAb might select for escape mutants, although the extent to which resistance will impact the effectiveness of Abs in SARS-CoV-2 therapeutic and vaccine settings is still a matter of intense investigation.^10, 53-55^ In this arena, the purpose of this work is to provide an atomistic-based, *in silico* perspective of the role eventually played by currently circulating S-RBD_CoV-2_ mutations in escaping binding of the two mAbs bamlanivimab and etesevimab as a proof of concept. A computational alanine scanning (CAS) mutagenesis^56^ initially allowed us to identify the main molecular determinants of each Ab/S-RBD_CoV-2_ recognition; then, each spike residues that, according to the CAS results, contributes to the relevant viral protein/mAb binding interface was mutated into all currently reported circulating mutations at that position,^57^ and the corresponding variation in affinity of all mutated spikes for each mAb with respect to the wild-type viral protein was estimated using a consolidate protocol.^58^ To quickly and effectively ranking the different spike mutants with respect to their mAb escaping potential, a color-coded criterion based on the predicted free energy difference range of values was adopted, as shown in Table 1. Of note, this criterion is identical to the one we adopted and validated in our previous work for ranking the effect of both ACE2 and S-RBD_CoV-2_ mutations on their mutual binding.^58^

**Table 1.** Color-coded criterion based on the predicted free energy difference (ΔΔG) range of values adopted to rank the affinity of SASR-CoV-2 spike mutants for the LY-CoV-555 (bamlanivimab) and LY-CoV016 (etesevimab) monoclonal antibodies. Negative/positive ΔΔG values indicate unfavorable/favorable substitutions for the mutant residue in the relevant position, respectively.

| Mutation effect | $\Delta\Delta G$ range (kcal/mol) | Color code |
| --- | --- | --- |
| Neutral mutations | $-0.25 \leq \Delta\Delta G \leq +0.25$ | Gray |
| Mildly destabilizing mutations | $-2.00 \leq \Delta\Delta G < -0.25$ | Light Yellow |
| Destabilizing mutations | $-4.00 \leq \Delta\Delta G < -2.00$ | Light Red |
| Highly destabilizing mutations | $-4.00 < \Delta\Delta G$ | Red |

## RESULTS AND DISCUSSION

### Computational Alanine Scanning of the SARS-CoV-2 Spike Protein Residues at the Binding Interface with the LY-CoV555 (Bamlanivimab) Monoclonal Antibody

Within distance and energetic cutoffs of 4.0 Å and 0.5 kcal/mol, respectively, the analysis of the equilibrated molecular dynamics (MD) trajectory of LY-CoC555 in complex with the S-protein RBD of SARS-CoV-2 shows that a total of 10 residues of S-RBD_CoV-2_ stably and effectively contact 19 residues of the fragment antigen-binding (Fab) portion of the LY-CoV555 mAb, 14 of which locate on the heavy chain (HC) and 5 on the light chain (LC), respectively (Figure 2A and Table S1).

**Figure 2.**
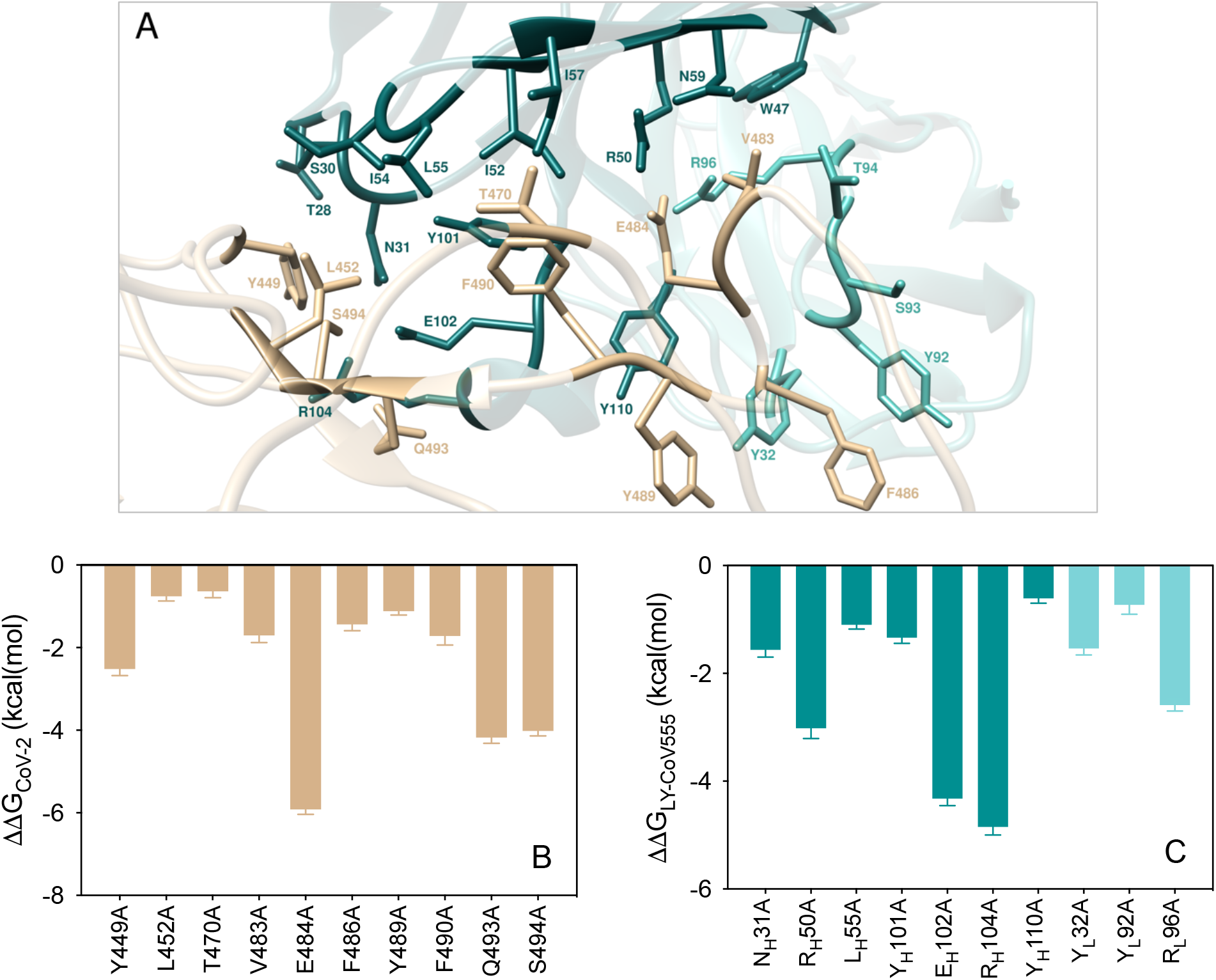
(A) Structural details of the binding interface between the LY-CoV555 (bamlanivimab) mAb and the viral spike protein receptor-binding domain of SARS-CoV-2 (S-RBD_CoV-2_). The secondary structures of LY-CoV555 and S-RBD_CoV-2_ are portrayed as light teal and light Tiffany ribbons, respectively. Each interacting protein residue is highlighted in dark matching-colored sticks and labeled. Binding energy change (ΔΔG = ΔG_WILD-TYPE_ − ΔG_ALA_) obtained from the computational alanine-scanning (CAS) mutagenesis for the S-RBD_CoV-2_ residues at the binding interface with the LY-CoV555 mAb (B) and for the LY-CoV555 mAb residues at the binding interface with the viral protein RBD (C). Negative ΔΔG values indicate unfavorable substitution for alanine in the relevant position. For the numerical values of ΔΔG and all related energy terms, see the text and Tables S2-S3.

The results from the CAS (Figure 2B-C) identify both the S protein and the LY-CoV555 mAb residues that afford a significant contribution to the binding interface. Furthermore, CAS data clearly indicate residues E484, Q493 and S494 on the S-RBD_CoV-2_ and R_H_50, R_H_96, E_H_102, and R_H_104 on the mAb heavy chain as key positions contributing to shaping and determining the stability of the relevant protein−protein interface, as discussed in details below. *E484*. The confirmation of the E484 as a crucial residue was an expected result as a glutamic acid (E) to lysine (K) substitution at this position (E484K) in the S-RBD_CoV-2_ is present in the rapidly spreading variants of concern belonging to the B.1.351 (aka South African) and P.1 (Brazilian) lineages, while the E484Q/L452R double mutation is a component of the B.1.617 lineage that is currently dramatically spreading in India (*vide infra*). E484 locates at the tip of a long, flexible loop in the S-RBD_CoV-2_; as such, any intermolecular interaction involving E484 and LY-CoV555 could be important in eventually anchoring the entire superstructure. The MD trajectory of the S-RBD_CoV-2_/LY-CoV555 complex shows that E484 is involved in two tight and bifurcated salt-bridges with residues R_H_50 (2.74 ± 0.10 Å and 3.05 ± 0.14 Å) and R_L_96 (2.82 ± 0.11 Å and 2.96 ± 0.13 Å) on the mAb HC and LC, respectively, flanked by contact interactions (CIs) with the side chains of Y_H_110 and Y_H_101 (Figure 3A, Table S1). When E484 is replaced with alanine in CAS, these interface-stabilizing interactions - along with the slightly beneficial contribution from the intramolecular van der Waals contact with the two Ab HC tyrosines - are no longer made, reflecting a loss of the corresponding binding free energy of ΔΔG_CoV-2_(E484A) = −5.92 ± 0.12 kcal/mol (Figure 2B, Table S2). Contextually, the corresponding values of ΔΔG_LY-CoV555_(R_H_50A) = −3.02 ± 0.19 kcal/mol and ΔΔG_LY-CoV555_(R_L_96A) = −2.59 ± 0.11 kcal/mol (Figure 2C, Table S3) are in line with the important contribution these residues provide to the formation of the corresponding viral protein/antibody interface described above (Figure 3A).

**Figure 3.**
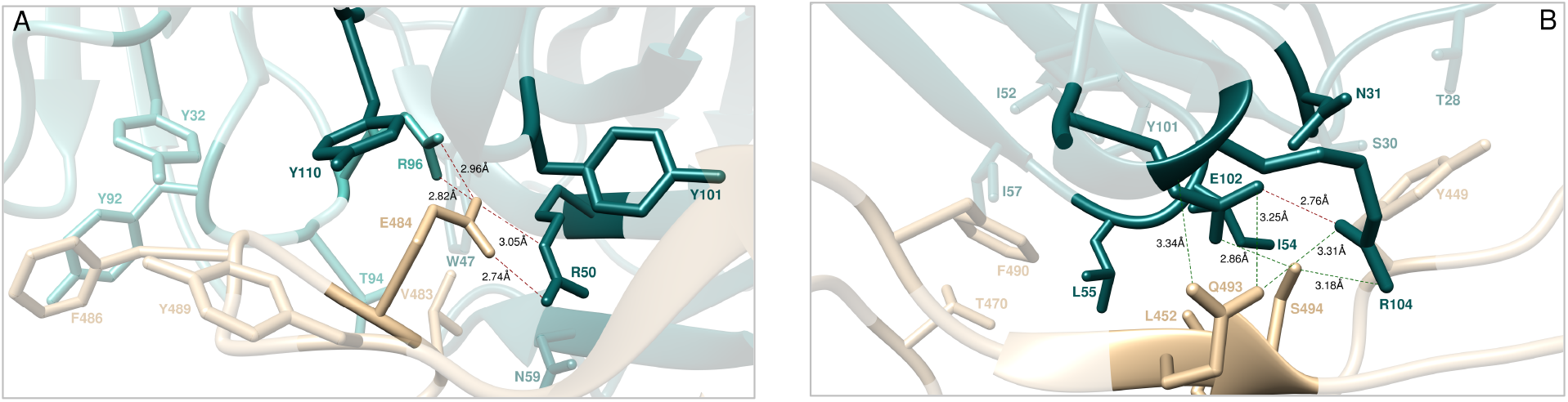
Main interactions involving the viral S-RBD_CoV-2_ residues E484 (A), Q493 and S494 (B), and L452 (C) at the interface with the LY-CoV555 (bamlanivimab) mAb as obtained from equilibrated MD simulations. In this and all remaining figures, the secondary structures of the S-RBD_CoV-2_ is shown as a light tan ribbon, while the HC and LC of the LY-CoV555 mAb are portrayed as light teal and light Tiffany ribbons, respectively. Each S-RBD_CoV-2_ residue under discussion and all other residues directly interacting with it are highlighted in dark matching-colored sticks and labeled; further residues/interactions related to the residue under investigation are evidenced in light matching-colored sticks and labelled in light gray. Hydrogen bonds (HBs) and salt bridges (SBs) are represented as dark green and dark red broken lines, respectively, and the relevant average distances are reported accordingly. Further important HBs and SBs detected in each complex are also indicated using light green/red broken lines and light gray labels (see Table S1 for details).

#### Q493

At the 493 position of the SARS-CoV-2 S protein, Q493 forms three stabilizing HBs across the protein-protein interface, one with the side chain of LY-CoV555 E_H_102 (3.25 ± 0.18 Å) and two with the side chain and the C=O backbone of R_H_104 (3.31 ± 0.12 Å and 3.34 ± 0.17 Å), respectively (Figure 3B, Table S1). Thus, abrogation of these intermolecular contacts by replacing the wild-type glutamine with alanine is accompanied by a ∼4.2 kcal/mol loss in binding free energy (ΔΔG_CoV-2_(Q493A) = −4.18 ± 0.14 kcal/mol, Figure 2B, Table S2). Similarly, when either of the two LY-CoV555 residues E_H_102 or R_H_104 are mutated into alanine, the related values of ΔΔG nicely reflect their importance in binding S-RBD_CoV-2_, as ΔΔG_LY-CoV555_(E_H_102A) = −4.32 ± 0.13 kcal/mol and ΔΔG_LY-CoV555_(R_H_104A) = −4.58 ± 0.15 kcal/mol, respectively (Figure 2C, Table S3). Of note, these two mAb amino acids are engaged in a fundamental internal SB (2.76 ± 0.11 Å, Figure 3B, Table S1) that appears to play a major structural role in properly orienting their side chains for binding both Q493 and another spike key residue – S494 – as discussed below.

#### S494

Serine 494 is an interesting SARS-CoV-2 RBD residue that has been previously reported by us to form an internal HB with the side chain of the adjacent Q493, instrumental to direct the latter in H-bridging aspartic acid 35 (D35) on the human ACE2 across their binding interface.^56, 58^ When in complex with LY-CoV555, S494 engages the side chains of the LY-CoV555 mAb residues E_H_102 and R_H_104 in two intermolecular HBs (2.86 ± 0.16 Å and 3.18 ± 0.19 Å, respectively), alongside a strong polar interaction with the viral N31 (Figure 3B, Table S1). Thus, the S494A mutation actually shows a considerable variation in the corresponding ΔΔG value (ΔΔG_CoV-2_(S494A) = −4.02 ± 0.12 kcal/mol, Figure 2B, Table S2) making S494 the spike third protein/protein key residue.

#### V483, F486, and Y489

Predictably, the SARS-CoV-2 RBD residues V483, F486, and Y489 afford only a network of stabilizing intermolecular CIs to the viral protein/antibody binding interface region centered around the nearby key residue E484 (Figure 3A). Specifically, the side chain of V483 interacts – *via* van der Waals/hydrophobic contacts – with the side chains of W_H_47, R_H_50 and N_H_59 on the Ab HC, and the side chains of T_L_94 and R_L_96 on the Ab LC, respectively (Figure 3A, Table S1). Contextually, the phenyl ring of F486 engages two π/π stacking interactions involving the side chains of the Ab LC Y_L_32 and Y_L_92, while the aromatic moiety of Y489 establish dispersive/polar interactions with the LY-CoV555 residues Y_H_110 and Y_L_32 on the Ab HC and LC, respectively (Figure 3A, Table S1). The absence of these CIs when each of these residues is mutated into alanine reflects the moderate variations of the corresponding free energy of binding (Figure 2B, Table S2), that is, ΔΔG_CoV-2_(V483A) = −1.70 ± 0.18 kcal/mol, ΔΔG_CoV-2_(F486A) = −1.44 ± 0.15 kcal/mol, and ΔΔG_CoV-2_(Y489A) = −1.12 ± 0.09 kcal/mol, respectively.

#### Y449, L452, T470, and F490

In analogy to what just discussed a few lines above, the main role of residues Y449, L452, T470 and F490 on the S-RBD_CoV-2_ is also to reinforce the viral protein/antibody binding interface region centered – in this case – around the two other important residues Q493 and S494 by providing a number of favorable intermolecular CIs (Figure 3B). In particular, Y449 provides three stabilizing polar interactions with the side chains of T_H_28, S_H_30, and N_H_31, and is in van der Waals distance with I_H_54, all on the Ab HC (Figure 3B, Table S1). L452 contacts the side chains of I_H_54 and L_H_55 on the LY-Cov555 Ab HC, while the last two spike residues T470 and F490 exchange nonpolar interactions with the side chains of the Ab HC residues I_H_52, I_H_54, L_H_55 and I_H_57, in addition to the π/π stacking observed between F490 and the Ab HC Y_H_101 (Figure 3B, Table S1). This is supported by the calculated ΔΔG value obtained by changing these amino acids into alanine in the S-RBD_CoV-2_/LY-CoV555 Ab complex, that is, ΔΔG_CoV-2_(Y449A) = −1.93 ± 0.16, ΔΔG_CoV-2_(L452A) = −0.76 ± 0.11 kcal/mol, ΔΔG_CoV-2_(T470A) = −0.64 ± 0.15, and ΔΔG_CoV-2_(F490A) = −2.38 ± 0.22 (Figure 2B, Table S2).

### *In silico* mutagenesis of the SARS-CoV-2 Spike Protein Residues at the Binding Interface with the LY-CoV555 (Bamlanivimab) Monoclonal Antibody

The recent survey of data reported by Starr *et al*.^57^ led to the following list of naturally occurring mutations at the SARS-CoV-2 S protein residues contacting the LY-CoV555 mAb: E484A/D/G/K/Q/R/V, Q493H/K/L/R, S494A/P/R/T, L452M/Q/R, Y449D/F/H/N/S, T470A/I/K/N, V483A/F/G/I/L, F486I/L/S, Y489C/F/H/S, and F490L/S/V/Y. In what follows, we report and discuss different effects exerted by each of these spike mutant residues on the structure and strength of the resulting S-RBD_CoV-2_ /LY-CoV555 binding interface. In analogy with our previous work focused on the estimation of the difference in binding affinity between different allelic variants of ACE2 or S-RBD_CoV-2_,^58^ in this study we adopt the same color-coded criterion based on the predicted free energy difference range of values shown in Table 1.

#### E484

The CAS results discussed above clearly show that glutamic acid at the position 484 along the S protein wild-type sequence (E484) is a key player in the LY-CoV555/S-RBD_CoV-2_ interaction (Figures 2B and 3A, Tables S1-S2). Interesting, replacing the viral spike E484 with each of the alternative residues considered (*i*.*e*., E484A/D/G/K/Q/R/V) reflects into a robust interface disrupting behavior, with the mild exception of the E484D substitution (Figure 4A, Figure S1, and Tables S4-S5). Figure 4B shows the results for the E484K as a representative example. As seen from this Figure, in the presence of the K484 mutation the two topical, bifurcate interface-stabilizing SBs between E484 and the side chains of LY-CoV55 R_H_50 and R_L_96 (Figure 3A, Table S5) cannot obviously be established, while the background network of CIs involving residues V483, F486, and Y489 on the S protein and W_H_47, Y_L_92, Y_H_110 and Y_H_32 on the LY-CoV555 mAb remains almost unperturbed (Figure 4B, Figure S1, Table S5).

**Figure 4.**
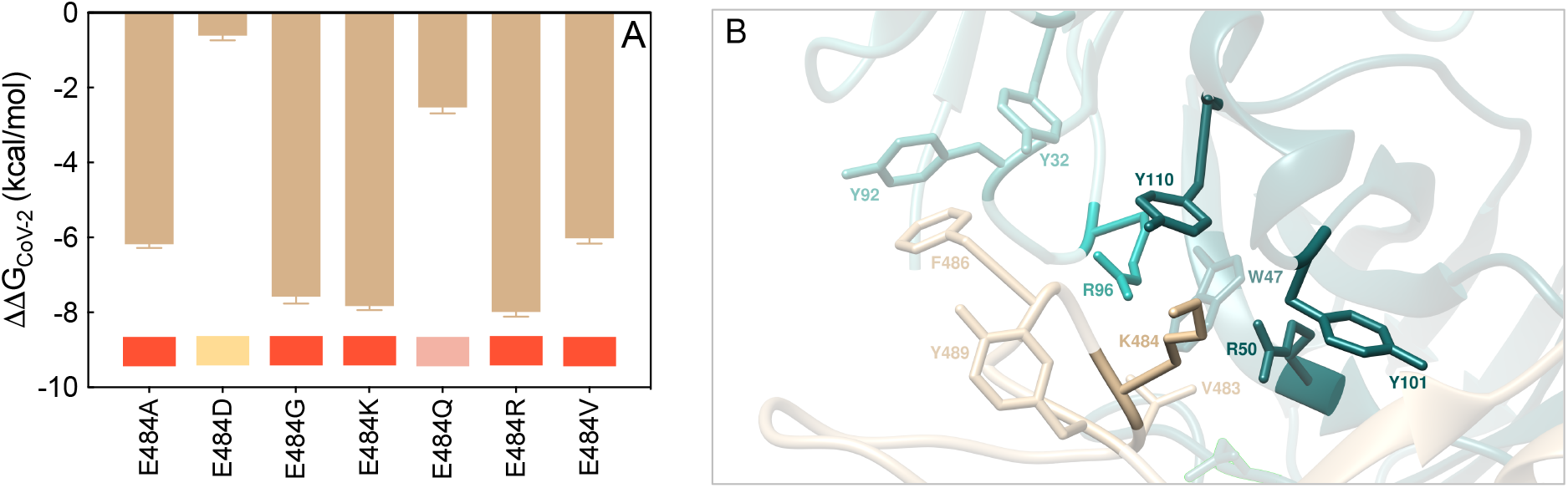
(A) Change in binding free energy (ΔΔG = ΔG_WILD-TYPE_ − ΔG_MUTANT_) predicted by computational mutagenesis of the S-RBD_CoV-2_ wild-type residue E484 for the corresponding S-RBD_CoV-2_/LY-CoV555 mAb complex. Negative ΔΔG values indicate unfavorable substitutions for the mutant residue in the relevant position. The numerical values of ΔΔG, all related energy terms, and all underlying intermolecular interactions are reported in Table S4, Figures S1, and Table S5. In this and all other similar figures, the colored boxes below each bar in the graphs show the classification of the destabilizing effects of the corresponding mutation on the S-RBD_CoV-2_/LY-CoV555 mAb complex. Color legend: gray, neutral mutations; light yellow, mildly destabilizing mutations; light red, destabilizing mutations; red, highly destabilizing mutations (see Table 1). (B) Main interactions involving the S-RBD_CoV-2_ E484K mutant at the interface with the LY-CoV555 (bamlanivimab) mAb as obtained from the relevant equilibrated MD simulation. Images for all other A/D/G/Q/R and V mutants are shown in Figure S1 (see also Table S5 for details). Colors and other explanations are the same as in Figure 3.

An utterly similar situation is observed in the presence of the A, G, R, and V mutants (see Figure S1 and Table S5 for details). In line with this, the predicted changes in binding free energy for the replacement of the wild-type E484 with A/G/K/R/V in the S-RBD_CoV-2_/LY-CoV555 relevant complexes (ΔΔG_CoV-2_(E484A) = −6.18 ± 0.10 kcal/mol, ΔΔG_CoV-2_(E484G) = −7.58± 0.18 kcal/mol, ΔΔG_CoV-2_(E484K) = −7.83 ± 0.11 kcal/mol, ΔΔG_CoV-2_(E484R) = −7.99 ± 0.12 kcal/mol, ΔΔG_CoV-2_(E484V) = −6.02 ± 0.14 kcal/mol, Figure 4A and Table S4) support the prominent contribution played by this residue in anchoring the viral protein/LY-CoV555 mAb binding interface and the LY-CoV555 escaping potential of the E484A, E484G, E484K, E484R, and E484V SARS-CoV-2 circulating mutants. The charged-to-neutral isosteric replacement E484Q has a moderately destabilizing effect (ΔΔG_CoV-2_(E484Q) = −2.53 ± 0.16 kcal/mol, Figure 4A, Table S4), since the two strong bifurcated SBs characterizing the wild-type complex are replaced with two single HBs, one between the side chain of Q484 and the side chain of arginine at position 50 of the mAb HC (2.96 ± 0.11 Å), and one between the backbone C=O group of Q484 and the side chain of arginine 96 of the Ab LC (2.70 ± 0.16 Å), respectively (Figure S1, Table S5). Finally, similarly to E484 the mutated D484 can establish SB interactions with the side chains of LY-CoV555 R_H_50 (2.94 ± 0.13 Å) and R_L_96 (2.81 ± 0.19 Å and 3.26 ± 0.22 Å), along with the full network of CIs seen in the wild-type complex, overall resulting in a predicted neutral effect on the related protein/protein interface (ΔΔG_CoV-2_(E484D) = −0.61 ± 0.13 kcal/mol, Figure 4A, Table S4, Figure S1 and Table S5).

#### Q493

According to relevant CAS-based prediction, Q493 also plays a primary stabilizing role at the S-protein/LY-CoV555 mAb interface (Figure 2B, Tables S1-S2). The analysis of the MD trajectories of all considered mutants (Q493H/K/L/R) reveals that, with respect to the wild-type Q493, all residues except H493 induce a strong destabilizing effect at the interface with the LY-CoV555 mAb (Figure 5A, Figure S2, Tables S4 and S6). With R493 as a proof-of-principle, Figure 5B shows that this mutant is no longer able to form the three fundamental HBs across the protein/protein interface with E_H_102, and R_H_104 on the Ab heavy chain, respectively (see also Table S6). Moreover, the spike Y449 no longer engages the side chains of the two LY-CoV555 mAb HC residues T_H_28 and S_H_31 in polar interactions, leaving the rest of the CI network substantially unchanged (Figures 3B and 5B, Table S6). Accordingly, the predicted affinity of this mutant viral protein for the LY-CoV55 mAb is markedly lower than that of the native counterpart (ΔΔG_CoV-2_(Q493R) = −4.57 ± 0.11 kcal/mol, Figure 5A, Table S4). Analogous effects are predicted for the other two mutants Q493K and Q493L, reflecting in a comparable decrease of protein/protein binding strength (ΔΔG_CoV-2_(Q493K) = −4.83 ± 0.12 kcal/mol and ΔΔG_CoV-2_(Q493L) = −4.26 ± 0.18 kcal/mol, respectively, Figure 5A, Table S4, Figure S2 and Table S6). These data therefore suggest that the Q493K/L/R mutants could all be LY-CoV555 escaping mutants. At variance with these, mutating Q493 into histidine introduce a somewhat less drastic changes in the topology of the viral protein-antibody interface. In particular, Q493H is still able to preserve one HB with the side chain of the LY-CoV555 R_H_104 (3.39 ± 0.15 Å) while the second HB interaction with the same mAb residue is replaced by a π/cation interaction (Figure S2, Table S6). Notably, the HB between H493 and E_H_102 is also missing in the entire MD trajectory of this S-RBD_CoV-2_ mutant/mAb complex, in addition to the polar CI between Y449 on spike and T_H_28 on LY-CoV555 (Figure S2, Table S6). In line with this, a moderate variation of the corresponding free energy of binding (Figure 5A, Table S4), that is, ΔΔG_CoV-2_(Q493H) = −1.95 ± 0.11 kcal/mol.

**Figure 5.**
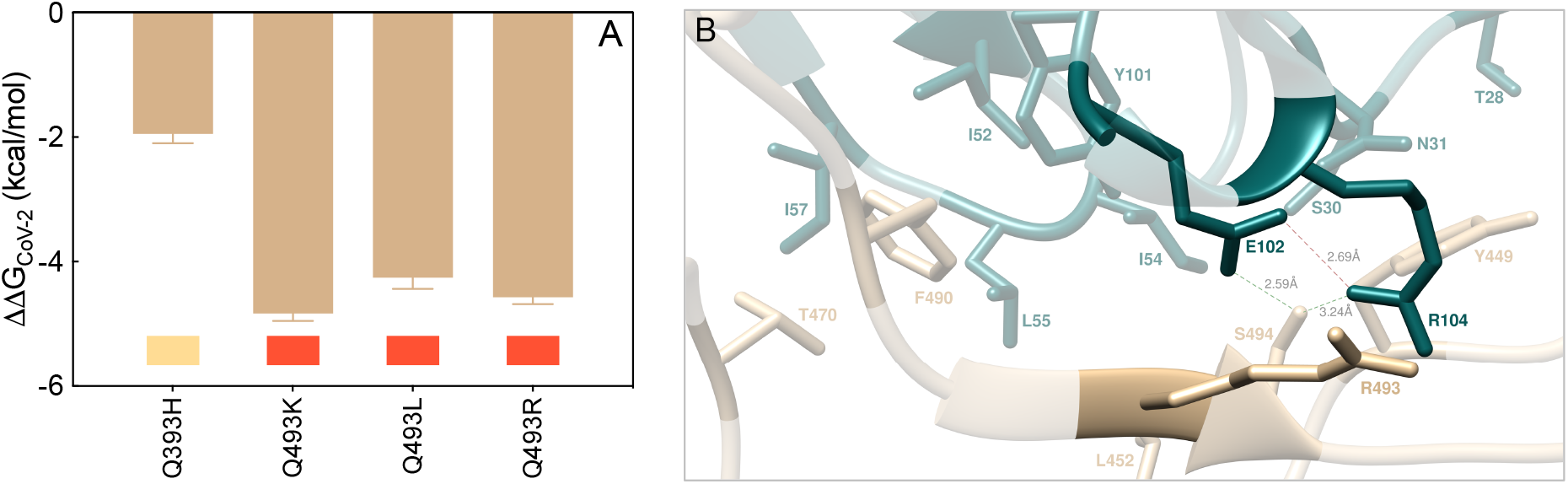
(A) Change in binding free energy (ΔΔG = ΔG_WILD-TYPE_ − ΔG_MUTANT_) predicted by computational mutagenesis of the S-RBD_CoV-2_ wild-type residue Q493 for the corresponding S-RBD_CoV-2_/LY-CoV555 Ab complex. Colors and other explanations as in Figure 4. The numerical values of ΔΔG, all related energy terms, and all underlying intermolecular intramolecular interactions are reported in Table S4, Figure S2, and Table S6. (B) Main interactions involving the S-RBD_CoV-2_ Q493R mutant at the interface with the LY-CoV555 (bamlanivimab) mAb as obtained from the relevant equilibrated MD simulation. Images for all other H/K and L mutants are shown in Figure S2 (see also Tables S6 for details). Colors and other explanations are the same as in Figure 3. New HBs and SBs eventually detected in each mutant complex are also indicated using dark green/red broken lines and black labels. Further important HBs and SBs detected in each complex are also indicated using light green/red broken lines and light gray labels.

#### S494

The third SARS-CoV-2 spike position highlighted by the CAS results as a key residue in binding the LY-CoV555 mAb is S494 (Figure 2B). Mutagenesis of this residue into A, P, R, and T reflects into strong interface destabilizing effects, exception made for the S494T substitution for which only a mild effect is observed (Figure 6A, Table S4). In the case of the R494 mutant, the current MD simulations show that both main intermolecular HBs in which the wild-type residue is involved (*i*.*e*., S494-E_H_102 and S494-R_H_104) are longer detected in the trajectory of the correspondent mutant viral protein/mAb complex (Figure 6B, Table S7). Also, two out of three further stabilizing HBs between the adjacent and important Q493 residue on S-RBD_CoV-2_ and the side chains of LY-CoV555 E_H_102 and R_H_104 are no longer formed in the presence of the R494 mutation (Figures 3B and 6B, Table S7). These evidences, along with several missing stabilizing CIs at the protein/protein interface (see Table S7 for details), concur to lower the predicted affinity of the R494 mutant S-RBD_CoV-2_ for the LY-CoV555 mAb (ΔΔG_CoV-2_(S494R) = −5.81 ± 0.17 kcal/mol, Figure 6A, Table S4). Accordingly, the three circulating mutants S494A, S494P, and S494R are all predicted to be potential LY-CoV555 escaping variants.

**Figure 6.**
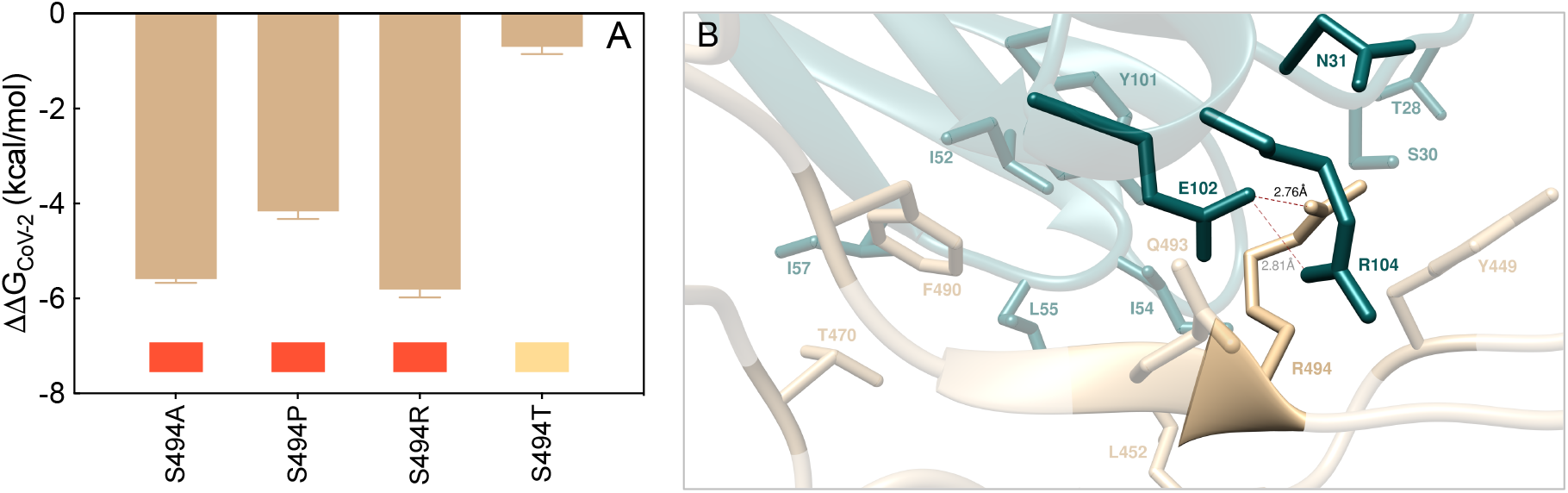
(A) Change in binding free energy (ΔΔG = ΔG_WILD-TYPE_ − ΔG_MUTANT_) predicted by computational mutagenesis of the S-RBD_CoV-2_ wild-type residue S494 for the corresponding S-RBD_CoV-2_/LY-CoV555 Ab complex. Colors and other explanations as in Figure 4. The numerical values of ΔΔG, all related energy terms, and all underlying intermolecular intramolecular interactions are reported in Table S4, Figure S3, and Table S7. (B) Main interactions involving the S-RBD_CoV-2_ S494R mutant at the interface with the LY-CoV555 (bamlanivimab) mAb as obtained from the relevant equilibrated MD simulation. Images for all other A/P and T mutants are shown in Figure S3 (see also Tables S7 for details). Colors and other explanations are the same as in Figure 3.

When S-RBD_CoV-2_ S494 is mutated into threonine (S494), the MD-predicted interaction network at the corresponding Ab binding interface is only moderately perturbed with respect to that described above for the wild-type complex; in particular, only the HBs between T494 on the SARS-CoV-2 RBD and E_H_102 on the HC of LY-CoV555, and between the viral Q493 and the same glutamic acid on the Ab HC are replaced by two polar CIs (Figure S3, Table S7). In line with this, the related value of ΔΔG_CoV-2_(S494T) is slightly unfavorable and equal to - 0.70 ± 0.15 kcal/mol (Figure 6A and Table S4).

#### V483, F486, and Y489

The mutagenesis results obtained by mutating these three viral spike amino acids into the reported variants (V483A/F/G/I/L, F486I/L/S, and Y489C/F/H/S, respectively) ultimately confirm the minor role played by these residues at the SARS-CoV-2 RBD/LY-CoV555 mAb binding interface (Figure 7A-C, Table S4).

**Figure 7.**
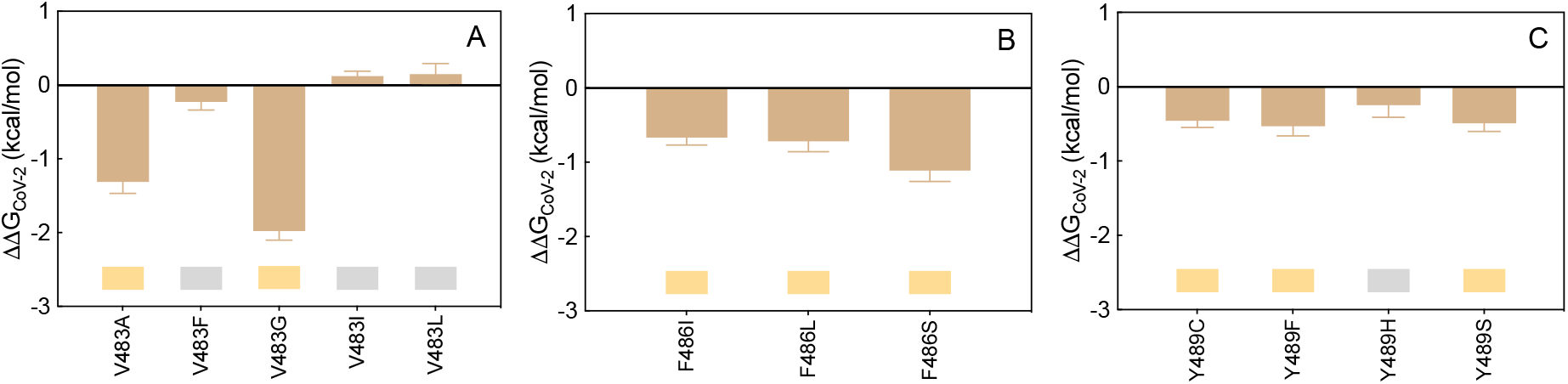
Change in binding free energy (ΔΔG = ΔG_WILD-TYPE_ − ΔG_MUTANT_) predicted by computational mutagenesis of the S-RBD_CoV-2_ wild-type residues V483 (A), F486 (B), and Y489 (C) for the corresponding S-RBD_CoV-2_/LY-CoV555 mAb complexes. Colors and other explanations as in Figure 4. The numerical values of ΔΔG, all related energy terms, and all underlying intermolecular intramolecular interactions are reported in Table S4, Figures S4-S6, and Tables S8-S10.

Indeed, the analysis of the respective MD trajectories reveals that each interaction network is practically conserved in all relevant supramolecular assemblies (see Figures S4-S6 and Tables S8-S10 for details). Accordingly, the SARS-CoV-2 spike mutations at residues 483, 486, and 489 reported so far in circulating viral populations are predicted to be tolerated at each respective position.

#### Y449, L452, T470 and F490

As it could be anticipated from the relevant CAS data discussed above, the *in silico* mutagenesis results for these further four viral protein residues into the reported variants (Y449D/F/H/N/S, L452M/Q/R, T470A/I/K/N and F490L/S/V/Y, respectively) also confirm a remarkable degree of tolerability to substitution at each of these spike positions in binding the LY-CoV555 Ab, with the remarkable exceptions of the L452R and – albeit to a lower extent – the F490S mutations (Figure 8A-D, Table S4, Figures S7-S10, Tables S11-S14).

**Figure 8.**
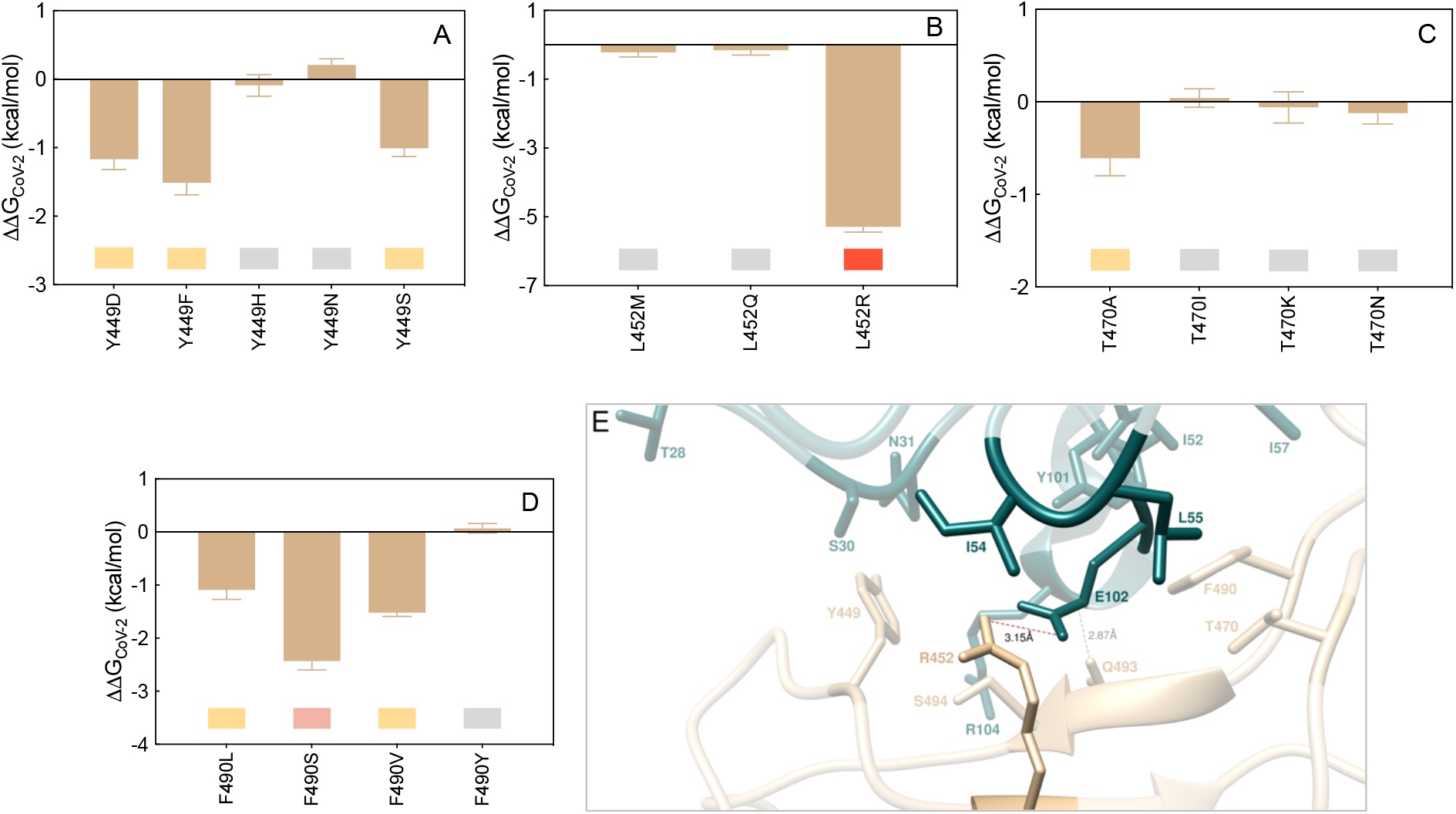
Change in binding free energy (ΔΔG = ΔG_WILD-TYPE_ − ΔG_MUTANT_) predicted by computational mutagenesis of the S-RBD_CoV-2_ wild-type residues Y449 (A), L452 (B), T470 (C) and F490 (D) for the corresponding S-RBD_CoV-2_/LY-CoV555 mAb complexes. Colors and other explanations as in Figure 4. The numerical values of ΔΔG, all related energy terms, and all underlying intermolecular intramolecular interactions are reported in Table S4, Figures S7-S10, and Tables S11-S14. (E) Main interactions involving the S-RBD_CoV-2_ L452 R mutant at the interface with the LY-CoV555 (bamlanivimab) mAb as obtained from the relevant equilibrated MD simulation. Images for the other M and Q mutants are shown in Figure S8 (see also Tables S12 for details). Colors and other explanations are the same as in Figure 3.

Together with T470 and F490, the spike residue L452 is a part of a hydrophobic region at the binding interface with LY-CoV555; accordingly, when this amino acid is replaced by the small non-polar methionine or even by the polar glutamine, the corresponding viral/antibody interface remains almost unaffected (Figures 8B and S8, Table S12). On the contrary, upon mutation of L452 into the positively charged and long-chained asparagine, a substantial modification of the relative binding region is observed in the corresponding MD trajectory, as shown in Figure 8E. Specifically, the structurally-important internal SB between the side chains of LY-CoV555 E_H_102 and R_H_104 on the Ab HC (Figure 3B) is no longer detected, as it is replaced by an analogous yet intermolecular interaction between the former mAb residue and the mutant spike R452 (3.15 ± 0.11 Å). Contextually, however, the formation of this new interface SB is accompanied by the loss of all other crucial intermolecular interactions involving both S-RBD_CoV-2_ key residues Q493 and S494 (Figure 8E, Table S12). This, in turn, properly reflects in the substantial variation of the corresponding binding free energy value, so that ΔΔG_CoV-2_(L452R) = −5.29 ± 0.15 kcal/mol, Figure 8B, Table S4), thereby supporting the LY-CoV55 escaping potential of this SARS-CoV-2 spike isoform.

### Computational Alanine Scanning of the SARS-CoV-2 Spike Protein Residues at the Binding Interface with the LY-CoV016 (Etesevimab) Monoclonal Antibody

Within the same distance and energetic cutoffs adopted for the analysis of the viral S-protein/LY-CoV555 mAb complex (4.0 Å and 0.5 kcal/mol, respectively), the inspection of the equilibrated MD trajectory of the alternative S-RBD_CoV-2_/LY-CoV016 mAb assembly reveals that 13 viral protein residues persistent contact 16 residues of the Fab portion of the LY-CoV016 Ab at the relative binding interface. Of the latter, 13 amino acids locate on the mAb HC and 3 belong to the mAb LC, respectively (Figure 9, Table S15).

**Figure 9.**
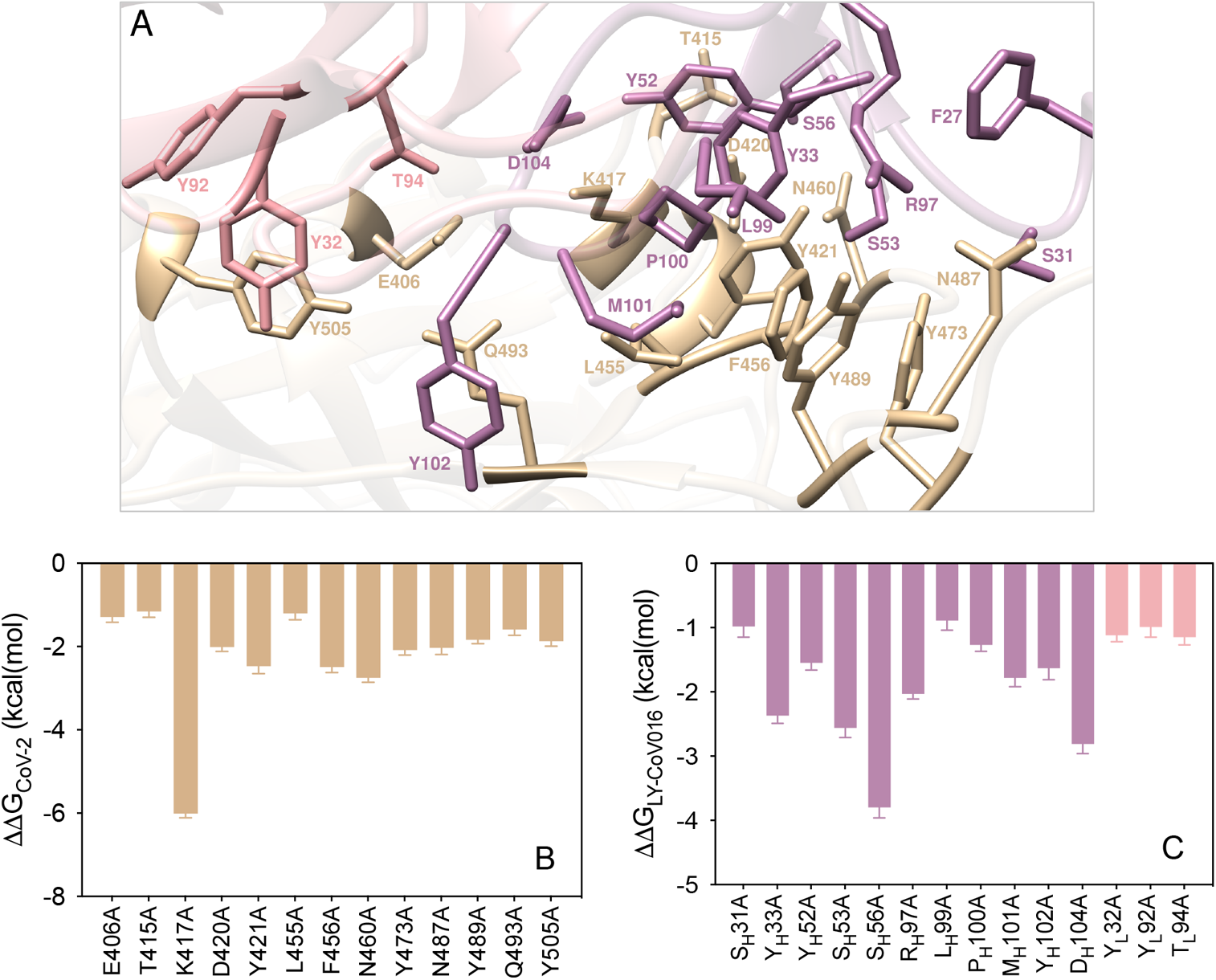
(A) Structural details of the binding interface between the LY-CoV016 (etesevimab) mAb and the viral spike protein receptor-binding domain of SARS-CoV-2 (S-RBD_CoV-2_). The secondary structures of LY-CoV555 and S-RBD_CoV-2_ are portrayed as light mulberry and light pink icing ribbons, respectively. Each interacting protein residue is highlighted in dark matching-colored sticks and labeled. Binding energy change (ΔΔG = ΔG_WILD-TYPE_ − ΔG_ALA_) obtained from the computational alanine-scanning (CAS) mutagenesis for the S-RBD_CoV-2_ residues at the binding interface with the LY-CoV016 mAb (B) and for the LY-CoV016 mAb residues at the binding interface with the viral protein RBD (C). Negative ΔΔG values indicate unfavorable substitution for alanine in the relevant position. For the numerical values of ΔΔG and all related energy terms, see the text and Tables S16-S17.

According to the CAS results shown in Figure 9B-C (see also Tables S16-S17), and at variance with the binding mode just discussed for the alternative mAb LY-Cov555, the viral RBD/LY-CoV016 binding interface is substantially more diffused and characterized by four distinct regions, the first of which locates in the area centered around the S-RBD_CoV-2_ residues K417 and N460 (Figure 10A).

**Figure 10.**
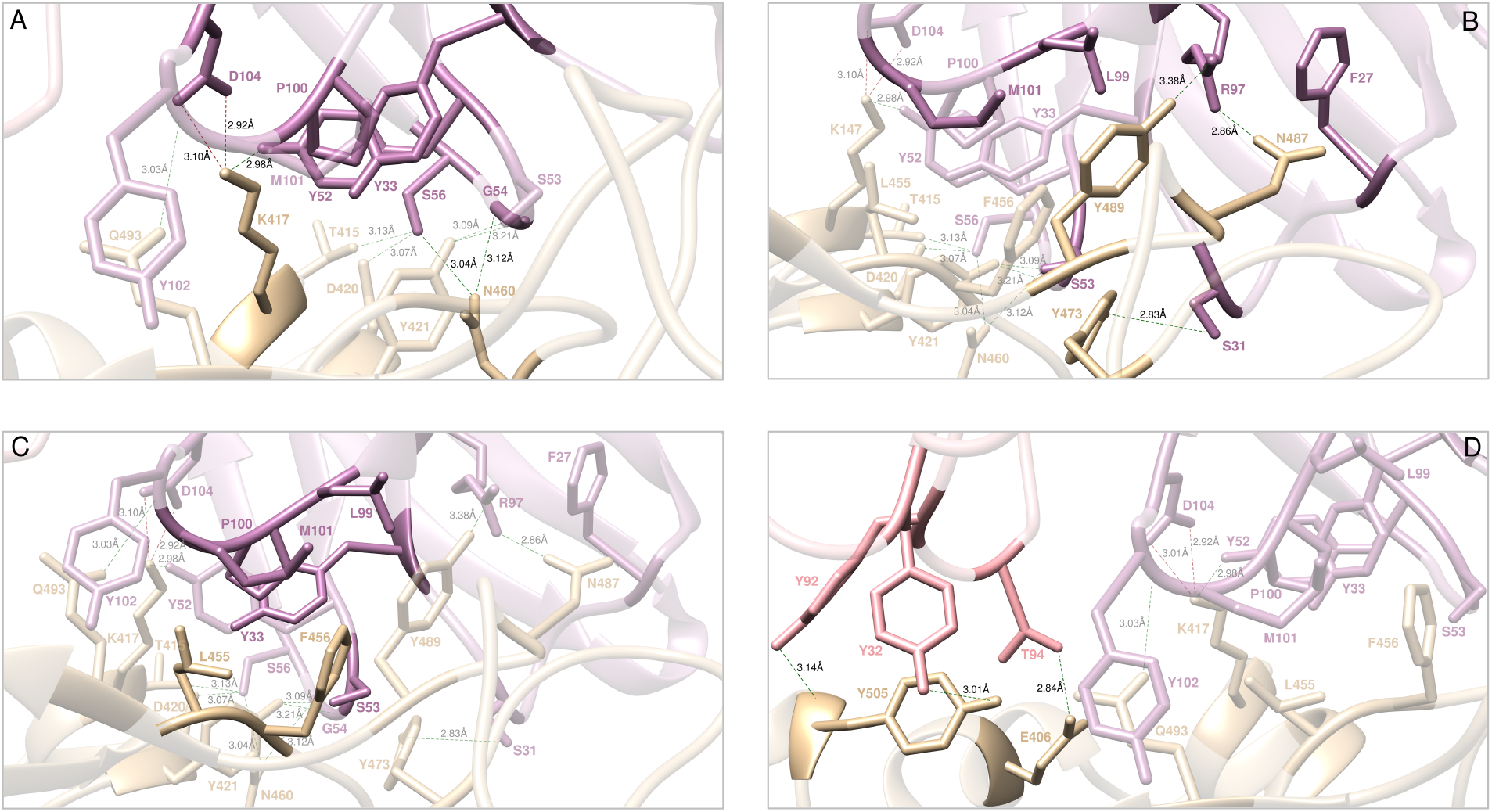
Main interactions involving the viral S-RBD_CoV-2_ residues K417 and N460 (A), Y473, N487, and Y489 (B), L455 and F456 (C), and E406 and Y505 at the interface with the LY-CoV016 (etesevimab) mAb as obtained from equilibrated MD simulations. In this and all remaining figures, the secondary structures of the S-RBD_CoV-2_ is shown as a light tan ribbon, while the HC and LC of the LY-CoV016 mAb are portrayed as light mulberry and light pink icing ribbons, respectively. Each S-RBD_CoV-2_ residue under discussion and all other residues directly interacting with it are highlighted in dark matching-colored sticks and labeled; further residues/interactions related to the residue under investigation are evidenced in light matching-colored sticks and labelled in light gray. Hydrogen bonds (HBs) and salt bridges (SBs) are represented as dark green and dark red broken lines, respectively, and the relevant average distances are reported accordingly. Further important HBs and SBs detected in each complex are also indicated using light green/red broken lines and light gray labels (see Table S15 for details).

Specifically, the MD trajectory of the S-RBD_CoV-2_/LY-CoV016 complex shows that K417 binds the side chain of D_H_104 on the mAb HC *via* a bifurcated SB (2.92 ± 0.15 Å and 3.10 ± 0.17 Å). Additionally, the K417 side chain is also involved in a stable HB with Y_H_52 (2.98 ± 0.14 Å) and in CI distance with the side chains of Y_H_33 (ΔΔG_LY-CoV016_(Y_H_33A) = −2.37 ± 0.12 kcal/mol) and P_H_100 (ΔΔG_LY-CoV016_(P_H_100A) = −1.27 ± 0.10 kcal/mol (see Figures 9C and 10A, and Tables S15 and S17). In agreement with this interaction pattern, the K417A mutation in CAS reduces the binding affinity of the corresponding S-RBD_CoV-2_ for the LY-CoV016 mAb by 6 kcal/mol (ΔΔG _CoV-2_(K417A) = −6.01 ± 0.10 kcal/mol, Figure 9B, Table S16). In the same context, the corresponding values of ΔΔG_LY-CoV016_(D_H_104A) = −2.81 ± 0.15 kcal/mol and ΔΔG_LY-CoV016_(Y_H_52A) = −1.55 ± 0.11 kcal/mol (Figure 9C, Table S17) properly rank the relative importance of these LY-Cov016 mAb residues at corresponding viral protein/antibody interface described above.

On the other hand, the S-RBD_CoV-2_ N460 residue is involved in two permanent HBs with the side chain of S_H_56 (3.04 ± 0.09 Å) and with the oxygen atom of the backbone of G_H_54 (3.12 ± 0.17 Å), and the relevant value of ΔΔG obtained by CAS for the N460A mutation (ΔΔG_CoV-2_(N460A) = −2.75 ± 0.11 kcal/mol, Figure 9B, Table S16) confirms the fundamental role of this spike residue at the binding interface. Interestingly, the change in binding free energy predicted by CAS for the S_H_56A substitution on the LY-Cov016 mAb (ΔΔG_LY-CoV016_(S_H_56A) = −3.80 ± 0.16 kcal/mol, Figure 9C, Table S17) accounts for the existence of additional stabilizing interactions in this region, characterized by a strong and virtuous network of HBs and CIs. In detail, S_H_56 is also involved in two stable HBs with the hydroxyl group of T415 (3.13 ± 0.10 Å, ΔΔG_CoV-2_(T415A) = −1.16 ± 0.14 kcal/mol) and the side chain of D420 (3.07 ± 0.15 Å, ΔΔG_CoV-2_(D420A) = −2.01 ± 0.11 kcal/mol), respectively (Figures 9B and 10A, and Tables S15-S16). Moreover, two additional S-RBD_CoV-2_ residues concur in determining the stability of the protein/protein interface (Figure 10A, Table S15): Y421 (ΔΔG_CoV-2_(Y421A) = −2.47 ± 0.18 kcal/mol) and Q493 (ΔΔG_CoV-2_(Q493A) = −1.59 ± 0.14 kcal/mol) (Figure 9B and Table S16). Specifically, the spike tyrosine 421 performs two HBs with LY-CoV016 S_H_53 (3.09 ± 0.11 Å, ΔΔG_LY-CoV016_(S_H_53A) = −2.56 ± 0.15 kcal/mol) and the nitrogen backbone atom of G_H_54 (3.21 ± 0.14 Å), respectively, while glutamine 493 is involved in the same type of intermolecular interaction with the amide moiety of the backbone of Y_H_102 (3.03 ± 0.18 Å), in addition to a favorable CI with the side chain of M_H_101 (Figures 9C and 10A, Table S15 and S17).

The second important region for the stabilization of the S-RBD_CoV-2_/LY-CoV016 complex is mainly composed by the viral residues Y473, N487 and Y489 (Figure 10B, Table S15) and the mAb HC residue R_H_97. Indeed, two HBs are detected between the guanidine moiety of R_H_97 (ΔΔG_LY-CoV016_(R_H_97A) = −2.03 ± 0.08 kcal/mol, Figure 9C, Table S17) and the side chain of N487 (2.86 ± 0.15 Å, ΔΔG_CoV-2_(N487A) = −2.03 ± 0.16 kcal/mol) and the hydroxyl group of Y489 (3.38 ± 0.13 Å, ΔΔG_CoV-2_(Y489A) = −1.84 ± 0.09 kcal/mol), respectively (Figures 9B and 10B, and Tables S15-S16). Additionally, the CIs of these viral protein amino acids with the LY-CoV016 HC residues F_H_27, L_H_99 (ΔΔG_LY-CoV016_(L_H_99A) = −0.89 ± 0.15 kcal/mol), and M_H_101 (ΔΔG_LY-CoV016_(M_H_101A) = −1.78 ± 0.14 kcal/mol) further contribute to binding interface stabilization (Figure 10B, Table S15, Figure 9C, Table S17). Moreover, the S-RBD_CoV-2_ Y473 (ΔΔG_CoV-2_(Y473A) = −2.08 ± 0.12 kcal/mol) is stably engaged in an HB with the side chain of S_H_31 (2.83 ± 0.21 Å, ΔΔG_LY-CoV016_(S_H_31A) = −0.98 ± 0.17 kcal/mol) and in a polar interaction with S_H_53 (Figure 9B-C, Figure10B, Tables S15-S17).

Located in between the two protein/protein interface regions just described, the third binding zone is identified by a network of van der Waals and hydrophobic interactions mainly involving the S-RBD_CoV-2_ residues L455 (ΔΔG_CoV-2_(L455A) = −1.20 ± 0.16 kcal/mol) and F456 (ΔΔG_CoV-2_(F456A) = −2.49 ± 0.13 kcal/mol) (see Figures 9B and 10C, Tables S1 and S16). In particular, this hydrophobic patch at the S-RBD_CoV-2_/LY-CoV016 mAb interface sees F456 as the pivot point that coordinates and to appropriately orient the mAb residues Y_H_33, S_H_53, L_H_99, P_H_100 and M_H_101 for further protein/protein interactions (Figure 10C, Table S15).

Finally, the last detected binding region – although apparently not a primary determinant of the viral/Ab interface stabilization – supports the optimization of the mutual protein/protein recognition. Indeed, this region involves only the LY-CoV016 mAb LC residues Y_L_32 (ΔΔG_LY-CoV016_(Y_L_32A) = −1.12 ± 0.10 kcal/mol), Y_L_92 (ΔΔG_LY-CoV016_(Y_L_92A) = −0.99 ± 0.16 kcal/mol), and T_L_94 (ΔΔG_LY-CoV016_(T_L_94A) = −1.15 ± 0.12 kcal/mol) in a set of stable HBs with the viral spike residues E406 (2.84 ± 0.27 Å, ΔΔG_CoV-2_(E406A) = −1.29 ± 0.13 kcal/mol), the hydroxyl group of Y505 (3.01 ± 0.19 Å, ΔΔG_CoV-2_(Y505A) = −1.87 ± 0.12 kcal/mol), and the nitrogen backbone atom of the same tyrosine (3.14 ± 0.13 Å) (Figures 9B-C and 10D, and Tables S15-S17).

### Mutagenesis of the SARS-CoV-2 spike protein at the interface with the LY-CoV016 antibody

The same data survey reported by Starr and coworkers^57^ led us to identify the following naturally occurring mutations at the SARS-CoV-2 S protein residues contacting the LY-CoV016 Ab: E406D/Q, T415A/I/N/P/S, K417E/N/R/T, D420A/G/N, L455F/S/V, F456L/Y, N460I/K/S/T, Y473F/H, N487D, Y489C/F/H/S, Q493H/K/L/R and Y505F/H/W. Below, we report and discuss different effects exerted by each of these spike mutant residues on the structure and strength of the resulting S-RBD_CoV-2_ /LY-CoV016 binding interface by adopting again the same color-coded criterion shown in Table 1.

#### K417

Our CAS data highlight the wild-type K417 as a hot-spot residue in the interaction between the S-RBD_CoV-2_ and the LY-CoV016 mAb (Figure 9B and 10A, Tables S15-S16). As such, is not surprising that replacing K417 on the viral protein with each of the alternative circulating mutants (K417E/N/R/T) reflects into a very strong interface disrupting behavior, with the exception of the substitution K417R, for which our *in silico* mutagenesis data anticipate a neutral effect (Figures 11 and S11, Tables S18-S19). As seen in Figure 11B for the K417N mutant as a paradigm, the current MD simulations show that both the double SB with the side chain of LY-CoV016 D_H_104 and the HB between the charged amine group of K417 and the hydroxyl moiety of Y_H_52 cannot longer be detected in the MD trajectory of the mutant complex. Also, the K417 network of underlying CIs involving a polar interaction with Y_H_33 and a hydrophobic contact with P_H_100 is likewise perturbed when K is replaced by N at the same position (Figure 11B, Table S19). Quite importantly, in the same binding region the presence of N417 affects other protein/protein interactions, including the absence of the three stabilizing HBs between N460 and G_H_54, T415 and S_H_56, and Y421 and S_H_53, respectively (Figure 11B, Table S19). These evidences ultimately translate into a drastically lower affinity of the N417 mutant spike protein for the LY-CoV016 mAb (ΔΔG_CoV-2_(K417N) = −7.27 ± 0.07 kcal/mol, Figure 11A and Table S18).

**Figure 11.**
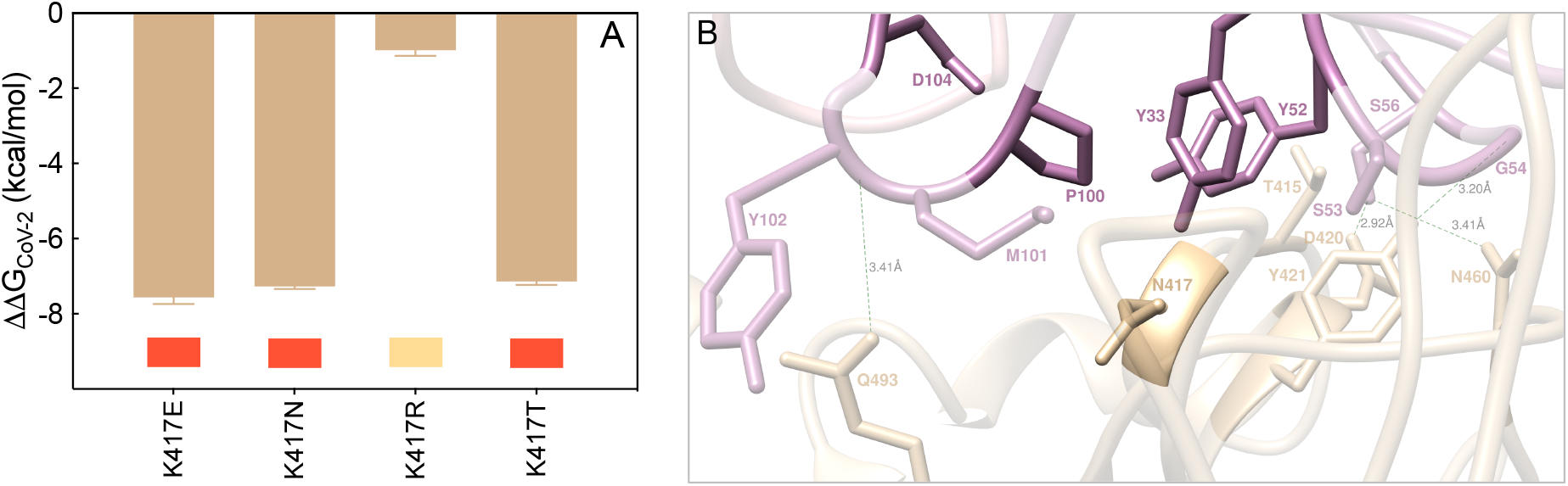
(A) Change in binding free energy (ΔΔG = ΔG_WILD-TYPE_ − ΔG_MUTANT_) predicted by computational mutagenesis of the S-RBD_CoV-2_ wild-type residue K417 for the corresponding S-RBD_CoV-2_/LY-CoV016 mAb complex. Negative ΔΔG values indicate unfavorable substitutions for the mutant residue in the relevant position. The numerical values of ΔΔG, all related energy terms, and all underlying intermolecular interactions are reported in Table S18, Figure S11, and Table S19. In this and all other similar figures, the colored boxes below each bar in the graphs show the classification of the destabilizing effects of the corresponding mutation on the S-RBD_CoV-2_/LY-CoV016 Ab complex. Color legend: gray, neutral mutations; light yellow, mildly destabilizing mutations; light red, destabilizing mutations; red, highly destabilizing mutations (see Table 1). (B) Main interactions involving the S-RBD_CoV-2_ K417N mutant at the interface with the LY-CoV016 (etesevimab) mAb as obtained from the relevant equilibrated MD simulation. Images for the K417E/R/T mutants are shown in Figure S11 (see also Table S19 for details). Colors and other explanations are the same as in Figure 10.

The effects observed for the E417 and T417 spike mutants are completely similar to those just described for the N417 isoform, although in both these cases the interface HB between the side chains of N460 and G_H_54 is again detected yet at the expenses of the analogous interaction between the viral Q493 and the mAb Y_H_102, which is missing along the entire MD trajectories of the corresponding supramolecular assemblies. As such, the variation in binding free energy between the wild-type and a mutant spike protein carrying either E or T at position 417 in complex with the LY-CoV016 mAb is predicted to be quite significant (ΔΔG_CoV-2_(K417E) = −7.56 ± 0.18 kcal/mol and ΔΔG_CoV-2_(K417T) = −7.14 ± 0.09 kcal/mol, Figure 11A, Table S18). On the other hand, only minor interface perturbations are observed in the presence of the R417 mutation (Figure S11, Table S19), in line with the predicted small change in protein/protein affinity (ΔΔG_CoV-2_(K417R) = −0.99 ± 0.15 kcal/mol, Figure 11A, Table S18).

#### D420 and N460

Converting the SARS-CoV-2 spike residues D420 and N460 in alanine *via* CAS analysis suggests that these two mutant isoforms induce only limited perturbing effects at the relative S-RBD_CoV-2_/LY-CoV016 mAb binding interface (Figures 9B and 10A, and Table S16). Surprisingly, however, the computational mutagenesis data for all circulating viral mutations at these two spike positions (D420A/G/N and N460I/K/S/T) reveal strong interface-destabilizing effects in all cases, with difference in free energy of binding with respect to the wild-type protein ranging from ∼ -5 to ∼ -3 kcal/mol (*i*.*e*., ΔΔG_CoV-2_(D420A) = −4.36 ± 0.17 kcal/mol, ΔΔG_CoV-2_(D420G) = −4.39 ± 0.10 kcal/mol, ΔΔG_CoV-2_(D420N) = −4.23 ± 0.10 kcal/mol, ΔΔG_CoV-2_(N460I) = −5.01 ± 0.14 kcal/mol, ΔΔG_CoV-2_(N460K) = −3.28 ± 0.18 kcal/mol, ΔΔG_CoV-2_(N460S) = −4.06 ± 0.09 kcal/mol, ΔΔG_CoV-2_(N460T) = −4.12 ± 0.14 kcal/mol) (see also Table S18).

The molecular rationale for these results relies not only on the fact all D420 and N460 S-RBD_CoV-2_ variants remove all direct interactions provided by aspartic acid (420) or glutamine (460) but also exert a domino effect on the nearby spike residues populating the same binding region, including T415 and, above all, the hot spot K417. Considering A420 as an exemplar of all D420 mutant behavior, from Figure 12B it is quickly seen that the wild-type HB with the side chain of S_H_56 is evidently missing, as are the three topical HBs between T415 and S_H_56, between K417 and Y_H_52, and between N460 and G_H_54, respectively (see also Figure S12 and Table S20). Similarly, taking I460 as a proof-of-concept for the N460 variants, the relevant MD trajectory reveals the absence of the same HBs engaged by the side chains of T145 and K147 on the viral spike and those of S_H_56 and Y_H_52 on the Ab, respectively. At the same time the direct wild-type 460 intermolecular HBs with G_H_54 and S_H_56 are obviously suppressed in the I460 Spike mutant/mAb complex, along with loss of the same interaction between Y421 and S_H_53 across the respective protein/protein interface (Figures 12D and S13, Table S21).

**Figure 12.**
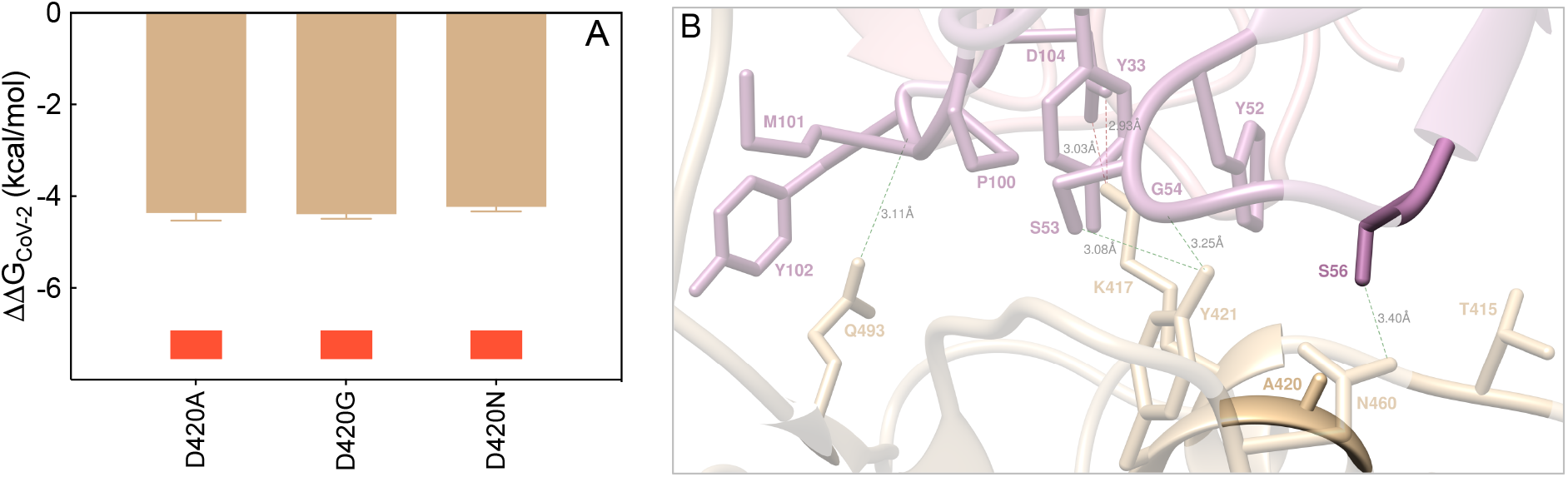
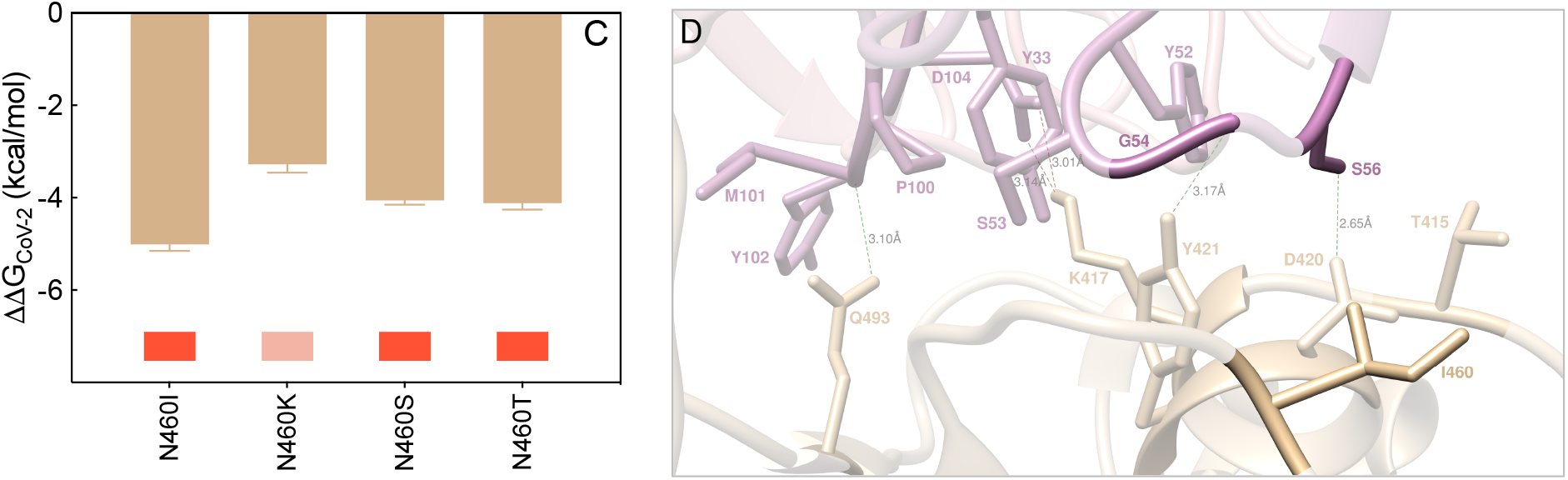
Change in binding free energy (ΔΔG = ΔG_WILD-TYPE_ − ΔG_MUTANT_) predicted by computational mutagenesis of the S-RBD_CoV-2_ wild-type residue D420 (A) and N460 (C) for the corresponding S-RBD_CoV-2_/LY-CoV016 mAb complex. Negative ΔΔG values indicate unfavorable substitutions for the mutant residue in the relevant position. The numerical values of ΔΔG, all related energy terms, and all underlying intermolecular interactions are reported in Table 18, Figures S12-S13, and Tables S20-S21. Main interactions involving the S-RBD_CoV-2_ D420A (B) and N460I (D) mutants at the interface with the LY-CoV016 (etesevimab) mAb as obtained from the relevant equilibrated MD simulations. Images for the D420G/N and N460K/S/T mutants are shown in Figures S12-S13 (see also Tables S20-S21 for details). Colors and other explanations are the same as in Figure 10.

#### T415 and Q493

Although these two SARS-CoV-2 S protein residues belong to the first binding region centered around two key viral amino acids in the stabilization of the S-RBD_CoV-2_/LY-CoV016 mAb interface – N460 and K417, respectively (Figure 10A) – the ΔΔG values currently predicted for replacement of both these spike positions with all reported variants (T415A/I/N/S and Q493H/K/L/R) indicate only moderate interface perturbation outcomes, with the notable deviation of the T415P mutant, for which a robust loss in affinity of this viral variant for the mAb is anticipated (Figures 13A-B and S14-S15, Tables S18 and S22-23).

**Figure 13.**
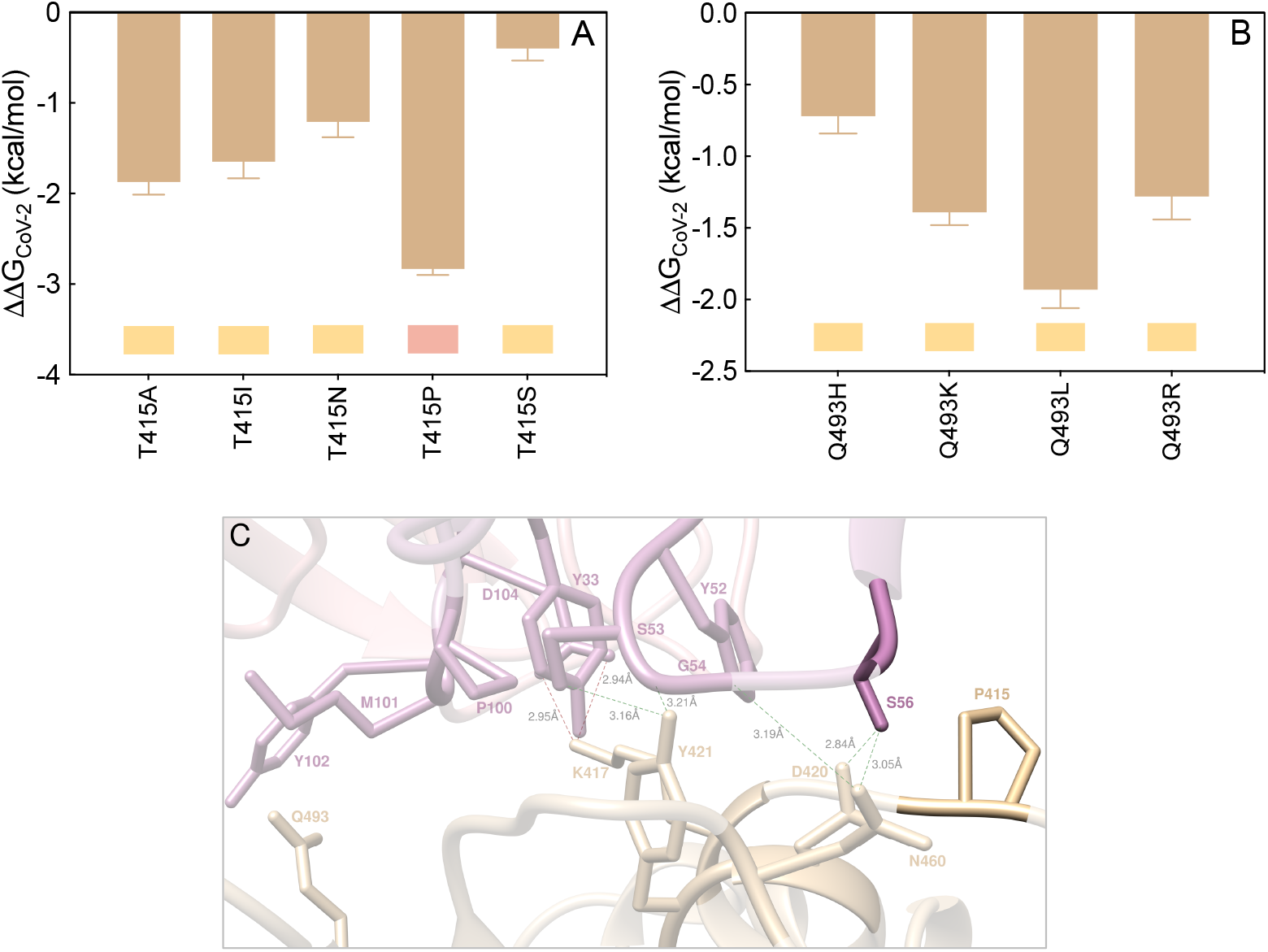
Change in binding free energy (ΔΔG = ΔG_WILD-TYPE_ − ΔG_MUTANT_) predicted by computational mutagenesis of the S-RBD_CoV-2_ wild-type residues T415 (A) and Q493 (B) for the corresponding S-RBD_CoV-2_/LY-CoV016 mAb complexes. Colors and other explanations as in Figure 4. The numerical values of ΔΔG, all related energy terms, and all underlying intermolecular intramolecular interactions are reported in Table S18, Figures S14-S15, and Tables S22-S23. (C) Main interactions involving the S-RBD_CoV-2_ T415P mutant at the interface with the LY-CoV016 (etesevimab) Ab as obtained from the relevant equilibrated MD simulation. Images for the T415A/I/N/S mutants and for all Q493 mutations are shown in Figures S14-S15 (see also Tables S22-S23 for details). Colors and other explanations are the same as in Figure 10.

In detail, while the conservative mutation T415S ensues the preservation of the wild-type interaction network, in the case of the T415A/I/N variants the analysis of the present simulations shows that the two spike-mAb anchoring intermolecular HBs in which the wild-type residue is involved (*i*.*e*., T415-S_H_56 and K417-Y_H_52, Figure 10A) cannot longer be detected in the trajectory of the mutant complexes. However, the extensive underlying network of other SBs, HBs, and CIs remains almost unaffected across the corresponding binding interfaces (Figure S14, Table S22), ultimately resulting in a limited decrement of the corresponding free energy variations (Figure 13A, Table S18). In the case of the T415P variant, the remarkably negative effect on spike/mAb affinity predicted by our *in silico* mutagenesis is sensibly linked – aside for the same perturbating effects just discussed for the other mutations at the same viral protein location – to the absence of the additional interface HB and CIs between the side chains of Q493 on the spike and of Y_H_102 on the mAb HC (Figure 13C, Table S22). Accordingly, the S-RBD_CoV-2_ T415P mutation reported so far in circulating viral populations is predicted to be potentially destabilizing for the S-RBD_CoV-2_/LY-CoV016 interface (ΔΔG_CoV-2_(T415P) = −2.83 ± 0.07 kcal/mol, Figure 13A and Table S18).

#### Y473, N487 and Y489

These viral residues belong to the spike/LY-Cov016 binding region that, according to the relevant MD trajectories, is characterized by an important network of stabilizing hydrogen bonds. Nevertheless, the present computational mutagenesis data report only neutral-to-mild interface destabilizing effects for the circulating SARS-CoV-2 RBD variants of Y473 (Y473F/H) and N487 (N487D) (see Table S18, Figures S16-S17, and Tables S24-S25 for details). Briefly, in the case of the Y473F mutation the loss of the HB between the wild-type tyrosine and the side chain of S_H_31 on the LY-CoV016 mAb HC (Figure 10B) detected in the MD trajectories of all variants has only minor effects on all other important intermolecular interactions populating same region, while the phenylalanine-to-histidine mutation is virtually conservative (ΔΔG_CoV-2_(Y473F) = −1.67 ± 0.08 kcal/mol and ΔΔG_CoV-2_(Y473H) = −0.19 ± 0.16 kcal/mol, respectively, Table S18, Figure S16, Table S24). The predicted minor loss in binding affinity of the D487 spike variant for the LY-CoV016 mAb (ΔΔG_CoV-2_(N487D) = −0.70 ± 0.09 kcal/mol, Table S18), on the other hand, is the result of a compensatory effect as the mutant aspartic acid provides a permanent intermolecular SB with the guanidine group of the Ab R_H_97 that makes up for the loss of the two HBs between Y473 and S_H_31 and Y489 and R_H_97, respectively (Figures 10B and S17, Table S25).

Finally, according to our MD analysis the circulating Y489 S-RBD_CoV-2_ variants induce a moderate decrease in affinity of the viral spike protein for the LY-CoV016 mAb (Figure 14A, Table S18). In particular, the conversion of tyrosine 489 into cysteine or serine results in the abrogation of the direct HB with R_H_97 as well as the hydrophobic contact with L_H_99. Moreover, the HB involving Y473 and S_H_31 is also missing along the entire MD trajectories of the Y489C and Y489S S-RBD_CoV-2_ mutant proteins, as shown in Figure 14B for the S489 isoform (see also Figure S18 and Table S26).

**Figure 14.**
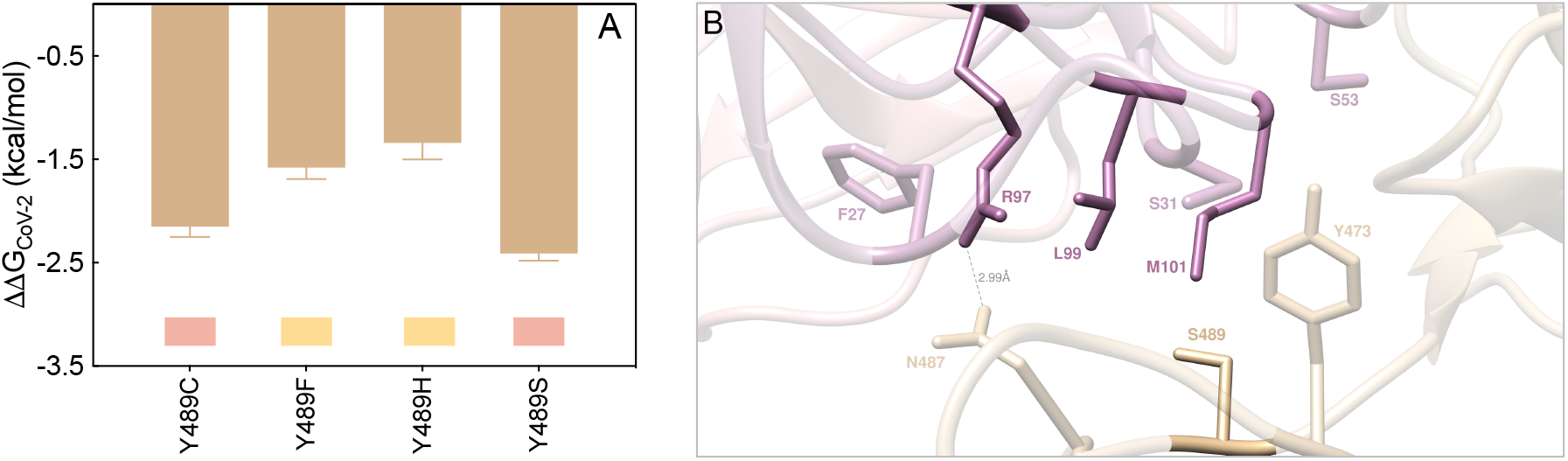
(A) Change in binding free energy (ΔΔG = ΔG_WILD-TYPE_ − ΔG_MUTANT_) predicted by computational mutagenesis of the S-RBD_CoV-2_ wild-type residue Y489 for the corresponding S-RBD_CoV-2_/LY-CoV016 mAb complex. Colors and other explanations as in Figure 4. The numerical values of ΔΔG, all related energy terms, and all underlying intermolecular intramolecular interactions are reported in Table S18, Figure S18 and Table S26. (B) Main interactions involving the S-RBD_CoV-2_ Y489S mutant at the interface with the LY-CoV016 (etesevimab) mAb as obtained from the relevant equilibrated MD simulation. Images for the Y489C/F/H mutants are shown in Figure S18 (see also Table S26 for details). Colors and other explanations are the same as in Figure 10.

In line with this, the calculated ΔΔG values numerically support moderate interface destabilizing effects upon substitution of the wild-type tyrosine with these two residues (ΔΔG_CoV-2_(Y489C) = −2.15 ± 0.10 kcal/mol, and ΔΔG_CoV-2_(Y489S) = −2.41 ± 0.07 kcal/mol, respectively, Figure 14A and Table S18).

#### E406, L455, F456 and Y505

The actual computational data for mutating these four viral protein residues into the SARS-CoV-2 circulating variants (E406D/Q, L455F/S/V, F456L/Y and Y505F/H/W, respectively) account for neutral-to-mildly negative effects on the stability of the corresponding S-RBD_CoV-2_/LY-CoV016 mAb binding interface, with estimated ΔΔG values all below 1 kcal/mol for all alternative amino acids considered (Figure 15, see also Table S18, Figures S19-S22 and Tables S27-S30).

**Figure 15.**
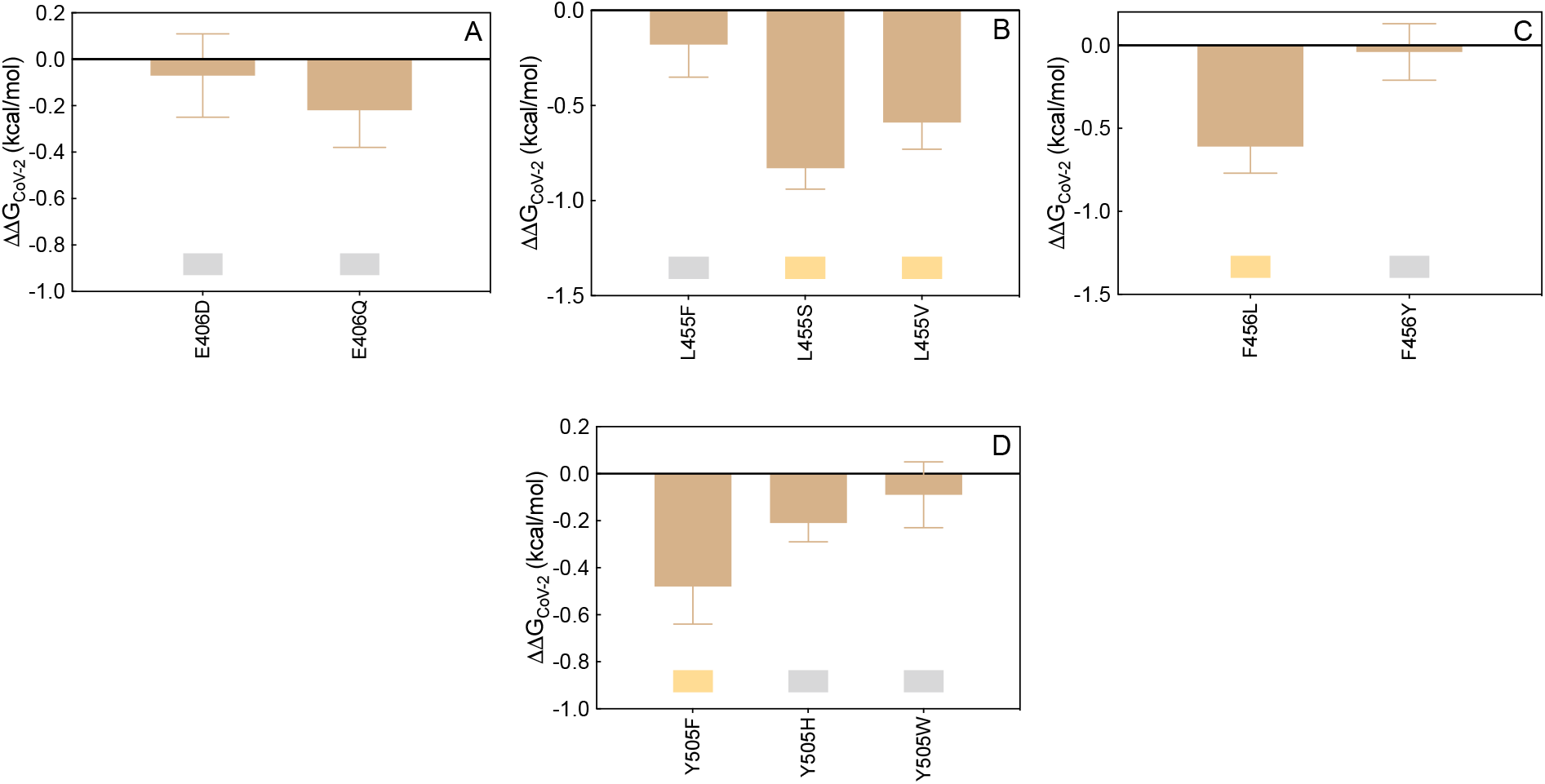
Change in binding free energy (ΔΔG = ΔG_WILD-TYPE_ − ΔG_MUTANT_) predicted by computational mutagenesis of the S-RBD_CoV-2_ wild-type residues E406 (A), L455 (B), F456 (C) and Y505 (D) for the corresponding S-RBD_CoV-2_/LY-CoV016 mAb complexes. Colors and other explanations as in Figure 4. The numerical values of ΔΔG, all related energy terms, and all underlying intermolecular intramolecular interactions are reported in Table S18, Figures S19-S22 and Tables S27-S30.

Therefore, all these SARS-CoV-2 spike position variants do not appear to have a significant role in escaping the LY-CoV016 antibody.

## CONCLUSIONS

The purpose of this work was to provide an *in silico* molecular rationale of the role eventually played by currently circulating S-RBD_CoV-2_ mutations in evading the immune surveillance effects elicited by the two Eli Lilly LY-CoV555/bamlanivimab and LY-CoV016/etesevimab monoclonal antibodies. Table 2 summarizes the main findings from this study and shows that, compared to the wild-type SARS-CoV-2 spike protein, all mutants highlighted in light or dark red are predicted to be markedly more resistant to neutralization by both these mAbs, those shown in yellow might exert only mildly perturbing protein/protein binding, while those listed in gray are not likely to confer any mAb escaping advantage to the viral protein.

**Table 2.** Color-code ranking of circulating SARS-CoV-2 spike protein mutants with respect to their predicted resistance to neutralization by LY-CoV-555 (bamlanivimab) and LY-CoV016 (etesevimab) monoclonal antibodies. Colors as in Table 1.

| S-RBD<sub>CoV-2</sub> wild-type position | LY-CoV555 | S-RBD<sub>CoV-2</sub> wild-type position | LY-CoV016 |
| --- | --- | --- | --- |
| E484 | 484A/D/G/K/Q/R/V | K417 | 417E/N/R/T |
| Q493 | 493H/K/L/R | D420 | 420A/G/N |
| S494 | 494A/P/R/T | N460 | 460I/K/S/T |
| V483 | 483A/F/G/I/L | T415 | 415A/I/N/P/S |
| F486 | 486I/L/S | Q493 | 493H/K/L/R |
| Y489 | 489C/F/H/S | Y473 | 473F/H |
| Y449 | 449D/F/H/N/S | N487 | 487D |
| L452 | 452M/Q/R | Y489 | 489C/F/H/S |
| T470 | 470A/I/K/N | E406 | 406D/Q |
| F490 | 490L/S/V/Y | L455 | 455F/S/V |
|  |  | F456 | 456L/Y |
|  |  | Y505 | 505F/H/W |

According to the most updated version (March 18, 2021) of the” Fact sheet for health care providers – emergency use authorization (EUA) of bamlanivimab and etesevimab”, ^59^ resistant variants to both mAbs were already reported by Eli-Lilly researchers using S-protein directed evolution and serial passages in cell cultures of SARS-CoV-2 in the presence of either antibody. On the other hand, resistant variants were not reported when the two mAbs were tested together using the same methodology. Spike variants identified in these studies that presented reduced susceptibility to the LY-CoV555 mAb included the following substitutions: E484D/K/Q, F490S, Q493R, and S494P. Concerning the spike position 484, after our CAS approach identified E484 as a key player residue at the S-RBD_CoV-2_/LY-CoV555 binding interface (Figures 2A and 3A, Table S2), we considered all possible mutations actually reported at this position in circulating viral variants (*i*.*e*., E484A/D/G/K/Q/R/V), and found that all these amino acid variations should confer strong escaping ability to bamlanivimab (Figures 4, Table 2). From a validation standpoint, the E484K mutation is present in a large number of VOC/VOI/VUM - including the lineages B.1.525 (firstly reported in Nigeria on 12/20), P.1 and P.2 (Brazil, 12/20), P.3 (The Philippines, 01/21), B.1.351 (South Africa, 09/20), B.1.621 (Colombia, 01/2021), and some strains of lineages B.1.1.7 (firstly reported in the United Kingdom on 09/20) and B.1.526 (reported on 11/20 in the city of New York, USA)^51, 52^ - and it is indeed well known to confer substantial loss of sensitivity to neutralizing Abs found in sera of convalescent and vaccinated individuals.^60 61-71^ Further, for all these variants there is evidence of a significant reduction in neutralization by the LY-CoV555/LY-CoV016 and other mAb treatments.^57, 60, 72-75^ Collier and coworkers very recently reported that the introduction of the E484K mutation in the B.1.1.7 background (to account for the new VOC B.1.1.7+E484K found in the virus isolated both in UK and in Pennsylvania, USA)^76^ led to robust loss of neutralizing activity by 19 out of 31 vaccine-elicited antibodies and mAbs if compared with the decrease in sensitivity conferred by the mutations in B.1.1.7 alone.^77^ Moreover, the E484Q/V/A/G/D mutations have been just described by Chen *et al*. as critical in promoting escape not only from Eli Lilly mAbs but also from other similar therapeutics that are currently in clinical trials.^78^

Mutating the wild-type spike F490 into alanine also flagged this position as a residue affording an important contribution to the protein/protein interface (Figure 2A, Table S2). Interestingly, the corresponding mutagenesis into all reported variants (F490L/S/V/Y) revealed that only the F490S spike mutant is a potential escapee for LY-CoV555 (Figures 8, Table 2), in agreement with Lilly’s and other experimental observations.^59, 66, 78^ Of note F490S, although listed in the actual spike circulating mutations, is not a component of any VOC or VOI listed so far.^51, 52^ Finally, CAS predicted viral spike residues Q493 and S494 to be the two remaining hot spots at the viral protein/bamlanivimab binding interface (Figures 2A and 3B, Table S2). *In silico* mutagenesis of Q493 and S494 into the circulating variants (Q493H/K/L/R, and S494A/P/R/T) not only confirms Lilly’s data about Q493R and S494P as resistant mutations for LY-CoV555^59^ but also predicts a potential role of other substitutions at these two S-protein positions (*i*.*e*., Q493K/L and S494A/P/R) in mediating evasion to this mAb (Figures 5-6, Table 2). In line with these predictions, three new studies highlighted all these mutants as viral proteins that may hinder the efficiency of existing vaccines and expand in response to the increasing after-infection or vaccine-induced seroprevalence.^66, 78, 79^ Remarkably, the spike S494P mutation is a component of the B.1.17+S494P VOC^51^/VUM^52^ identified in United Kingdom in January 2021.

In the fact sheet produced by Lilly^59^ the spike 452 position was not mentioned as a possible site of LY-CoV555 escaping mutant *per se*. However, L452R is a spike mutation of interest (MOI)^52^ present in the VOC lineages B.1.427/B.1.429 (reported in California, USA, on 09/20), B.1.526.1 (New York City, USA, 10/20), and in the B1.617.1/B.1.617.2/B.1.617.3 lineages now rapidly and deadly spreading in India (12/20-02/21), where it is always found along with the D614G substitution. Importantly, the L452R mutation is also present in tandem with E484Q, in particular in the B.1.617.1 variant that is responsible for actual disease outbreaks in 49 countries in all six WHO regions.^80^ Using a pseudo-virus expressing the spike protein from the B.1.427/B.1.429 lineages, or the L452R substitution only, however, the researchers at Lilly reported reduced susceptibility to bamlanivimab and etesevimab together of 7.7-fold or 7.4-fold, respectively.^59^ Further experimental works^81-84^ already reported increased viral load/transmissibility and escape ability from neutralizing antibodies for this variant when tested against vaccine-elicited sera. Actually, in their preprint work Hoffmann *et al*. analyzed whether the SARS-CoV-2 VOC B.1.617 is more adept in entering cells and/or evade Ab responses.^85^ They found that B.1617 entered two out of 8 cell lines tested (specifically, the human lung- and intestine-derived Calu-3 and Caco-2 cell lines, respectively) with slightly increased efficiency, and was blocked by an entry inhibitor. However, in stark contrast, B.1.167 was found to be fully resistant to LY-CoV555 and partially resistant against neutralization by Abs elicited upon infection or vaccination with the Comirnaty/Pfizer-BioNTech vaccine. Our present data support the escaping potential of the L452R viral mutation with respect to bamlanivimab (Figure 8, panels B and E, Table 2), while we did not detect any effect in terms of changed affinity of this mutant protein toward etesevimab. Moreover, our data also suggest that the co-presence of the E484Q (Figure 5, Table 2) may synergistically contribute in rendering the two B.1.617.1 and B.1.617.3 variants potent evaders of antibody surveillance.

Concerning the alternative LY-CoV016 mAb, the official Lilly’s fact sheet^59^ reports that SARS-CoV-2 spike mutants showing reduced susceptibility to etesevimab include substitutions K417N, D420N, and N460K/S/T. In agreement with this and other evidences,^72, 79^ our current computational alanine/mutagenesis study marks K417 and all its reported variants (K417E/N/R/T) as the strongest hot spots in eliciting potential escape to the LY-CoV016 mAb (Figure 11A, Table 2). Of note, the K417N and K417T in particular are spike MOIs in the SARS-CoV2 VOC lineages B.1.351 and P.1, respectively. Similarly, not only the D420N but all reported circulating spike mutations at positions 420 are predicted by our study to be endowed with high LY-CoV016 escaping potential (Figure 12A, Table 2), in line with recent findings.^79^ Finally, and in full agreement with Lilly’s data,^59^ LY-CoV016 is also found to be escaped by all spike N460 variations (N460I/K/S/T) (Figure 12C, Table 2).

In addition, in the current study we identify three further single amino acid changes along the primary sequence of SARS-CoV-2 spike protein that – although not reported as current VOC/VOI/VUM – could escape the action of LY-CoV016, that is the T415P and the Y489C/S mutations (Figures 13A and 14A, Table2). Since these spike mutants are present in circulating viral variants, in our opinion they should be taken into consideration as they might limit the therapeutic usefulness of this mAb, both *per se* and in its cocktail combination with LY-CoV555.

As a conclusive remark concerning available anti-SARS-CoV-2 vaccines, according to the report by Andreano and Rappuoli published on May 10, 2021 in Nature Medicine^86^ the efficacy of the FDA/EMA approved Ad26.COV2-S vaccine (now Janssen COVID-19 Vaccine) and the EMA approved Oxford–AstraZeneca ChAdOx1 (now Vaxzevria) against the variant B.1.351 (South Africa, with E484K, K417N and N501Y as spike MOIs) decreased from 85% to 57% and from 62% to 10%, respectively. In parallel, the titer neutralizing antibodies induced by the m-RNA vaccines approved by both governmental agencies (*i*.*e*., the BNT162b2 Pfizer/BioNTech COVID-19 vaccine/Comirnaty and COVID-19 vaccine Moderna) against the same SARS-CoV-2 variant is reported to decline by 7- to 12 -fold, while no negative effect on neutralization is seen for the B.1.1.7 variant (with N501Y/D614G as spike MOIs). Additionally, the work of Planas and collaborators documented low titers of neutralizing antibodies against the B.1.351 variant in a cohort of 19 individuals after both doses of the Comirnaty vaccine.^87^ In all these cases, the spike E484K mutation appears to be the real key player in reducing neutralization by antibodies induced by the vaccines. And this, in turn, support the view that vaccination elicits a natural infection–like antibody response, and that spike variants like E484K may spread as antigenic evolutions of SARS-CoV-2 to efficiently evade this response. On the bright side, all vaccines currently approved appear at least to protect from the severe forms of infection,^88, 89^ and second-generation vaccines and mAbs aiming at containing VOC spreading are under investigation.^90^

In concluding this work, we report that a challenge of our global *in silico* results against the relevant experimental data just published by the Starr group^57^ resulted in an overall 90% agreement. This achievement not only constitutes a further, robust validation of our computer-based approach but also yields a molecular-based rationale for all relative experimental findings, and leads us to conclude that the current circulating SARS-CoV-2 and all possible emergent variants carrying these mutations in the spike protein can present new challenges for mAb-based therapies and ultimately threaten the fully-protective efficacy of currently available vaccines.

## Supporting information

Supporting information_part1

Supporting information_part2

Supporting information_part3

