## Supporting information_part1 for "*In Silico* Molecular-Based Rationale for SARS-CoV-2 Spike Circulating Mutations Able to Escape Bamlanivimab and Etesevimab Monoclonal Antibodies"

**Table S1.** Main intermolecular and intramolecular interactions between residues at the protein-protein interface detected during MD simulations of the LY-CoV555 antibody in complex with the RBDs of SARS-CoV-2 S-RBD (COV2). HB = hydrogen bond; SB = salt bridge; CI = contact interactions, including van der Waals/hydrophobic (vdW/h), polar (p), $\pi/\pi$ and $\pi$/cation ($\pi$/c) interactions. In the HB column, s-s indicates side chain-side chain interactions while s-b or b-s indicate side chain-backbone or backbone- side chain interactions, respectively.

| **HB** | **COV2** | **Length (Å)** | **LY-CoV555** |
|---|---|---|---|
| s-s | Q493 | $3.31 \pm 0.12$ | $R_H$104 |
| s-b | | $3.34 \pm 0.17$ | $R_H$104 |
| s-s | | $3.25 \pm 0.18$ | $E_H$102 |
| s-s | S494 | $3.18 \pm 0.19$ | $R_H$104 |
| s-s | | $2.86 \pm 0.16$ | $E_H$102 |
| **SB** | **COV2** | **Length (Å)** | **LY-CoV555** |
| | E484 | $2.74 \pm 0.10$ | $R_H$50 |
| | | $3.05 \pm 0.14$ | |
| | E484 | $2.82 \pm 0.11$ | $R_L$96 |
| | | $2.96 \pm 0.13$ | |
| **SB** | **LY-CoV555** | **Length (Å)** | **LY-CoV555** |
| s-s | $E_H$102 | $2.76 \pm 0.11$ | $R_H$104 |
| **CI** | **COV2** | **LY-CoV555** | |
| p | Y449 | $T_H$28 | |
| p | | $S_H$30 | |
| p | | $N_H$31 | |
| vdW/h | | $I_H$54 | |
| vdW/h | L452 | $I_H$54 | |
| vdW/h | | $L_H$55 | |
| vdW/h | T470 | $L_H$55 | |
| vdW/h | | $I_H$57 | |
| vdW/h | V483 | $W_H$47 | |
| vdW/h | | $R_H$50 | |
| vdW/h | | $N_H$59 | |
| vdW/h | | $T_L$94 | |
| vdW/h | | $R_L$96 | |
| vdW/h | E484 | $Y_H$101 | |
| vdW/h | | $Y_H$110 | |
| $\pi/\pi$ | F486 | $Y_L$32 | |
| $\pi/\pi$ | | $Y_L$92 | |
| vdW/h | Y489 | $Y_H$110 | |
| p | | Y'32 | |

| | | |
|---|---|---|
| vdW/h | | $I_H52$ |
| vdW/h | F490 | $L_H55$ |
| vdW/h | | $I_H57$ |
| π/π | | $Y_H101$ |
| p | S494 | $N_H31$ |

**Table S2.** Relative binding free energy and its components calculated by the combined computational alanine scanning mutagenesis – interaction entropy approach for the S-RBD of SARS-CoV-2(COV2) residues effectively involved in the binding interface with the LY-CoV55 antibody (see the SI Materials and Methods section for details). IE = interaction entropy. $\Delta\Delta G = \Delta G_{WILDTYPE} - \Delta G_{ALA}$ (see text for details).

| | **Y449A** | **L452A** | **T470A** | **V483A** | **E484A** | **F486A** | **Y489A** | **F490A** | **Q493A** | **S494A** |
|---|---|---|---|---|---|---|---|---|---|---|
| $\Delta\Delta E_{DISP}$ | -0.50 | -0.76 | -0.42 | -1.67 | -0.63 | -0.76 | -0.54 | -2.49 | -0.46 | -0.21 |
| $\Delta\Delta E_{ELE}$ | -1.22 | 0.09 | -0.19 | 0.09 | -5.57 | -0.77 | -0.49 | -0.07 | -3.93 | -3.66 |
| $\Delta\Delta H$ | -1.72 | -0.67 | -0.61 | -1.58 | -6.20 | -1.53 | -1.03 | -2.56 | -4.39 | -3.87 |
| $\Delta\Delta IE$ | -0.21 | -0.09 | -0.03 | -0.12 | 0.28 | 0.09 | -0.09 | 0.18 | 0.21 | -0.15 |
| **$\Delta\Delta G_{COV2}$** | **-1.93 (0.16)** | **-0.76 (0.11)** | **-0.64 (0.15)** | **-1.70 (0.18)** | **-5.92 (0.12)** | **-1.44 (0.15)** | **-1.12 (0.09)** | **-2.38 (0.22)** | **-4.18 (0.14)** | **-4.02 (0.12)** |

**Table S3.** Relative binding free energy and its components calculated by the combined computational alanine scanning mutagenesis – interaction entropy approach for LY-CoV555 antibody residues effectively involved in the binding interface with the S-RBD of SARS-CoV-2 (COV2) (see the SI Materials and Methods section for details). IE = interaction entropy. $\Delta\Delta G = \Delta G_{WILDTYPE} - \Delta G_{ALA}$ (see text for details).

| | **$N_H$31A** | **$R_H$50A** | **$L_H$55A** | **$Y_H$101A** | **$E_H$102A** | **$R_H$104A** | **$Y_H$110A** | **$Y_L$32A** | **$Y_L$92A** | **$R_L$96A** |
|---|---|---|---|---|---|---|---|---|---|---|
| $\Delta\Delta E_{DISP}$ | 0.12 | -0.22 | -1.15 | -0.56 | -1.35 | -1.27 | -0.79 | -0.24 | -0.32 | -0.48 |
| $\Delta\Delta E_{ELE}$ | -1.57 | -3.01 | 0.07 | -0.89 | -3.19 | -3.56 | 0.06 | -1.14 | -0.46 | -2.29 |
| $\Delta\Delta H$ | -1.45 | -3.23 | -1.08 | -1.45 | -4.54 | -4.83 | -0.73 | -1.38 | -0.78 | -2.77 |
| $\Delta\Delta IE$ | -0.11 | 0.21 | -0.02 | 0.11 | 0.22 | 0.25 | 0.12 | -0.16 | 0.05 | 0.18 |
| **$\Delta\Delta G_{COV2}$** | **-1.56 (0.14)** | **-3.02 (0.19)** | **-1.10 (0.08)** | **-1.34 (0.10)** | **-4.32 (0.13)** | **-4.58 (0.15)** | **-0.61 (0.09)** | **-1.54 (0.12)** | **-0.73 (0.17)** | **-2.59 (0.11)** |

**Table S4.** Relative binding free energy and its components for the mutated S-RBD of SARS-CoV-2 (COV2) residues in the binding interface with the LY-CoV555 antibody. IE = interaction entropy. IE = interaction entropy. $\Delta\Delta G = \Delta G_{WILDTYPE} - \Delta G_{MUTANT}$ (see text for details).

| | **Y449D** | **Y449F** | **Y449H** | **Y449N** | **Y449S** | **L452M** | **L452Q** | **L452R** | **T470A** | **T470I** | **T470K** | **T470N** |
|---|---|---|---|---|---|---|---|---|---|---|---|---|
| $\Delta\Delta E_{DISP}$ | -0.52 | -0.34 | -0.22 | -0.19 | -0.75 | -0.17 | -0.10 | -1.13 | -0.42 | 0.21 | 0.18 | 0.09 |
| $\Delta\Delta E_{ELE}$ | -0.54 | -1.13 | 0.03 | 0.32 | -0.31 | 0.04 | 0.15 | -4.28 | -0.27 | -0.12 | -0.35 | -0.20 |
| $\Delta\Delta H$ | -1.06 | -1.47 | -0.19 | 0.13 | -1.06 | -0.13 | 0.05 | -5.41 | -0.69 | 0.09 | -0.17 | -0.11 |
| $\Delta\Delta IE$ | -0.11 | -0.04 | 0.10 | 0.08 | 0.05 | -0.09 | -0.21 | 0.12 | 0.08 | -0.05 | 0.11 | -0.01 |
| **$\Delta\Delta G_{COV2}$** | **-1.17** | **-1.51** | **-0.09** | **0.21** | **-1.01** | **-0.22** | **-0.16** | **-5.29** | **-0.61** | **0.04** | **-0.06** | **-0.12** |
| | **(0.15)** | **(0.18)** | **(0.16)** | **(0.09)** | **(0.12)** | **(0.13)** | **(0.14)** | **(0.15)** | **(0.19)** | **(0.10)** | **(0.17)** | **(0.12)** |

| | **V483A** | **V483F** | **V483G** | **V483I** | **V483L** | **E484A** | **E484D** | **E484G** | **E484K** | **E484Q** | **E484R** | **E484V** |
|---|---|---|---|---|---|---|---|---|---|---|---|---|
| $\Delta\Delta E_{DISP}$ | -1.62 | 0.07 | -0.34 | 0.41 | 0.43 | -2.20 | -0.23 | -2.49 | -2.01 | -0.14 | -2.12 | -1.96 |
| $\Delta\Delta E_{ELE}$ | 0.21 | -0.34 | -1.37 | -0.16 | -0.12 | -3.86 | -0.20 | -4.92 | -5.75 | -2.41 | -5.99 | -4.03 |
| $\Delta\Delta H$ | -1.41 | -0.27 | -1.71 | 0.25 | 0.31 | -6.06 | -0.43 | -7.41 | -7.76 | -2.55 | -8.11 | -5.99 |
| $\Delta\Delta IE$ | 0.05 | 0.04 | -0.27 | -0.13 | -0.16 | -0.16 | -0.18 | -0.17 | -0.07 | 0.02 | 0.12 | -0.03 |
| **$\Delta\Delta G_{COV2}$** | **-1.36** | **-0.23** | **-1.98** | **0.12** | **0.15** | **-6.18** | **-0.61** | **-7.58** | **-7.83** | **-2.53** | **-7.99** | **-6.02** |
| | **(0.16)** | **(0.11)** | **(0.12)** | **(0.07)** | **(0.14)** | **(0.10)** | **(0.13)** | **(0.18)** | **(0.11)** | **(0.16)** | **(0.12)** | **(0.14)** |

| | **F486I** | **F486L** | **F486S** | **Y489C** | **Y489F** | **Y489H** | **Y489S** | **F490L** | **F490S** | **F490V** | **F490Y** | **Q493H** |
|---|---|---|---|---|---|---|---|---|---|---|---|---|
| $\Delta\Delta E_{DISP}$ | -0.52 | -0.74 | -1.55 | -0.05 | -0.08 | -0.09 | -0.24 | -0.97 | -1.52 | -1.38 | -0.20 | -0.78 |
| $\Delta\Delta E_{ELE}$ | 0.05 | -0.10 | 0.30 | -0.31 | -0.54 | -0.05 | -0.34 | 0.06 | -0.93 | -0.17 | 0.28 | -1.33 |
| $\Delta\Delta H$ | -0.47 | -0.84 | -1.25 | -0.36 | -0.62 | -0.14 | -0.58 | -0.91 | -2.45 | -1.55 | 0.08 | -2.11 |
| $\Delta\Delta IE$ | 0.16 | 0.12 | 0.14 | -0.10 | 0.04 | -0.11 | 0.09 | -0.18 | 0.02 | 0.03 | -0.01 | 0.16 |
| **$\Delta\Delta G_{COV2}$** | **-0.67** | **-0.72** | **-1.11** | **-0.46** | **-0.53** | **-0.25** | **-0.49** | **-1.09** | **-2.43** | **-1.52** | **0.07** | **-1.95** |
| | **(0.10)** | **(0.14)** | **(0.15)** | **(0.09)** | **(0.13)** | **(0.16)** | **(0.11)** | **(0.18)** | **(0.17)** | **(0.07)** | **(0.09)** | **(0.11)** |

| | **Q493K** | **Q493L** | **Q493R** | **S494A** | **S494P** | **S494R** | **S494T** |
|---|---|---|---|---|---|---|---|
| $\Delta\Delta E_{DISP}$ | -1.25 | -0.78 | -1.00 | -1.52 | -0.87 | -0.91 | -0.26 |
| $\Delta\Delta E_{ELE}$ | -3.49 | -3.61 | -3.38 | -4.17 | -3.06 | -5.09 | -0.45 |
| $\Delta\Delta H$ | -4.74 | -4.39 | -4.38 | -5.69 | -3.93 | -6.00 | -0.71 |
| $\Delta\Delta IE$ | -0.09 | 0.13 | -0.19 | 0.10 | -0.23 | 0.19 | 0.01 |
| **$\Delta\Delta G_{COV2}$** | **-4.83** | **-4.26** | **-4.57** | **-5.59** | **-4.16** | **-5.81** | **-0.70** |
| | **(0.12)** | **(0.18)** | **(0.11)** | **(0.08)** | **(0.16)** | **(0.17)** | **(0.15)** |

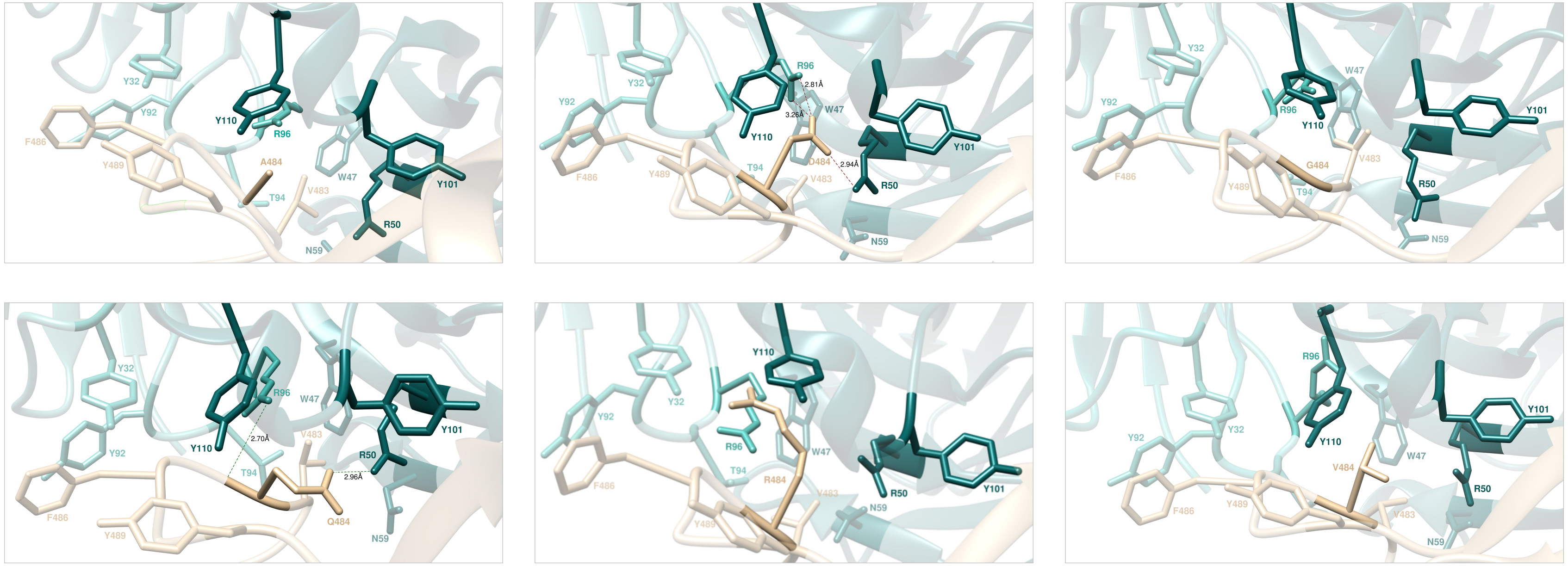


**Figure S1.** Main interactions involving the S-RBD$_{CoV-2}$ A484 (top left), D484 (top middle), G484 (top right), Q484 (bottom left), R484 (bottom middle), and V484 (bottom right) at the interface with the LY-CoV555 antibody as obtained from the corresponding equilibrated MD simulations. The wild-type E484 and the K484 mutant complexes are presented and discussed in the main text (Figures 3A and 4B). In this and all remaining Figures, the secondary structure of the S-RBD$_{CoV-2}$ is shown as a light tan ribbon, while the heavy and light chains of the LY-CoV555 antibody are portrayed as light teal and light Tiffany green ribbons, respectively. Each protein residue under discussion and all other residues directly interacting with it are highlighted in dark matching-colored sticks and labelled; further residues/interactions related to the residue under investigation are evidenced in light matching-colored sticks and labelled in light gray. Hydrogen bonds and salt bridges directly involving the residue under discussion are represented as dark green and dark red broken lines, respectively, and the relevant average distances are reported (in black) accordingly. New HBs and SBs eventually detected in each mutant complex are also indicated using dark green/red broken lines and black labels. Further important HBs and SBs detected in each complex are also indicated using light green/red broken lines and light gray labels. For further details see Tables S1 and S5.

**Table S5.** Main interactions between the wild-type S-RBD$_{CoV-2}$ residue E484 and all considered mutants* at the protein-protein interface detected during MD simulations of the RBD of SARS-CoV-2 (COV2) in complex with the LY-CoV555 antibody. HB = hydrogen bond; SB = salt bridge; CI = contact interactions, including van der Waals/hydrophobic (vdW/h), polar (p), $\pi/\pi$ and $\pi$/cation ($\pi$/c) interactions. In the HB column, s-s indicates side chain-side chain interactions while s-b or b-s indicate side chain-backbone or backbone-side chain interactions, respectively. Preserved/new or lost interactions are marked with the symbols ✓ and ✗, respectively. Relevant changes in the type/nature of the interactions are indicated in parenthesis. For HBs and SBs, the relevant average lengths (in Å) are also reported (their standard deviations, all within 10%, are not shown for clarity). *Mutant K484 is discussed in detail in main text.

| SB | COV2 | LY-CoV555 | E484 | A484 | D484 | G484 | K484 | Q484 | R484 | V484 |
|---|---|---|---|---|---|---|---|---|---|---|

| | | | | | | | | | | |
|---|---|---|---|---|---|---|---|---|---|---|
| s-s | X484 | $R_H$50 | ✓(2.74,3.05) | ✗ | ✓(2.94) | ✗ | ✗ | ✓(HB,2.96) | ✗ | ✗ |
| s-s | X484 | $R_L$96 | ✓(2.82,2.96) | ✗ | ✓(2.81,3.26) | ✗ | ✗ | ✓(HB,s-b,2.70) | ✗ | ✗ |
| **CI** | **COV2** | **LY-CoV555** | **E484** | **A484** | **D484** | **G484** | **K484** | **Q484** | **R484** | |
| vdW/h | X484 | $Y_H$101 | ✓ | ✓ | ✓ | ✗ | ✓ | ✓ | ✗ | ✓ |
| vdW/h | X484 | $Y_H$110 | ✓ | ✗ | ✓ | ✗ | ✓(p) | ✓ | ✓(π/c) | ✓ |
| **CI** | **COV2** | **LY-CoV555** | **E484** | **A484** | **D484** | **G484** | **K484** | **Q484** | **R484** | **V484** |
| vdW/h | V483 | $W_H$47 | ✓ | ✓ | ✓ | ✓ | ✓ | ✓ | ✗ | ✓ |
| vdW/h | V483 | $R_H$50 | ✓ | ✓ | ✓ | ✓ | ✓ | ✓ | ✓ | ✓ |
| vdW/h | V483 | $N_H$59 | ✓ | ✓ | ✓ | ✓ | ✓ | ✓ | ✓ | ✓ |
| vdW/h | V483 | $T_L$94 | ✓ | ✓ | ✓ | ✓ | ✓ | ✓ | ✓ | ✓ |
| vdW/h | V483 | $R_L$96 | ✓ | ✓ | ✓ | ✓ | ✓ | ✓ | ✓ | ✗ |
| π/π | F486 | $Y_L$32 | ✓ | ✓ | ✓ | ✓ | ✓ | ✓ | ✗(vdW/h) | ✓ |
| π/π | F486 | $Y_L$92 | ✓ | ✓ | ✓ | ✓ | ✓ | ✓ | ✗(vdW/h) | ✓ |
| vdW/h | Y489 | $Y_H$110 | ✓ | ✓ | ✓ | ✓ | ✗ | ✓ | ✓ | ✓ |
| p | Y489 | $Y_L$32 | ✓ | ✓ | ✗ | ✗ | ✗ | ✗ | ✗ | ✓ |

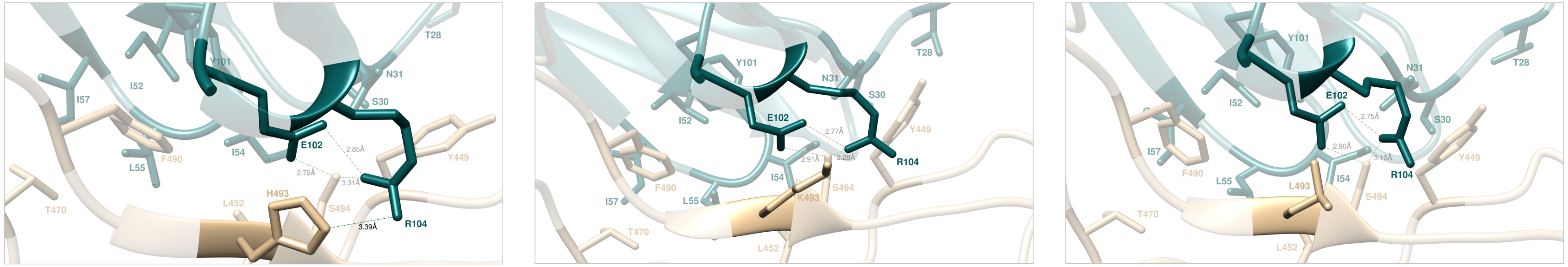


**Figure S2.** Main interactions involving the S-RBD$_{CoV-2}$ H493 (left), K493 (middle), and L493 (right) at the interface with the LY-CoV555 antibody as obtained from the corresponding equilibrated MD simulations. The wild type Q493 and the R493 mutant complexes are presented and discussed in the main text (Figures 3B and 5B). Colors and other explanations as in Figure S1. For further details see Tables S1 and S6.

**Table S6.** Main interactions between the wild-type S-RBD$_{CoV-2}$ residue Q493 and all considered mutants* at the protein-protein interface detected during MD simulations of RBD of SARS-CoV-2 (COV2) in complex with the LY-CoV555 antibody. Acronyms and other explanations as in Table S5. *Mutant R493 is discussed in detail in main text.

| **HB** | **COV2** | **LY-CoV555** | **Q493** | **H493** | **K493** | **L493** | **R493** |
|---|---|---|---|---|---|---|---|
| s-s | X493 | $E_H$102 | ✓(3.25) | ✗ | ✗ | ✗(vdW/h) | ✗(vdW/h) |
| s-s | X493 | $R_H$104 | ✓(3.31) | ✓(3.39) | ✗(vdW/h) | ✗(vdW/h) | ✗(vdW/h) |
| s-b | X493 | $R_H$104 | ✓(3.34) | ✗(π/c) | ✗ | ✗ | ✗ |
| **HB** | **COV2** | **LY-CoV555** | **Q493** | **H493** | **K493** | **L493** | **R493** |
| s-s | S494 | $E_H$102 | ✓(2.86) | ✓(2.79) | ✓(2.91) | ✓(2.90) | ✓(2.59) |
| s-s | S494 | $R_H$104 | ✓(3.18) | ✓(3.31) | ✓(3.28) | ✓(3.13) | ✓(3.24) |
| **CI** | **COV2** | **LY-CoV555** | **Q493** | **H493** | **K493** | **L493** | **R493** |
| p | Y449 | $T_H$28 | ✓ | ✗ | ✗ | ✗ | ✗ |
| p | Y449 | $S_H$30 | ✓ | ✓ | ✗ | ✗ | ✗(vdW/h) |
| p | Y449 | $N_H$31 | ✓ | ✓ | ✓ | ✓ | ✓ |
| vdW/h | Y449 | $I_H$54 | ✓ | ✓ | ✓ | ✓ | ✓ |
| vdW/h | L452 | $I_H$54 | ✓ | ✓ | ✓ | ✓ | ✓ |
| vdW/h | L452 | $L_H$55 | ✓ | ✓ | ✓ | ✓ | ✓ |
| vdW/h | T470 | $L_H$55 | ✓ | ✓ | ✓ | ✓ | ✓ |

| | | | | | | | |
|---|---|---|---|---|---|---|---|
| vdW/h | T470 | $I_H$57 | ✓ | ✓ | ✓ | ✓ | ✓ |
| vdW/h | F490 | $I_H$52 | ✓ | ✓ | ✓ | ✓ | ✓ |
| vdW/h | F490 | $L_H$55 | ✓ | ✓ | ✓ | ✓ | ✓ |
| vdW/h | F490 | $I_H$57 | ✓ | ✓ | ✓ | ✓ | ✓ |
| π/π | F490 | $Y_H$101 | ✓ | ✓ | ✓ | ✓ | ✓ |
| p | S494 | $N_H$31 | ✓ | ✓ | ✓ | ✓ | ✓ |
| **SB** | **LY-CoV555** | **LY-CoV555** | **Q493** | **H493** | **K493** | **L493** | **R493** |
| s-s | $E_H$102 | $R_H$104 | ✓(2.76) | ✓(2.85) | ✓(2.77) | ✓(2.75) | ✓(2.69) |

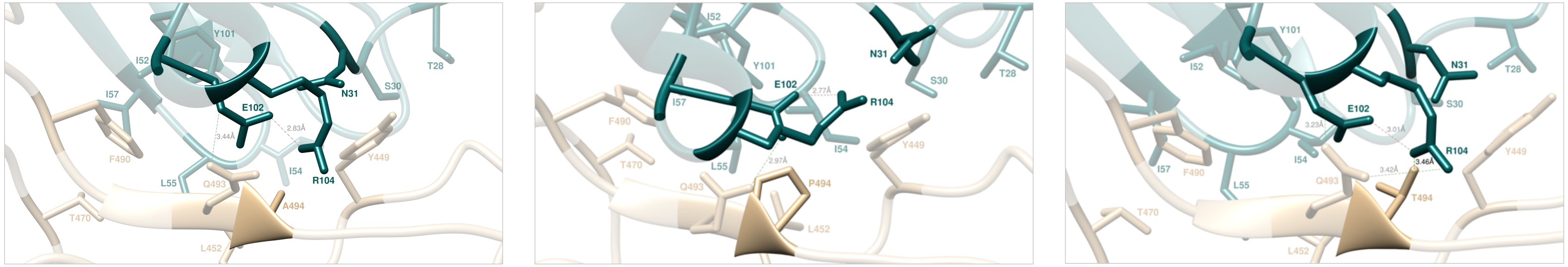


**Figure S3.** Main interactions involving the S-RBD$_{CoV-2}$ A494 (left), P494 (middle), and T494 (right) at the interface with the LY-CoV555 antibody as obtained from the corresponding equilibrated MD simulations. The wild type S494 and the R494 mutant complexes are presented and discussed in the main text (Figures 3B and 6B). Colors and other explanations as in Figure S1. For further details see Tables S1 and S7.

**Table S7.** Main interactions between the wild-type S-RBD$_{CoV-2}$ residue S494 and all considered mutants* at the protein-protein interface detected during MD simulations of RBD of SARS-CoV-2 (COV2) in complex with the LY-CoV555 antibody. Acronyms and other explanations as in Table S5. *Mutant R494 is discussed in detail in main text.

| **HB** | **COV2** | **LY-CoV555** | **S494** | **A494** | **P494** | **R494** | **T494** |
|---|---|---|---|---|---|---|---|
| s-s | X494 | $E_H$102 | ✓(2.86) | ✗ | ✗ | ✓(SB,2.76) | ✗(p) |
| s-s | X494 | $R_H$104 | ✓(3.18) | ✗ | ✗ | ✗ | ✓(3.46) |
| **CI** | **COV2** | **LY-CoV555** | **S494** | **A494** | **P494** | **R494** | **T494** |
| p | X494 | $N_H$31 | ✓ | ✗ | ✗ | ✓ | ✓ |
| **HB** | **COV2** | **LY-CoV555** | **S494** | **A494** | **P494** | **R494** | **T494** |
| s-s | Q493 | $E_H$102 | ✓(3.25) | ✗(p) | ✓(2.97) | ✗(p) | ✗(p) |
| s-s | Q493 | $R_H$104 | ✓(3.31) | ✗(p) | ✗ | ✗(p) | ✓(3.42) |
| s-b | Q493 | $R_H$104 | ✓(3.34) | ✓(3.44) | ✗ | ✗ | ✓(3.23) |
| **CI** | **COV2** | **LY-CoV555** | **S494** | **A494** | **P494** | **R494** | **T494** |
| p | Y449 | $T_H$28 | ✓ | ✗ | ✗ | ✗ | ✓ |
| p | Y449 | $S_H$30 | ✓ | ✗ | ✓ | ✗ | ✓ |
| p | Y449 | $N_H$31 | ✓ | ✓ | ✓ | ✗ | ✓ |
| vdW/h | Y449 | $I_H$54 | ✓ | ✓ | ✓ | ✗ | ✓ |
| vdW/h | L452 | $I_H$54 | ✓ | ✓ | ✓ | ✓ | ✓ |

| | | | | | | | |
|---|---|---|---|---|---|---|---|
| vdW/h | L452 | $L_H55$ | ✓ | ✓ | ✓ | ✓ | ✓ |
| vdW/h | T470 | $L_H55$ | ✓ | ✓ | ✓ | ✓ | ✓ |
| vdW/h | T470 | $I_H57$ | ✓ | ✓ | ✓ | ✓ | ✓ |
| vdW/h | F490 | $I_H52$ | ✓ | ✓ | ✓ | ✓ | ✓ |
| vdW/h | F490 | $L_H55$ | ✓ | ✓ | ✓ | ✓ | ✓ |
| vdW/h | F490 | $I_H57$ | ✓ | ✓ | ✓ | ✓ | ✓ |
| π-π | F490 | $Y_H101$ | ✓ | ✓ | ✓ | ✓ | ✓ |
| **SB** | **LY-CoV555** | **LY-CoV555** | **S494** | **A494** | **P494** | **R494** | **T494** |
| s-s | $E_H102$ | $R_H104$ | ✓(2.76) | ✓(2.83) | ✓(2.77) | ✓(2.81) | ✓(3.01) |

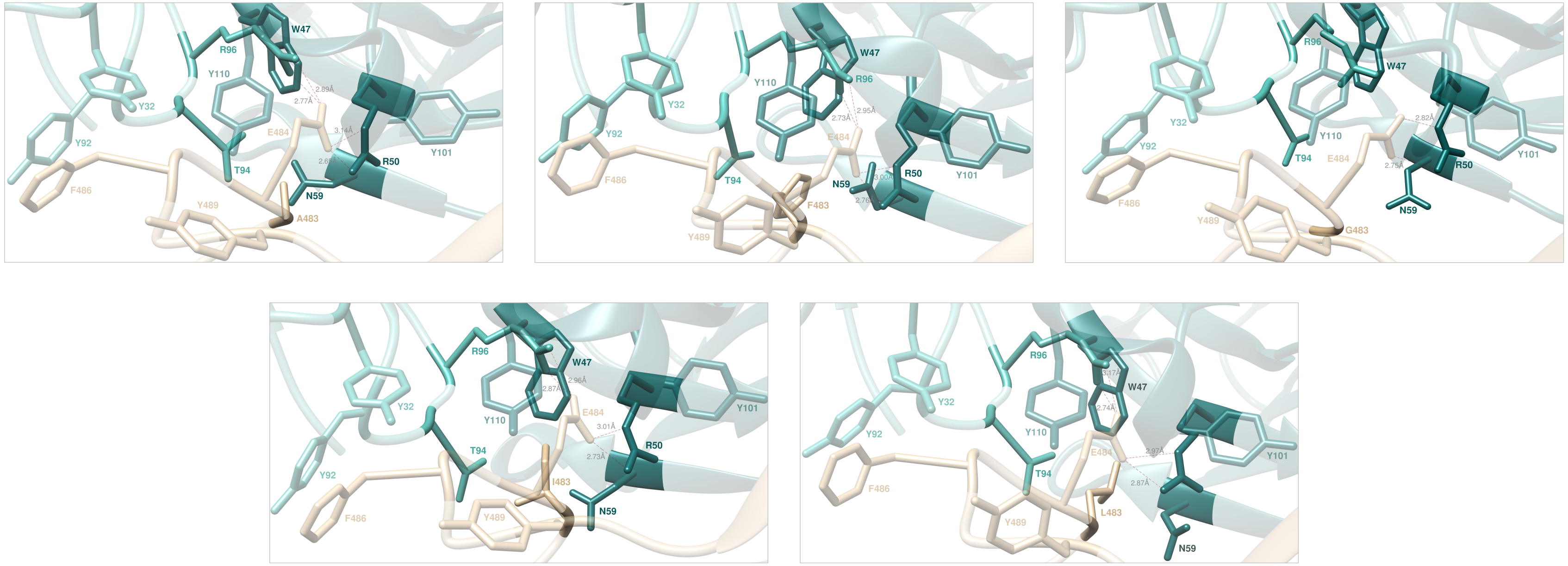


**Figure S4.** Main interactions involving the S-RBD$_{CoV-2}$ A483 (top left), F483 (top middle), G483 (top right), I483 (bottom left), and L483 (bottom right) at the interface with the LY-CoV555 antibody as obtained from the corresponding equilibrated MD simulations. The wild type V483 is presented and discussed in the main text (Figures 3A). Colors and other explanations as in Figure S1. For further details see Tables S1 and S8.

**Table S8.** Main interactions between the wild-type S-RBD$_{CoV-2}$ residue V483 and all considered mutants at the protein-protein interface detected during MD simulations of RBD of SARS-CoV-2 (COV2) in complex with the LY-CoV555 antibody. Acronyms and other explanations as in Table S5.

| **SB** | **COV2** | **LY-CoV555** | **V483** | **A483** | **F483** | **G483** | **I483** | **L483** |
|---|---|---|---|---|---|---|---|---|
| s-s | E484 | $R_H$50 | ✓(2.74,3.05) | ✓(2.65,3.14) | ✓(2.76,3.00) | ✓(2.75,2.82) | ✓(2.73,3.01) | ✓(2.87,2.97) |
| s-s | E484 | $R_L$96 | ✓(2.82,2.96) | ✓(2.77,2.89) | ✓(2.73,2.95) | ✗ | ✓(2.87,2.96) | ✓(2.74,3.17) |
| **CI** | **COV2** | **LY-CoV555** | **V483** | **A483** | **F483** | **G483** | **I483** | **L483** |
| vdW/h | E484 | $Y_H$101 | ✓ | ✓ | ✓ | ✓ | ✓ | ✓ |
| vdW/h | E484 | $Y_H$110 | ✓ | ✓ | ✓ | ✓ | ✓ | ✓ |
| **CI** | **COV2** | **LY-CoV555** | **V483** | **A483** | **F483** | **G483** | **I483** | **L483** |
| s-s | X483 | $W_H$47 | ✓ | ✓ | ✓ | ✗ | ✓ | ✓ |

| | | | | | | | | |
|---|---|---|---|---|---|---|---|---|
| s-s | X483 | $R_H50$ | ✓ | ✗ | ✓ | ✗ | ✓ | ✓ |
| s-s | X483 | $N_H59$ | ✓ | ✗ | ✓ | ✗ | ✓ | ✓ |
| s-s | X483 | $T_L94$ | ✓ | ✗ | ✓ | ✗ | ✓ | ✓ |
| s-s | X483 | $R_L96$ | ✓ | ✓ | ✓ | ✗ | ✓ | ✓ |
| π-π | F486 | $Y_L32$ | ✓ | ✓ | ✓ | ✓ | ✓ | ✓ |
| π-π | F486 | $Y_L92$ | ✓ | ✓ | ✓ | ✓ | ✓ | ✓ |
| vdW/h | Y489 | $Y_H110$ | ✓ | ✓ | ✓ | ✓ | ✓ | ✓ |
| p | Y489 | $Y_L32$ | ✓ | ✓ | ✓ | ✓ | ✓ | ✓ |

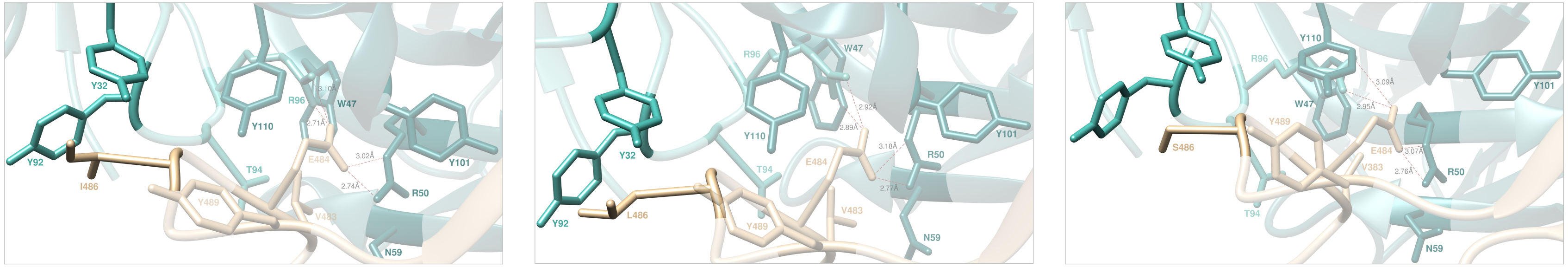


**Figure S5.** Main interactions involving the S-RBD$_{CoV\text{-}2}$ I486 (left), L486 (middle), and S486 (right) at the interface with the LY-CoV555 antibody as obtained from the corresponding equilibrated MD simulations. The wild type F486 is presented and discussed in the main text (Figures 3A). Colors and other explanations as in Figure S1. For further details see Tables S1 and S9.

**Table S9.** Main interactions between the wild-type S-RBD$_{CoV\text{-}2}$ residue V483 and all considered mutants at the protein-protein interface detected during MD simulations of RBD of SARS-CoV-2 (COV2) in complex with the LY-CoV555 antibody. Acronyms and other explanations as in Table S5.

| **SB** | **COV2** | **LY-CoV555** | **F486** | **I486** | **L486** | **S486** |
|---|---|---|---|---|---|---|
| s-s | E484 | R$_H$50 | ✓(2.74,3.05) | ✓(2.74,3.02) | ✓(2.77,3.18) | ✓(2.76,3.07) |
| s-s | E484 | R$_L$96 | ✓(2.82,2.96) | ✓(2.71,3.10) | ✓(2.89,2.92) | ✓(2.95,3.09) |
| **CI** | **COV2** | **LY-CoV555** | **F486** | **I486** | **L486** | **S486** |
| vdW/h | E484 | Y$_H$101 | ✓ | ✓ | ✓ | ✓ |
| vdW/h | E484 | Y$_H$110 | ✓ | ✓ | ✓ | ✓ |
| **CI** | **COV2** | **LY-CoV555** | **F486** | **I486** | **L486** | **S486** |
| s-s | V483 | W$_H$47 | ✓ | ✓ | ✓ | ✓ |
| s-s | V483 | R$_H$50 | ✓ | ✓ | ✓ | ✓ |
| s-s | V483 | N$_H$59 | ✓ | ✓ | ✓ | ✓ |
| s-s | V483 | T$_L$94 | ✓ | ✓ | ✓ | ✓ |
| s-s | V483 | R$_L$96 | ✓ | ✓ | ✓ | ✓ |
| $\pi$-$\pi$ | X486 | Y$_L$32 | ✓ | X(vdw) | X(vdw) | X |
| $\pi$-$\pi$ | X486 | Y$_L$92 | ✓ | X(vdw) | X(vdw) | X(vdw) |
| vdW/h | Y489 | Y$_H$110 | ✓ | ✓ | ✓ | ✓ |

| | | | | | | |
|---|---|---|---|---|---|---|
| p | Y489 | $Y_L32$ | ✓ | ✓ | ✓ | ✓ |

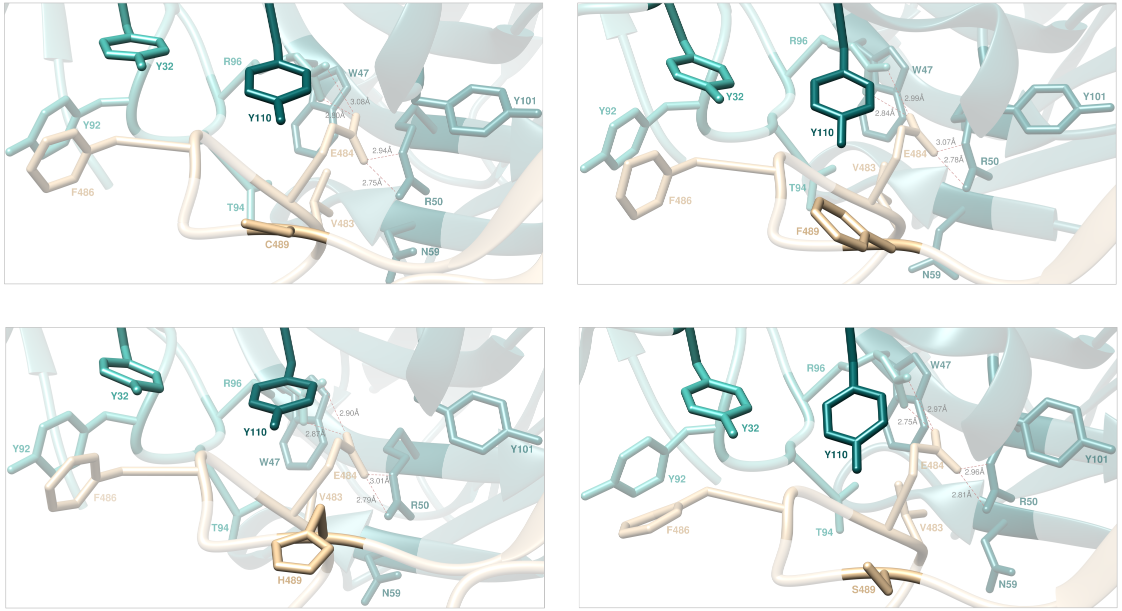


**Figure S6.** Main interactions involving the S-RBD$_{CoV\text{-}2}$ C489 (top left), F489 (top right), H489 (bottom left), and S486 (bottom right) at the interface with the LY-CoV555 antibody as obtained from the corresponding equilibrated MD simulations. The wild type Y489 is presented and discussed in the main text (Figures 3A). Colors and other explanations as in Figure S1. For further details see Tables S1 and S10.

**Table S10.** Main interactions between the wild-type S-RBD$_{CoV\text{-}2}$ residue Y489 and all considered mutants at the protein-protein interface detected during MD simulations of RBD of SARS-CoV-2 (COV2) in complex with the LY-CoV555 antibody. Acronyms and other explanations as in Table S5.

| **SB** | **COV2** | **LY-CoV555** | **Y489** | **C489** | **F489** | **H489** | **S489** |
|---|---|---|---|---|---|---|---|
| s-s | E484 | $R_H$50 | ✓(2.74,3.05) | ✓(2.75,2.94) | ✓(2.78,3.07) | ✓(2.79,3.01) | ✓(2.81,2.96) |
| s-s | E484 | $R_L$96 | ✓(2.82,2.96) | ✓(2.80,3.08) | ✓(2.84,2.99) | ✓(2.87,2.90) | ✓(2.75,2.97) |
| **CI** | **COV2** | **LY-CoV555** | **Y489** | **C489** | **F489** | **H489** | **S489** |
| vdW/h | E484 | $Y_H$101 | ✓ | ✓ | ✓ | ✓ | ✓ |
| vdW/h | E484 | $Y_H$110 | ✓ | ✓ | ✓ | ✓ | ✓ |
| **CI** | **COV2** | **LY-CoV555** | **Y489** | **C489** | **F489** | **H489** | **S489** |
| s-s | V483 | $W_H$47 | ✓ | ✓ | ✓ | ✓ | ✓ |

| | | | | | | | |
|---|---|---|---|---|---|---|---|
| s-s | V483 | $R_H50$ | ✓ | ✓ | ✓ | ✓ | ✓ |
| s-s | V483 | $N_H59$ | ✓ | ✓ | ✓ | ✓ | ✓ |
| s-s | V483 | $T_L94$ | ✓ | ✓ | ✓ | ✓ | ✓ |
| s-s | V483 | $R_L96$ | ✓ | ✓ | ✓ | ✓ | ✓ |
| π-π | F486 | $Y_L32$ | ✓ | ✓ | ✓ | ✓ | ✓ |
| π-π | F486 | $Y_L92$ | ✓ | ✓ | ✓ | ✓ | ✓ |
| vdW/h | X489 | $Y_H110$ | ✓ | ✓ | ✓ | ✓ | ✓ |
| p | X489 | $Y_L32$ | ✓ | ✗ | ✗ | ✗ | ✗ |

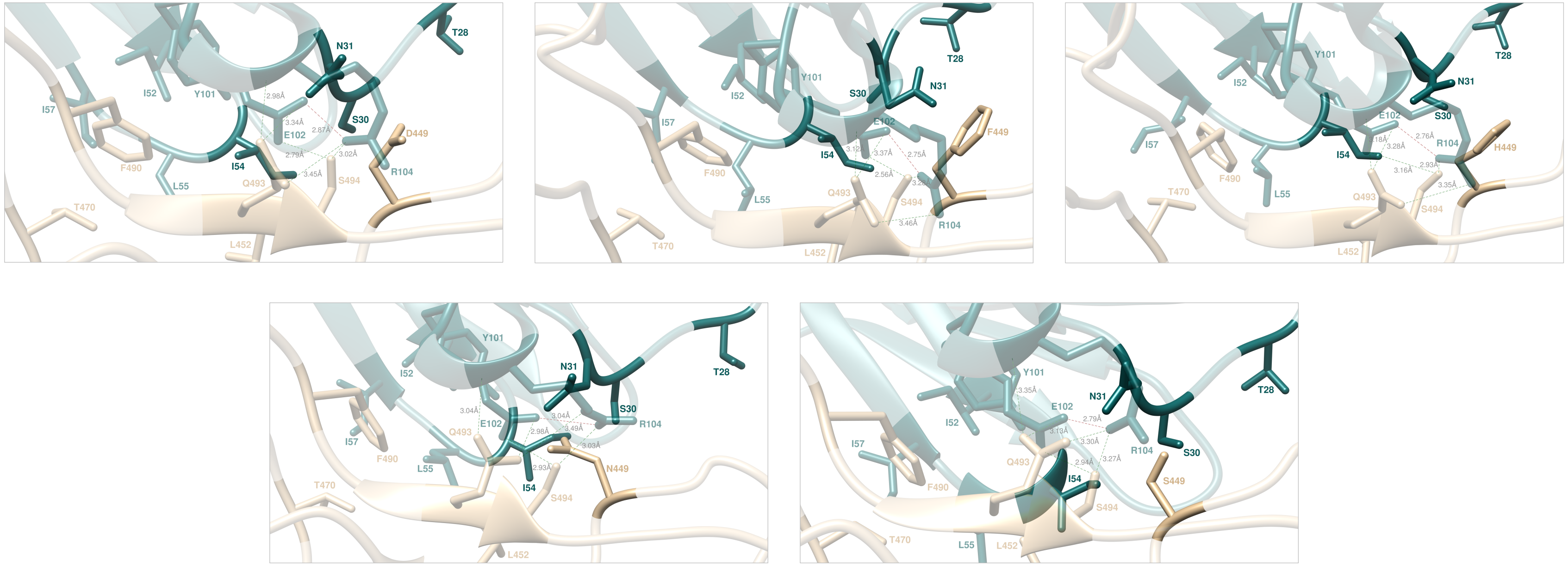


**Figure S7.** Main interactions involving the S-RBD$_{CoV-2}$ D449 (top left), F449 (top middle), H449 (top right), N449 (bottom left), and S449 (bottom right) at the interface with the LY-CoV555 antibody as obtained from the corresponding equilibrated MD simulations. The wild-type Y449 is presented and discussed in the main text (Figure 3B). For further details see Tables S1 and S11.

**Table S11.** Main interactions between the wild-type S-RBD$_{CoV-2}$ residue Y449 and all considered mutants at the protein-protein interface detected during MD simulations of the RBD of SARS-CoV-2 (COV2) in complex with the LY-CoV555 antibody. Acronyms and other explanations as in Table S5.

| **HB** | **COV2** | **LY-CoV555** | **Y449** | **D449** | **F449** | **H449** | **N449** | **S449** |
|---|---|---|---|---|---|---|---|---|
| s-s | Q493 | $E_H$102 | ✓(3.25) | ✓(3.34) | ✓(3.37) | ✓(3.28) | ✓(2.98) | ✓(3.13) |
| s-s | Q493 | $R_H$104 | ✓(3.31) | ✓(3.45) | ✓(3.46) | ✓(3.35) | ✓(3.49) | ✓(3.30) |
| s-b | Q493 | $R_H$104 | ✓(3.34) | ✓(2.98) | ✓(3.12) | ✓(3.18) | ✓(3.04) | ✓(3.35) |
| **HB** | **COV2** | **LY-CoV555** | **Y449** | **D449** | **F449** | **H449** | **N449** | **S449** |
| s-s | S494 | $E_H$102 | ✓(2.86) | ✓(2.79) | ✓(2.56) | ✓(3.16) | ✓(2.93) | ✓(2.94) |
| s-s | S494 | $R_H$104 | ✓(3.18) | ✓(3.02) | ✓(3.28) | ✓(2.93) | ✓(3.03) | ✓(3.27) |
| **CI** | **COV-2** | **LY-CoV555** | **Y449** | **D449** | **F449** | **H449** | **N449** | **S449** |

| | | | | | | | | |
|---|---|---|---|---|---|---|---|---|
| p | X449 | $T_H$28 | ✓ | ✗ | ✗(vdW/h) | ✓ | ✗ | ✗ |
| p | X449 | $S_H$30 | ✓ | ✓ | ✗ | ✓ | ✓ | ✓ |
| p | X449 | $N_H$31 | ✓ | ✓ | ✗(vdW/h) | ✓ | ✓ | ✓ |
| vdW/h | X449 | $I_H$54 | ✓ | ✗ | ✓ | ✓ | ✓ | ✗ |
| vdW/h | L452 | $I_H$54 | ✓ | ✓ | ✓ | ✓ | ✓ | ✓ |
| vdW/h | L452 | $L_H$55 | ✓ | ✓ | ✓ | ✓ | ✓ | ✓ |
| vdW/h | T470 | $L_H$55 | ✓ | ✓ | ✓ | ✓ | ✓ | ✓ |
| vdW/h | T470 | $I_H$57 | ✓ | ✓ | ✓ | ✓ | ✓ | ✓ |
| vdW/h | F490 | $I_H$52 | ✓ | ✓ | ✓ | ✓ | ✓ | ✓ |
| vdW/h | F490 | $L_H$55 | ✓ | ✓ | ✓ | ✓ | ✓ | ✓ |
| vdW/h | F490 | $I_H$57 | ✓ | ✓ | ✓ | ✓ | ✓ | ✓ |
| $\pi$-$\pi$ | F490 | $Y_H$101 | ✓ | ✓ | ✓ | ✓ | ✓ | ✓ |
| p | S494 | $N_H$31 | ✓ | ✓ | ✓ | ✓ | ✓ | ✓ |
| **SB** | **LY-CoV555** | **LY-CoV555** | **Y449** | **D449** | **F449** | **H449** | **N449** | **S449** |
| s-s | $E_H$102 | $R_H$104 | ✓(2.76) | ✓(2.87) | ✓(2.75) | ✓(2.76) | ✓(3.04) | ✓(2.79) |

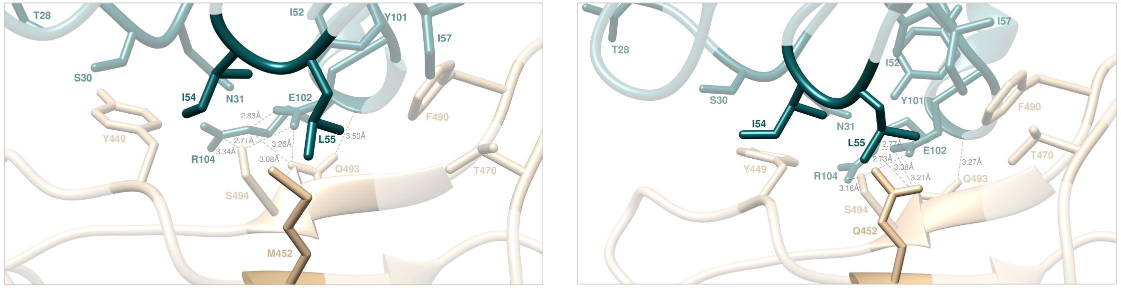


**Figure S8.** Main interactions involving the S-RBD$_{CoV-2}$ M452 (left) and Q452 (right) at the interface with the LY-CoV555 antibody as obtained from the corresponding equilibrated MD simulations. The wild type L452 and the R452 mutant complexes are presented and discussed in the main text (Figures 3B and 8E). Colors and other explanations as in Figure S1. For further details see Tables S1 and S12.

**Table S12.** Main interactions between the wild-type S-RBD$_{CoV-2}$ residue L452 and all considered mutants* at the protein-protein interface detected during MD simulations of RBD of SARS-CoV-2 (COV2) in complex with the LY-CoV555. Acronyms and other explanations as in Table S5. *Mutant R452 is discussed in detail in main text.

| **SB** | **COV2** | **LY-CoV555** | **L452** | **M452** | **Q452** | **R452** |
|---|---|---|---|---|---|---|
| s-s | X452 | $E_H$102 | ✗ | ✗ | ✗ | ✓(3.15) |
| **HB** | **COV2** | **LY-CoV555** | **L452** | **M452** | **Q452** | **R452** |
| s-s | Q493 | $E_H$102 | ✓(3.25) | ✓(3.26) | ✓(3.21) | ✗ |
| s-s | Q493 | $R_H$104 | ✓(3.31) | ✓(3.08) | ✓(3.38) | ✗(p) |
| s-b | Q493 | $R_H$104 | ✓(3.34) | ✓(3.50) | ✓(3.27) | ✓(2.87) |
| **HB** | **COV2** | **LY-CoV555** | **L452** | **M452** | **Q452** | **R452** |
| s-s | S494 | $E_H$102 | ✓(2.86) | ✓(2.71) | ✓(2.73) | ✗(p) |
| s-s | S494 | $R_H$104 | ✓(3.18) | ✓(3.34) | ✓(3.16) | ✗ |
| **CI** | **COV2** | **LY-CoV555** | **L452** | **M452** | **Q452** | **R452** |
| p | Y449 | $T_H$28 | ✓ | ✓ | ✓ | ✗ |
| p | Y449 | $S_H$30 | ✓ | ✓ | ✓ | ✗ |
| p | Y449 | $N_H$31 | ✓ | ✓ | ✓ | ✓ |
| vdW/h | Y449 | $I_H$54 | ✓ | ✓ | ✓ | ✗ |
| vdW/h | X452 | $I_H$54 | ✓ | ✓ | ✓ | ✓ |
| vdW/h | X452 | $L_H$55 | ✓ | ✓ | ✓ | ✗ |

| | | | | | | |
|---|---|---|---|---|---|---|
| vdW/h | T470 | $L_H$55 | ✓ | ✓ | ✓ | ✓ |
| vdW/h | T470 | $I_H$57 | ✓ | ✓ | ✓ | ✓ |
| vdW/h | F490 | $I_H$52 | ✓ | ✓ | ✓ | ✓ |
| vdW/h | F490 | $L_H$55 | ✓ | ✓ | ✓ | ✓ |
| vdW/h | F490 | $I_H$57 | ✓ | ✓ | ✓ | ✓ |
| $\pi$-$\pi$ | F490 | $Y_H$101 | ✓ | ✓ | ✓ | ✓ |
| p | S494 | $N_H$31 | ✓ | ✓ | ✓ | ✗ |
| **SB** | **LY-CoV555** | **LY-CoV555** | **L452** | **M452** | **Q452** | **R452** |
| s-s | $E_H$102 | $R_H$104 | ✓(2.76) | ✓(2.83) | ✓(2.77) | ✗ |

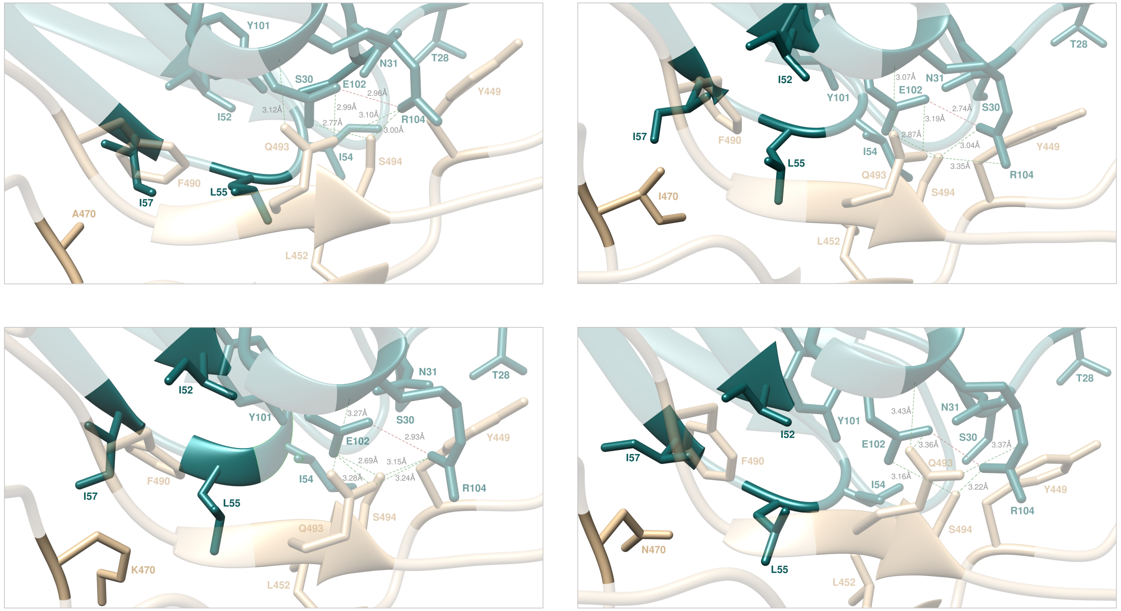


**Figure S9.** Main interactions involving the S-RBD$_{CoV-2}$ A470 (top left), I470 (top right), K470 (bottom left), and N470 (bottom right) at the interface with the LY-CoV555 antibody as obtained from the corresponding equilibrated MD simulations. The wild-type T470 is presented and discussed in the main text (Figure 3B). For further details see Tables S1 and S13.

**Table S13.** Main interactions between the wild-type S-RBD$_{CoV-2}$ residue T470 and all considered mutants at the protein-protein interface detected during MD simulations of the RBD of SARS-CoV-2 (COV2) in complex with the LY-CoV555 antibody.

| HB | COV2 | LY-CoV555 | T470 | A470 | I470 | K470 | N470 |
|---|---|---|---|---|---|---|---|
| s-s | Q493 | $E_H$102 | ✓(3.25) | ✓(2.99) | ✓(3.19) | ✓(3.28) | ✓(3.36) |
| s-s | Q493 | $R_H$104 | ✓(3.31) | ✓(3.16) | ✓(3.35) | ✓(3.24) | ✓(3.37) |
| s-b | Q493 | $R_H$104 | ✓(3.34) | ✓(3.12) | ✓(3.07) | ✓(3.27) | ✓(3.43) |
| **HB** | **COV2** | **LY-CoV555** | **T470** | **A470** | **I470** | **K470** | **N470** |
| s-s | S494 | $E_H$102 | ✓(2.86) | ✓(2.77) | ✓(2.87) | ✓(2.69) | ✓(3.16) |
| s-s | S494 | $R_H$104 | ✓(3.18) | ✓(3.00) | ✓(3.04) | ✓(3.15) | ✓(3.22) |
| **CI** | **COV2** | **LY-CoV555** | **T470** | **A470** | **I470** | **K470** | **N470** |

| | | | | | | | |
|---|---|---|---|---|---|---|---|
| p | Y449 | $T_H28$ | ✓ | ✓ | ✓ | ✓ | ✓ |
| p | Y449 | $S_H30$ | ✓ | ✓ | ✓ | ✓ | ✓ |
| p | Y449 | $N_H31$ | ✓ | ✓ | ✓ | ✓ | ✓ |
| vdW/h | Y449 | $I_H54$ | ✓ | ✓ | ✓ | ✓ | ✓ |
| vdW/h | L452 | $I_H54$ | ✓ | ✓ | ✓ | ✓ | ✓ |
| vdW/h | L452 | $L_H55$ | ✓ | ✓ | ✓ | ✓ | ✓ |
| vdW/h | X470 | $L_H55$ | ✓ | ✗ | ✓ | ✓ | ✓ |
| vdW/h | X470 | $I_H57$ | ✓ | ✓ | ✓ | ✓ | ✓ |
| vdW/h | F490 | $I_H52$ | ✓ | ✓ | ✓ | ✓ | ✓ |
| vdW/h | F490 | $L_H55$ | ✓ | ✓ | ✓ | ✓ | ✓ |
| vdW/h | F490 | $I_H57$ | ✓ | ✓ | ✓ | ✓ | ✓ |
| $\pi$-$\pi$ | F490 | $Y_H101$ | ✓ | ✓ | ✓ | ✓ | ✓ |
| p | S494 | $N_H31$ | ✓ | ✓ | ✓ | ✓ | ✓ |
| **SB** | **LY-CoV555** | **LY-CoV555** | **T470** | **A470** | **I470** | **K470** | **N470** |
| s-s | $E_H102$ | $R_H104$ | ✓(2.76) | ✓(2.96) | ✓(2.74) | ✓(2.93) | ✓(2.76) |

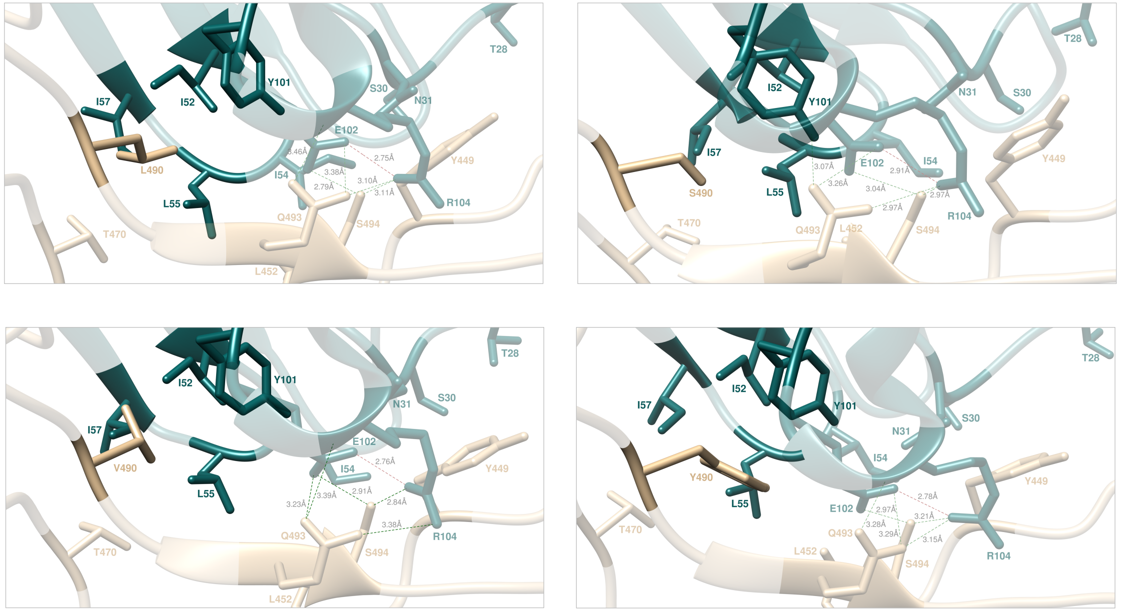


**Figure S10.** Main interactions involving the S-RBD$_{CoV-2}$ L490 (top left), S490 (top right), V490 (bottom left), and Y490 (bottom right) at the interface with the LY-CoV555 antibody as obtained from the corresponding equilibrated MD simulations. The wild-type F490 is presented and discussed in the main text (Figure 3B). For further details see Tables S1 and S13.

**Table S14.** Main interactions between the wild-type S-RBD$_{CoV-2}$ residue F490 and all considered mutants at the protein-protein interface detected during MD simulations of the RBD of SARS-CoV-2 (COV2) in complex with the LY-CoV555 antibody.

| HB | COV2 | LY-CoV555 | F490 | L490 | S490 | V490 | Y490 |
|---|---|---|---|---|---|---|---|
| s-s | Q493 | $E_H$102 | ✓(3.25) | ✓(3.38) | ✓(3.26) | ✓(3.23) | ✓(3.29) |
| s-s | Q493 | $R_H$104 | ✓(3.31) | ✓(3.10) | ✓(2.97) | ✓(3.38) | ✓(3.15) |
| s-b | Q493 | $R_H$104 | ✓(3.34) | ✓(3.46) | ✓(3.07) | ✓(3.39) | ✓(3.28) |
| **HB** | **COV2** | **LY-CoV555** | **F490** | **L490** | **S490** | **V490** | **Y490** |
| s-s | S494 | $E_H$102 | ✓(2.86) | ✓(2.79) | ✓(3.94) | ✓(2.91) | ✓(2.97) |
| s-s | S494 | $R_H$104 | ✓(3.18) | ✓(3.11) | ✓(2.97) | ✓(2.84) | ✓(3.21) |
| **CI** | **COV2** | **LY-CoV555** | **F490** | **L490** | **S490** | **V490** | **Y490** |

| | | | | | | | |
|---|---|---|---|---|---|---|---|
| p | Y449 | $T_H$28 | ✓ | ✓ | ✓ | ✓ | ✓ |
| p | Y449 | $S_H$30 | ✓ | ✓ | ✓ | ✓ | ✓ |
| p | Y449 | $N_H$31 | ✓ | ✓ | ✓ | ✓ | ✓ |
| vdW/h | Y449 | $I_H$54 | ✓ | ✓ | ✓ | ✓ | ✓ |
| vdW/h | L452 | $I_H$54 | ✓ | ✓ | ✓ | ✓ | ✓ |
| vdW/h | L452 | $L_H$55 | ✓ | ✓ | ✓ | ✓ | ✓ |
| vdW/h | T470 | $L_H$55 | ✓ | ✓ | ✓ | ✓ | ✓ |
| vdW/h | T470 | $I_H$57 | ✓ | ✓ | ✓ | ✓ | ✓ |
| vdW/h | X490 | $I_H$52 | ✓ | ✓ | ✗ | ✓ | ✓ |
| vdW/h | X490 | $L_H$55 | ✓ | ✓ | ✓ | ✓ | ✓ |
| vdW/h | X490 | $I_H$57 | ✓ | ✗ | ✗ | ✗ | ✓ |
| $\pi$-$\pi$ | X490 | $Y_H$101 | ✓ | ✗(vdW) | ✗ | ✗ | ✓ |
| p | S494 | $N_H$31 | ✓ | ✓ | ✓ | ✓ | ✓ |
| **SB** | **LY-CoV555** | **LY-CoV555** | **F490** | **L490** | **S490** | **V490** | **Y490** |
| s-s | $E_H$102 | $R_H$104 | ✓(2.76) | ✓(2.75) | ✓(2.91) | ✓(2.76) | ✓(2.78) |
