## Supporting information_part2 for "*In Silico* Molecular-Based Rationale for SARS-CoV-2 Spike Circulating Mutations Able to Escape Bamlanivimab and Etesevimab Monoclonal Antibodies"

**Table S15.** Main intermolecular and intramolecular interactions between residues at the protein-protein interface detected during MD simulations of the LY-Cov016 antibody in complex with the RBDs of SARS-CoV-2 S-RBD (COV2). HB = hydrogen bond; SB = salt bridge; CI = contact interactions, including van der Waals/hydrophobic (vdW/h), polar (p), $\pi/\pi$ and $\pi$/cation ($\pi$/c) interactions. In the HB column, s-s indicates side chain-side chain interactions while s-b or b-s indicate side chain-backbone or backbone-sidechain interactions, respectively.

| **HB** | **COV2** | **Length (Å)** | **LY-Cov016** |
|---|---|---|---|
| s-s | E406 | $2.84 \pm 0.27$ | $T_L$94 |
| s-s | T415 | $3.13 \pm 0.10$ | $S_H$56 |
| s-s | K417 | $2.98 \pm 0.14$ | $Y_H$52 |
| s-s | D420 | $3.07 \pm 0.15$ | $S_H$56 |
| s-s | Y421 | $3.09 \pm 0.11$ | $S_H$53 |
| s-b | | $3.21 \pm 0.14$ | $G_H$54 |
| s-b | N460 | $3.12 \pm 0.17$ | $G_H$54 |
| s-s | | $3.04 \pm 0.09$ | $S_H$56 |
| s-s | Y473 | $2.83 \pm 0.21$ | $S_H$31 |
| s-s | N487 | $2.86 \pm 0.15$ | $R_H$97 |
| s-s | Y489 | $3.38 \pm 0.13$ | $R_H$97 |
| s-b | Q493 | $3.03 \pm 0.18$ | $Y_H$102 |
| s-s | Y505 | $3.01 \pm 0.19$ | $Y_L$32 |
| b-s | | $3.14 \pm 0.13$ | $Y_L$92 |
| **SB** | **COV2** | **Length (Å)** | **LY-Cov016** |
| | K417 | $2.92 \pm 0.15$ | $D_H$104 |
| | | $3.10 \pm 0.17$ | |
| **CI** | **COV2** | **LY-Cov016** | |
| p | K417 | $Y_H$33 | |
| vdW/h | | $P_H$100 | |
| p | Y421 | $Y_H$33 | |
| vdW/h | | $Y_H$52 | |
| vdW/h | L455 | $Y_H$33 | |
| vdW/h | | $P_H$100 | |
| vdW/h | | $M_H$101 | |
| vdW/h | F456 | $Y_H$33 | |

| | | |
|---|---|---|
| vdW/h | | $S_{H}$53 |
| vdW/h | | $L_{H}$99 |
| vdW/h | | $P_{H}$100 |
| vdW/h | | $M_{H}$101 |
| p | Y473 | $S_{H}$53 |
| vdW/h | N487 | $F_{H}$27 |
| vdW/h | Y489 | $L_{H}$99 |
| vdW/h | | $M_{H}$101 |
| vdW/h | Q493 | $M_{H}$101 |
| vdW/h | | $Y_{H}$102 |

**Table S16.** Relative binding free energy and its components calculated by the combined computational alanine scanning mutagenesis – interaction entropy approach for the S-RBD of SARS-CoV-2 residues effectively involved in the binding interface with the LY-CoV016 antibody (see the SI Materials and Methods section for details). IE = interaction entropy. $\Delta\Delta G = \Delta G_{WILDTYPE} - \Delta G_{ALA}$ (see text for details).

| | E406A | T415A | K417A | D420A | Y421A | L455A | F456A | N460A | Y473A | N487A |
|---|---|---|---|---|---|---|---|---|---|---|
| $\Delta\Delta E_{DISP}$ | -0.25 | 0.08 | -0.81 | -0.30 | -1.04 | -1.37 | -2.62 | -1.21 | -0.44 | -0.47 |
| $\Delta\Delta E_{ELE}$ | -0.87 | -1.26 | -5.43 | -1.87 | -2.59 | 0.22 | 0.22 | -1.40 | -1.78 | -1.62 |
| $\Delta\Delta H$ | -1.12 | -1.18 | -6.24 | -2.17 | -3.63 | -1.15 | -2.40 | -2.61 | -2.22 | -2.09 |
| $\Delta\Delta IE$ | -0.17 | 0.02 | 0.23 | 0.16 | 0.16 | -0.05 | -0.09 | -0.14 | 0.14 | 0.06 |
| **$\Delta\Delta G_{ACE2}$** | **-1.29 (0.13)** | **-1.16 (0.14)** | **-6.01 (0.10)** | **-2.01 (0.11)** | **-2.47 (0.18)** | **-1.20 (0.16)** | **-2.49 (0.13)** | **-2.75 (0.11)** | **-2.08 (0.12)** | **-2.03 (0.16)** |

| | Y489A | Q493A | Y505A |
|---|---|---|---|
| $\Delta\Delta E_{DISP}$ | -1.49 | -0.58 | -0.83 |
| $\Delta\Delta E_{ELE}$ | -0.58 | -0.92 | -1.23 |
| $\Delta\Delta H$ | -2.07 | -1.50 | -2.06 |
| $\Delta\Delta IE$ | 0.23 | -0.09 | 0.19 |
| **$\Delta\Delta G_{ACE2}$** | **-1.84 (0.09)** | **-1.59 (0.14)** | **-1.87 (0.12)** |

**Table S17.** Relative binding free energy and its components calculated by the combined computational alanine scanning mutagenesis – interaction entropy approach for LY-Cov016 antibody residues effectively involved in the binding interface with the S-RBD of SARS-CoV-2 (see the SI Materials and Methods section for details). IE = interaction entropy. $\Delta\Delta G = \Delta G_{WILDTYPE} - \Delta G_{ALA}$ (see text for details).

| | $S_H$31A | $Y_H$33A | $Y_H$52A | $S_H$53A | $S_H$56A | $R_H$97A | $L_H$99A | $P_H$100A | $M_H$101A | $Y_H$102A |
|---|---|---|---|---|---|---|---|---|---|---|
| $\Delta\Delta E_{DISP}$ | -0.03 | -0.44 | -0.88 | -1.05 | -1.49 | -0.27 | -0.92 | -1.22 | -1.89 | -0.50 |
| $\Delta\Delta E_{ELE}$ | -0.88 | -2.01 | -0.83 | -1.47 | -2.40 | -1.99 | 0.08 | 0.12 | 0.04 | -1.32 |
| $\Delta\Delta H$ | -0.91 | -2.45 | -1.71 | -2.52 | -3.89 | -2.26 | -0.84 | -1.10 | -1.85 | -1.82 |
| $\Delta\Delta IE$ | -0.07 | 0.08 | 0.16 | -0.04 | 0.09 | 0.23 | -0.05 | -0.17 | 0.07 | 0.19 |
| **$\Delta\Delta G_{CoV-2}$** | **-0.98 (0.17)** | **-2.37 (0.12)** | **-1.55 (0.11)** | **-2.56 (0.15)** | **-3.80 (0.16)** | **-2.03 (0.08)** | **-0.89 (0.15)** | **-1.27 (0.10)** | **-1.78 (0.14)** | **-1.63 (0.18)** |

| | $D_H$104A | $Y_L$32A | $Y_L$92A | $T_L$94A |
|---|---|---|---|---|
| $\Delta\Delta E_{DISP}$ | -0.18 | -0.33 | -0.28 | 0.16 |
| $\Delta\Delta E_{ELE}$ | -2.84 | -0.92 | -0.86 | -1.15 |
| $\Delta\Delta H$ | -3.02 | -1.25 | -1.14 | -0.99 |
| $\Delta\Delta IE$ | 0.21 | 0.13 | 0.15 | -0.16 |
| **$\Delta\Delta G_{CoV-2}$** | **-2.81 (0.15)** | **-1.12 (0.10)** | **-0.99 (0.16)** | **-1.15 (0.12)** |

**Table S18.** Relative binding free energy and its components for the mutated S-RBD of SARS-CoV-2 residues in the binding interface with the LY-Cov016 antibody. IE = interaction entropy. IE = interaction entropy. $\Delta\Delta G = \Delta G_{WILDTYPE} - \Delta G_{MUTANT}$ (see text for details).

| | **E406D** | **E406Q** | **T415A** | **T415I** | **T415N** | **T415P** | **T415S** | **K417E** | **K417N** | **K417R** | **K417T** | **D420A** |
|---|---|---|---|---|---|---|---|---|---|---|---|---|
| $\Delta\Delta E_{DISP}$ | -0.23 | -0.18 | -0.59 | -0.37 | -0.61 | -1.32 | -0.32 | -2.76 | -2.37 | -0.31 | -2.99 | -1.89 |
| $\Delta\Delta E_{ELE}$ | 0.14 | -0.11 | -1.34 | -1.21 | -0.70 | -1.64 | -0.03 | -4.64 | -4.75 | -0.80 | -4.12 | -2.61 |
| $\Delta\Delta H$ | -0.09 | -0.29 | -1.93 | -1.58 | -1.31 | -2.96 | -0.35 | -7.40 | -7.13 | -1.11 | -7.11 | -4.50 |
| $\Delta\Delta IE$ | 0.02 | 0.07 | 0.06 | -0.07 | 0.10 | 0.13 | -0.05 | -0.16 | -0.15 | 0.12 | -0.03 | 0.14 |
| $\boldsymbol{\Delta\Delta G_{CoV2}}$ | **-0.07** | **-0.22** | **-1.87** | **-1.65** | **-1.21** | **-2.83** | **-0.40** | **-7.56** | **-7.27** | **-0.99** | **-7.14** | **-4.36** |
| | **(0.18)** | **(0.16)** | **(0.14)** | **(0.18)** | **(0.17)** | **(0.07)** | **(0.13)** | **(0.18)** | **(0.07)** | **(0.15)** | **(0.09)** | **(0.17)** |

| | **D420G** | **D420N** | **L455F** | **L455S** | **L455V** | **F456L** | **F456Y** | **N460I** | **N460K** | **N460S** | **N460T** | **Y473F** |
|---|---|---|---|---|---|---|---|---|---|---|---|---|
| $\Delta\Delta E_{DISP}$ | -2.06 | -2.72 | 0.15 | -1.07 | -0.83 | -0.58 | -0.19 | -1.97 | -1.70 | -2.35 | -1.94 | -0.33 |
| $\Delta\Delta E_{ELE}$ | -2.54 | -1.45 | -0.22 | 0.16 | 0.17 | -0.09 | -0.03 | -3.03 | -1.37 | -1.84 | -2.06 | -1.50 |
| $\Delta\Delta H$ | -4.60 | -4.17 | -0.07 | -0.91 | -0.66 | -0.67 | -0.22 | -5.00 | -3.07 | -4.19 | -4.00 | -1.83 |
| $\Delta\Delta IE$ | 0.21 | -0.06 | -0.11 | 0.08 | 0.07 | 0.06 | 0.18 | -0.01 | -0.21 | 0.13 | -0.12 | 0.16 |
| $\boldsymbol{\Delta\Delta G_{CoV-2}}$ | **-4.39** | **-4.23** | **-0.18** | **-0.83** | **-0.59** | **-0.61** | **-0.04** | **-5.01** | **-3.28** | **-4.06** | **-4.12** | **-1.67** |
| | **(0.10)** | **(0.10)** | **(0.17)** | **(0.11)** | **(0.14)** | **(0.16)** | **(0.17)** | **(0.14)** | **(0.18)** | **(0.09)** | **(0.14)** | **(0.08)** |

| | **Y473H** | **N487D** | **Y489C** | **Y489F** | **Y489H** | **Y489S** | **Q493H** | **Q493K** | **Q493L** | **Q493R** | **Y505F** | **Y505H** |
|---|---|---|---|---|---|---|---|---|---|---|---|---|
| $\Delta\Delta E_{DISP}$ | -0.07 | -0.68 | -0.77 | -0.49 | -0.56 | -1.01 | -0.23 | -0.42 | -0.45 | -0.37 | -0.23 | -0.34 |
| $\Delta\Delta E_{ELE}$ | -0.07 | 0.12 | -1.46 | -1.24 | -0.68 | -1.59 | -0.52 | -0.83 | -1.31 | -0.75 | -0.21 | 0.12 |
| $\Delta\Delta H$ | -0.14 | -0.56 | -2.23 | -1.73 | -1.24 | -2.60 | -0.75 | -1.25 | -1.76 | -1.12 | -0.44 | -0.22 |
| $\Delta\Delta IE$ | -0.05 | -0.14 | 0.08 | 0.15 | -0.10 | 0.19 | 0.03 | -0.14 | -0.17 | -0.16 | -0.04 | 0.01 |
| $\boldsymbol{\Delta\Delta G_{CoV-2}}$ | **-0.19** | **-0.70** | **-2.15** | **-1.58** | **-1.34** | **-2.41** | **-0.72** | **-1.39** | **-1.93** | **-1.28** | **-0.48** | **-0.21** |
| | **(0.16)** | **(0.09)** | **(0.10)** | **(0.11)** | **(0.16)** | **(0.07)** | **(0.12)** | **(0.09)** | **(0.13)** | **(0.16)** | **(0.16)** | **(0.08)** |

| | **Y505W** |
|---|---|
| $\Delta\Delta E_{DISP}$ | 0.15 |
| $\Delta\Delta E_{ELE}$ | -0.35 |
| $\Delta\Delta H$ | -0.20 |
| $\Delta\Delta IE$ | 0.11 |
| $\boldsymbol{\Delta\Delta G_{ACE2}}$ | **-0.09** |
| | **(0.14)** |

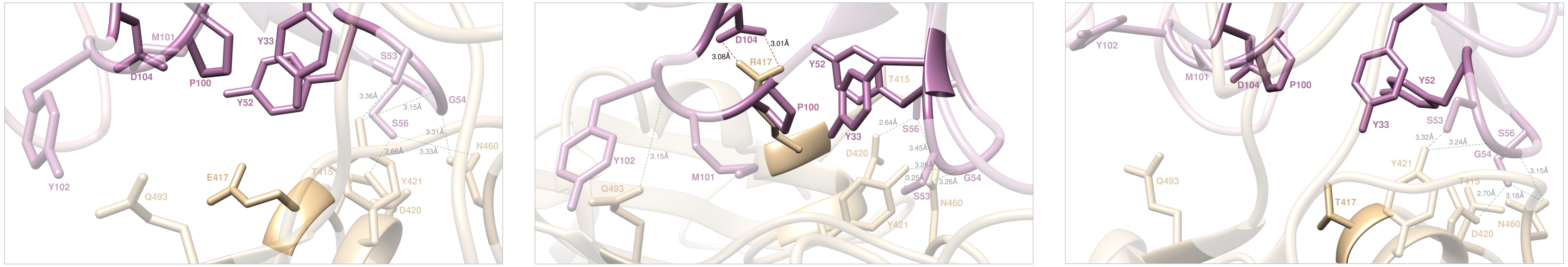


**Figure S11.** Main interactions involving the S-RBD$_{CoV-2}$ E417 (left), R417 (middle), and T417 (right) at the interface with the LY-Cov016 antibody as obtained from the corresponding equilibrated MD simulations. The wild-type K417 and the N417 mutant complexes are presented and discussed in the main text (Figures 10A and 11B). In this and all remaining Figures, the secondary structure of the S-RBD$_{CoV-2}$ is shown as a light tan ribbon, while the heavy and light chains of the LY-Cov016 antibody are portrayed as light mulberry and light pink icing ribbons, respectively. Each protein residue under discussion and all other residues directly interacting with it are highlighted in dark matching-colored sticks and labelled; further residues/interactions related to the residue under investigation are evidenced in light matching-colored sticks and labelled in light gray. Hydrogen bonds and salt bridges directly involving the residue under discussion are represented as dark green and dark red broken lines, respectively, and the relevant average distances are reported (in black) accordingly. New HBs and SBs eventually detected in each mutant complex are also indicated using dark green/red broken lines and black labels. Further important HBs and SBs detected in each complex are also indicated using light green/red broken lines and light gray labels. For further details see Tables S15 and S19.

**Table S19.** Main intermolecular interactions between the wild-type S-RBD$_{CoV-2}$ residue K417 and all considered mutants* at the protein-protein interface detected during MD simulations of the RBD of SARS-CoV-2 (COV-2) in complex with the LY-Cov016 antibody. HB = hydrogen bond; SB = salt bridge; CI = contact interactions, including van der Waals/hydrophobic (vdW/h), polar (p), $\pi/\pi$ and $\pi$/cation ($\pi$/c) interactions. In the HB column, s-s indicates side chain-side chain interactions while s-b or b-s indicate side chain-backbone or backbone-side chain interactions, respectively. Preserved/new or lost interactions are marked with the symbols ✓ and ✗, respectively. Relevant changes in the type/nature of the interactions are indicated in parenthesis. For HBs and SBs, the relevant average lengths (in Å) are also reported (their standard deviations, all within 10%, are not shown for clarity). *Mutant N417 is discussed in detail in main text.

| **SB** | **COV-2** | **LY-Cov016** | **K417** | **E417** | **N417** | **R417** | **T417** |
|---|---|---|---|---|---|---|---|
| | X417 | $D_H$104 | ✓(2.92,3.10) | ✗ | ✗ | ✓(3.01,3.08) | ✗ |
| **HB** | **COV-2** | **LY-Cov016** | **K417** | **E417** | **N417** | **R417** | **T417** |
| s-s | T415 | $S_H$56 | ✓(3.13) | ✗ | ✗(p) | ✗(p) | ✗(p) |
| **HB** | **COV-2** | **LY-Cov016** | **K417** | **E417** | **N417** | **R417** | **T417** |
| s-s | X417 | $Y_H$52 | ✓(2.98) | ✗(vdW/h) | ✗ | ✗(p) | ✗ |
| **HB** | **COV-2** | **LY-Cov016** | **K417** | **E417** | **N417** | **R417** | **T417** |
| s-s | D420 | $S_H$56 | ✓(3.07) | ✓(2.66) | ✓(2.92) | ✓(2.64) | ✓(2.70) |
| **HB** | **COV-2** | **LY-Cov016** | **K417** | **E417** | **N417** | **R417** | **T417** |

| | | | | | | | |
|---|---|---|---|---|---|---|---|
| s-s | Y421 | $S_H53$ | ✓(3.09) | ✓(3.36) | ✗(p) | ✓(3.25) | ✓(3.32) |
| s-b | Y421 | $G_H54$ | ✓(3.21) | ✓(3.15) | ✓(3.20) | ✓(3.26) | ✓(3.24) |
| **HB** | **COV-2** | **LY-Cov016** | **K417** | **E417** | **N417** | **R417** | **T417** |
| s-b | N460 | $G_H54$ | ✓(3.12) | ✓(3.31) | ✗ | ✓(3.26) | ✓(3.15) |
| s-s | N460 | $S_H56$ | ✓(3.04) | ✓(3.33) | ✓(3.41) | ✓(3.45) | ✓(3.18) |
| **HB** | **COV-2** | **LY-Cov016** | **K417** | **E417** | **N417** | **R417** | **T417** |
| s-b | Q493 | $Y_H102$ | ✓(3.03) | ✗(p) | ✓(3.41) | ✓(3.15) | ✗ |
| **CI** | **COV-2** | **LY-Cov016** | **K417** | E417 | N417 | R417 | T417 |
| p | X417 | $Y_H33$ | ✓ | ✗ | ✗ | ✓ | ✗ |
| vdW/h | X417 | $P_H100$ | ✓ | ✗ | ✗ | ✓ | ✗ |
| vdW/h | Y421 | $Y_H52$ | ✓ | ✓ | ✓ | ✓ | ✓ |
| p | Y421 | $Y_H33$ | ✓ | ✓ | ✓ | ✓ | ✓ |
| vdW/h | Q493 | $M_H101$ | ✓ | ✓ | ✓ | ✓ | ✓ |
| vdW/h | Q493 | $Y_H102$ | ✓ | ✗ | ✓ | ✓ | ✓ |

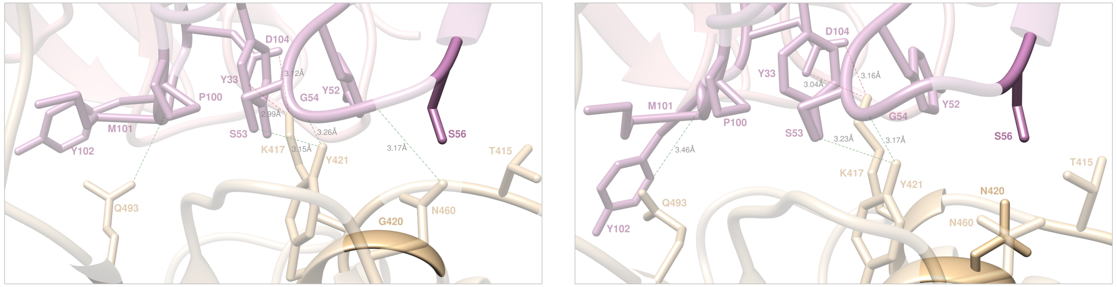


**Figure S12.** Main interactions involving the S-RBD$_{CoV-2}$ G420 (left) and N420 (right) at the interface with the LY-Cov016 antibody as obtained from the corresponding equilibrated MD simulations. The wild type D420 and the A420 mutant complexes are presented and discussed in the main text (Figures 10A and 12B). Colors and other explanations as in Figure S11. For further details see Tables S15 and S20.

**Table S20.** Main intermolecular and intramolecular interactions between the wild-type S-RBD$_{CoV-2}$ residue D420 and all considered mutants* at the protein-protein interface detected during MD simulations of RBD of SARS-CoV-2 (COV-2) in complex with the LY-Cov016 antibody. Acronyms and other explanations as in Table S19. *Mutant A420 is discussed in detail in main text.

| **SB** | **COV-2** | **LY-Cov016** | **D420** | **A420** | **G420** | **N420** |
|---|---|---|---|---|---|---|
| | K417 | $D_H104$ | √(2.92,3.10) | √(2.93,3.03) | √(2.99,3.12) | √(3.04,3.16) |
| **HB** | **COV-2** | **LY-Cov016** | **D420** | **A420** | **G420** | **N420** |
| s-s | T415 | $S_H56$ | √(3.13) | X | X | X(p) |
| **HB** | **COV-2** | **LY-Cov016** | **D420** | **A420** | **G420** | **N420** |
| s-s | K417 | $Y_H52$ | √(2.98) | X(p) | X(p) | X(p) |
| **HB** | **COV-2** | **LY-Cov016** | **D420** | **A420** | **G420** | **N420** |
| s-s | X420 | $S_H56$ | √(3.07) | X | X | X(p) |
| **HB** | **COV-2** | **LY-Cov016** | **D420** | **A420** | **G420** | **N420** |
| s-s | Y421 | $S_H53$ | √(3.09) | √(3.08) | √(3.15) | √(3.23) |
| s-b | Y421 | $G_H54$ | √(3.21) | √(3.25) | √(3.26) | √(3.17) |
| **HB** | **COV-2** | **LY-Cov016** | **D420** | **A420** | **G420** | **N420** |
| s-b | N460 | $G_H54$ | √(3.12) | X | √(3.17) | X |
| s-s | N460 | $S_H56$ | √(3.04) | √(3.40) | X(p) | X(p) |
| **HB** | **COV-2** | **LY-Cov016** | **D420** | **A420** | **G420** | **N420** |

| | | | | | | |
|---|---|---|---|---|---|---|
| s-b | Q493 | $Y_H102$ | √(3.03) | √(3.11) | √(3.30) | √(3.46) |
| **CI** | **COV-2** | **LY-Cov016** | **D420** | **A420** | **G420** | **N420** |
| p | K417 | $Y_H33$ | √ | √ | √ | √ |
| vdW/h | K417 | $P_H100$ | √ | √ | √ | √ |
| vdW/h | K421 | $Y_H52$ | √ | √ | √ | √ |
| p | Y421 | $Y_H33$ | √ | √ | √ | √ |
| vdW/h | Q493 | $M_H101$ | √ | √ | √ | √ |
| vdW/h | Q493 | $Y_H102$ | √ | √ | √ | √ |

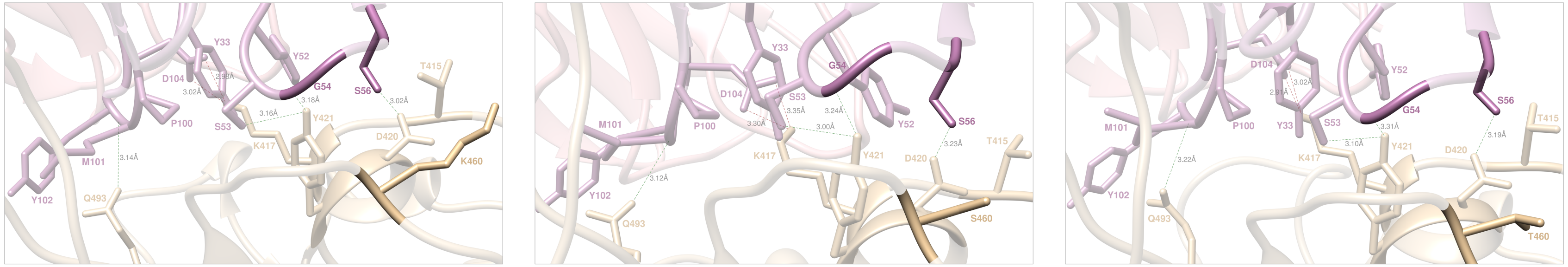


**Figure S13.** Main interactions involving the S-RBD$_{CoV-2}$ K460 (left), S460 (middle), and T460 (right) at the interface with the LY-Cov016 antibody as obtained from the corresponding equilibrated MD simulations. The wild type N460 and the I460 mutant complexes are presented and discussed in the main text (Figures 10A and 12D). Colors and other explanations as in Figure S11. For further details see Tables S15 and S21.

**Table S21.** Main intermolecular and intramolecular interactions between the wild-type S-RBD$_{CoV-2}$ residue N460 and all considered mutants* at the protein-protein interface detected during MD simulations of RBD of SARS-CoV-2 (COV-2) in complex with the LY-Cov016 antibody. Acronyms and other explanations as in Table S19. *Mutant I460 is discussed in detail in main text.

| **SB** | **COV-2** | **LY-Cov016** | **N460** | **I460** | **K460** | **S460** | **T460** |
|---|---|---|---|---|---|---|---|
| | K417 | D$_H$104 | √(2.92,3.10) | √(3.01,3.14) | √(2.98,3.02) | √(3.30,3.35) | √(2.91,3.02) |
| **HB** | **COV-2** | **LY-Cov016** | **N460** | **I460** | **K460** | **S460** | **T460** |
| s-s | T415 | S$_H$56 | √(3.13) | X(p) | X(p) | X(p) | X(p) |
| **HB** | **COV-2** | **LY-Cov016** | **N460** | **I460** | **K460** | **S460** | **T460** |
| s-s | K417 | Y$_H$52 | √(2.98) | X(p) | X(p) | X(p) | X(p) |
| **HB** | **COV-2** | **LY-Cov016** | **N460** | **I460** | **K460** | **S460** | **T460** |
| s-s | D420 | S$_H$56 | √(3.07) | √(2.65) | √(3.02) | √(3.23) | √(3.19) |
| **HB** | **COV-2** | **LY-Cov016** | **N460** | **I460** | **K460** | **S460** | **T460** |
| s-s | Y421 | S$_H$53 | √(3.09) | X(p) | √(3.16) | √(3.00) | √(3.10) |
| s-b | Y421 | G$_H$54 | √(3.21) | √(3.17) | √(3.18) | √(3.24) | √(3.31) |
| **HB** | **COV-2** | **LY-Cov016** | **N460** | **I460** | **K460** | **S460** | **T460** |
| s-b | X460 | G$_H$54 | √(3.12) | X | X | X | X |
| s-s | X460 | S$_H$56 | √(3.04) | X | X | X | X |
| **HB** | **COV-2** | **LY-Cov016** | **N460** | **I460** | **K460** | **S460** | **T460** |

| | | | | | | | |
|---|---|---|---|---|---|---|---|
| s-b | Q493 | $Y_H102$ | ✓(3.03) | ✓(3.10) | ✓(3.14) | ✓(3.12) | ✓(3.22) |
| **CI** | **COV-2** | **LY-Cov016** | **N460** | **I460** | **K460** | **S460** | **T460** |
| p | K417 | $Y_H33$ | ✓ | ✓ | ✓ | ✓ | ✓ |
| vdW/h | K417 | $P_H100$ | ✓ | ✓ | ✓ | ✓ | ✓ |
| vdW/h | K421 | $Y_H52$ | ✓ | ✓ | ✓ | ✓ | ✓ |
| p | Y421 | $Y_H33$ | ✓ | ✓ | ✓ | ✓ | ✓ |
| vdW/h | Q493 | $M_H101$ | ✓ | ✓ | ✓ | ✓ | ✓ |
| vdW/h | Q493 | $Y_H102$ | ✓ | ✓ | ✓ | ✓ | ✓ |

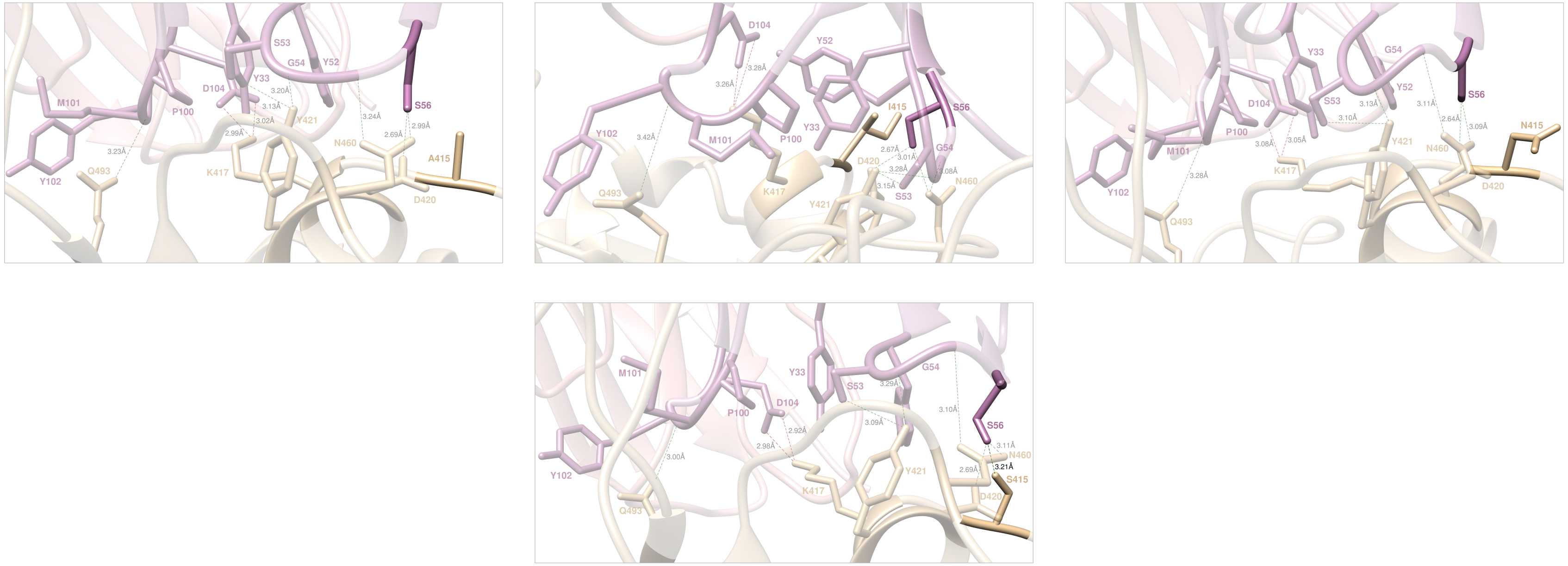

**Figure S14.** Main interactions involving the S-RBD$_{CoV-2}$ A415 (top left), I415 (top middle), N415 (top right), and S415 (bottom) at the interface with LY-Cov016 as obtained from the corresponding equilibrated MD simulations. The wild type T415 and the P415 mutant complexes are presented and discussed in the main text (Figures 10A and 13C). Colors and other explanations as in Figure S11. For further details see Tables S15 and S22.

**Table S22.** Main intermolecular and intramolecular interactions between the wild-type S-RBD$_{CoV-2}$ residue T415 and all considered mutants* at the protein-protein interface detected during MD simulations of RBD of SARS-CoV-2 (COV-2) in complex with the LY-Cov016. Acronyms and other explanations as in Table S19. *Mutant P415 is discussed in detail in main text.

| **SB** | **COV-2** | **LY-Cov016** | **T415** | **A415** | **I415** | **N415** | **P415** | **S415** |
|---|---|---|---|---|---|---|---|---|
| | K417 | $D_H$104 | ✓(2.92,3.10) | ✓(3.02,2.99) | ✓(3.26,3.28) | ✓(3.08,3.05) | ✓(2.95,2.94) | ✓(2.92,2.98) |
| **HB** | **COV-2** | **LY-Cov016** | **T415** | **A415** | **I415** | **N415** | **P415** | **S415** |
| s-s | X415 | $S_H$56 | ✓(3.13) | ✗ | ✗(vdW/h) | ✗(p) | ✗(vdW/h) | ✓(3.21) |
| **HB** | **COV-2** | **LY-Cov016** | **T415** | **A415** | **I415** | **N415** | **P415** | **S415** |
| s-s | K417 | $Y_H$52 | ✓(2.98) | ✗(p) | ✗(p) | ✗(p) | ✗(p) | ✗(p) |
| **HB** | **COV-2** | **LY-Cov016** | **T415** | **A415** | **I415** | **N415** | **P415** | **S415** |
| s-s | D420 | $S_H$56 | ✓(3.07) | ✓(2.69) | ✓(2.67) | ✓(2.64) | ✓(2.84) | ✓(2.69) |

| **HB** | **COV-2** | **LY-Cov016** | **T415** | **A415** | **I415** | **N415** | **P415** | **S415** |
|---|---|---|---|---|---|---|---|---|
| s-s | Y421 | $S_H$53 | ✓(3.09) | ✓(3.13) | ✓(3.15) | ✓(3.10) | ✓(3.16) | ✓(3.09) |
| s-b | Y421 | $G_H$54 | ✓(3.21) | ✓(3.20) | ✓(3.28) | ✓(3.13) | ✓(3.21) | ✓(3.29) |
| **HB** | **COV-2** | **LY-Cov016** | **T415** | **A415** | **I415** | **N415** | **P415** | **S415** |
| s-b | N460 | $G_H$54 | ✓(3.12) | ✓(3.24) | ✓(3.08) | ✓(3.11) | ✓(3.19) | ✓(3.10) |
| s-s | N460 | $S_H$56 | ✓(3.04) | ✓(2.99) | ✓(3.01) | ✓(3.09) | ✓(3.05) | ✓(3.11) |
| **HB** | **COV-2** | **LY-Cov016** | **T415** | **A415** | **I415** | **N415** | **P415** | **S415** |
| s-b | Q493 | $Y_H$102 | ✓(3.03) | ✓(3.23) | ✓(3.42) | ✓(3.28) | ✗(p) | ✓(3.00) |
| **CI** | **COV-2** | **LY-Cov016** | **T415** | **A415** | **I415** | **N415** | **P415** | **S415** |
| p | K417 | $Y_H$33 | ✓ | ✓ | ✓ | ✓ | ✓ | ✓ |
| vdW/h | K417 | $P_H$100 | ✓ | ✓ | ✓ | ✓ | ✓ | ✓ |
| vdW/h | Y421 | $Y_H$52 | ✓ | ✓ | ✓ | ✓ | ✓ | ✓ |
| p | Y421 | $Y_H$33 | ✓ | ✓ | ✓ | ✓ | ✓ | ✓ |
| vdW/h | Q493 | $M_H$101 | ✓ | ✓ | ✓ | ✓ | ✓ | ✓ |
| vdW/h | Q493 | $Y_H$102 | ✓ | ✓ | ✓ | ✓ | ✗ | ✓ |

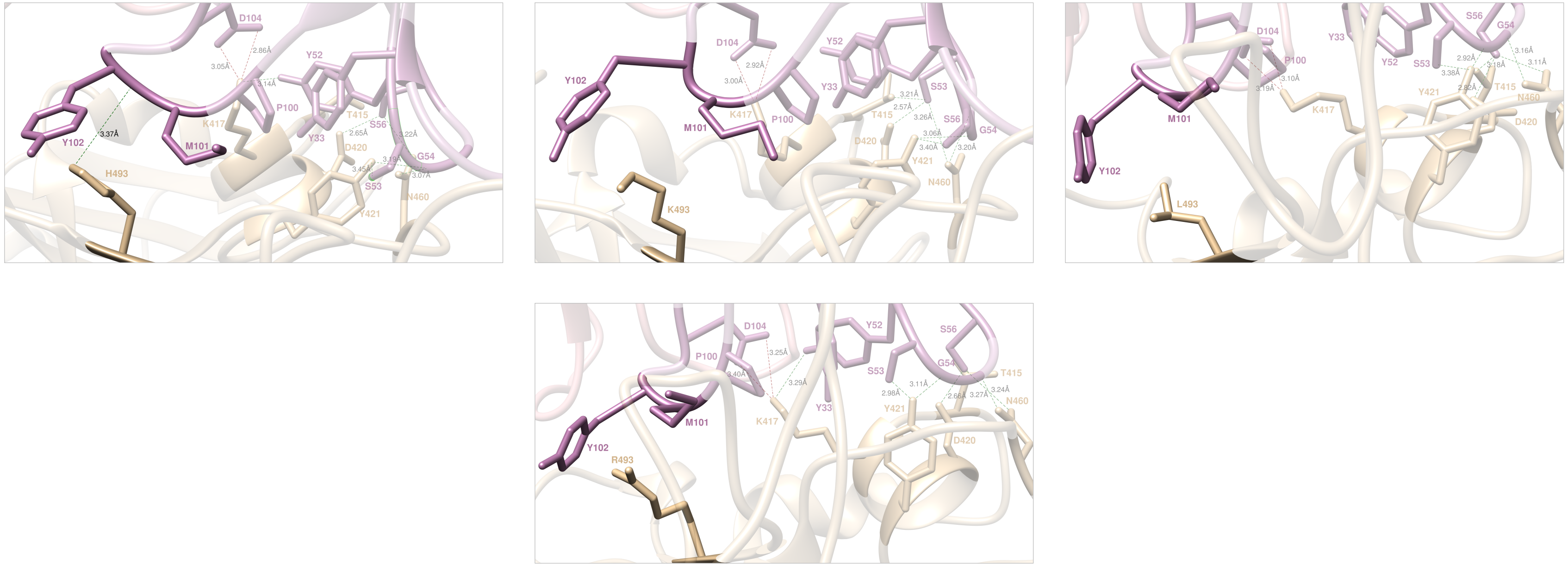

**Figure S15.** Main interactions involving the S-RBD$_{CoV-2}$ H493 (top left), K493 (top middle), L493 (top right), and R493 (bottom) at the interface with LY-Cov016 as obtained from the corresponding equilibrated MD simulations. The wild-type Q493 is presented and discussed in the main text (Figure 10A). Colors and other explanations as in Figure S11. For further details see Tables S15 and S23.

**Table S23.** Main intermolecular interactions between the wild-type S-RBD$_{CoV-2}$ residue Q493 and all considered mutants at the protein-protein interface detected during MD simulations of the RBD of SARS-CoV-2 (COV-2) in complex with the LY-Cov016 antibody. Acronyms and other explanations as in Table S19.

| **SB** | **COV-2** | **LY-Cov016** | **Q493** | **H493** | **K493** | **L493** | **R493** |
|---|---|---|---|---|---|---|---|
| | K417 | $D_H$104 | ✓(2.92,3.10) | ✓(3.05,2.86) | ✓(2.92,3.00) | ✓(3.19,3.10) | ✓(3.40,3.25) |
| **HB** | **COV-2** | **LY-Cov016** | **Q493** | **H493** | **K493** | **L493** | **R493** |
| s-s | T415 | $S_H$56 | ✓(3.13) | ✗(p) | ✓(3.21) | ✓(3.18) | ✗(p) |
| **HB** | **COV-2** | **LY-Cov016** | **Q493** | **H493** | **K493** | **L493** | **R493** |
| s-s | K417 | $Y_H$52 | ✓(2.98) | ✓(3.14) | ✗(p) | ✗(p) | ✓(3.29) |
| **HB** | **COV-2** | **LY-Cov016** | **Q493** | **H493** | **K493** | **L493** | **R493** |
| s-s | D420 | $S_H$56 | ✓(3.07) | ✓(2.65) | ✓(2.67) | ✓(2.82) | ✓(2.66) |

| HB | COV-2 | LY-Cov016 | Q493 | H493 | K493 | L493 | R493 |
|---|---|---|---|---|---|---|---|
| s-s | Y421 | $S_H53$ | ✓(3.09) | ✓(3.45) | ✓(3.40) | ✓(3.38) | ✓(2.98) |
| s-b | Y421 | $G_H54$ | ✓(3.21) | ✓(3.19) | ✓(3.06) | ✓(2.92) | ✓(3.11) |
| **HB** | **COV-2** | **LY-Cov016** | **Q493** | **H493** | **K493** | **L493** | **R493** |
| s-b | N460 | $G_H54$ | ✓(3.12) | ✓(3.22) | ✓(3.20) | ✓(3.16) | ✓(3.24) |
| s-s | N460 | $S_H56$ | ✓(3.04) | ✓(3.07) | ✓(3.26) | ✓(3.11) | ✓(3.27) |
| **HB** | **COV-2** | **LY-Cov016** | **Q493** | **H493** | **K493** | **L493** | **R493** |
| s-b | X493 | $Y_H102$ | ✓(3.03) | ✓(3.37) | ✗(p) | ✗ | ✗(p) |
| **CI** | **COV-2** | **LY-Cov016** | **Q493** | **H493** | **K493** | **L493** | **R493** |
| p | K417 | $Y_H33$ | ✓ | ✓ | ✓ | ✓ | ✓ |
| vdW/h | K417 | $P_H100$ | ✓ | ✓ | ✓ | ✓ | ✓ |
| vdW/h | K421 | $Y_H52$ | ✓ | ✓ | ✓ | ✓ | ✓ |
| p | Y421 | $Y_H33$ | ✓ | ✓ | ✓ | ✓ | ✓ |
| vdW/h | X493 | $M_H101$ | ✓ | ✓ | ✓ | ✓ | ✓ |
| vdW/h | X493 | $Y_H102$ | ✓ | ✓ | ✓ | ✓ | ✓ |

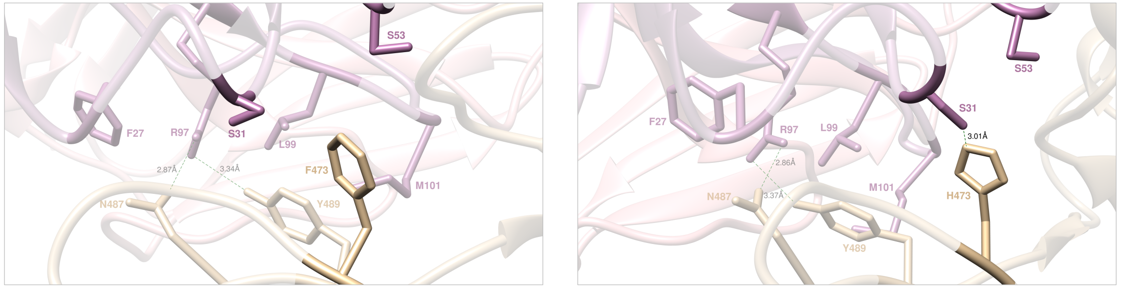


**Figure S16.** Main interactions involving the S-RBD$_{CoV-2}$ F473 (left) and H473 (right) at the interface with the LY-Cov016 antibody as obtained from the corresponding equilibrated MD simulations. The wild-type Y473 is presented and discussed in the main text (Figure 10B). Colors and other explanations as in Figure S11. For further details see Tables S15 and S24.

**Table S24.** Main intermolecular interactions between the wild-type S-RBD$_{CoV-2}$ residue Y473 and all considered mutants at the protein-protein interface detected during MD simulations of the RBD of SARS-CoV-2 (COV-2) in complex with the LY-Cov016 antibody. Acronyms and other explanations as in Table S19.

| **HB** | **COV-2** | **LY-Cov016** | **Y473** | **F473** | **H473** |
|---|---|---|---|---|---|
| s-s | X473 | $S_H$31 | ✓(2.83) | ✗(vdW/h) | ✓(3.01) |
| **HB** | **COV-2** | **LY-Cov016** | **Y473** | **F473** | **H473** |
| s-s | N487 | $R_H$97 | ✓(2.86) | ✓(2.87) | ✓(2.86) |
| **HB** | **COV-2** | **LY-Cov016** | **Y473** | **F473** | **H473** |
| s-s | Y489 | $R_H$97 | ✓(3.38) | ✓(3.34) | ✓(3.37) |
| **CI** | **COV-2** | **LY-Cov016** | **Y473** | **F473** | **H473** |
| p | X473 | $S_H$53 | ✓ | ✗(vdW/h) | ✓ |
| vdW/h | N487 | $F_H$27 | ✓ | ✓ | ✓ |
| vdW/h | Y489 | $L_H$99 | ✓ | ✓ | ✓ |
| vdW/h | Y489 | $M_H$101 | ✓ | ✓ | ✓ |

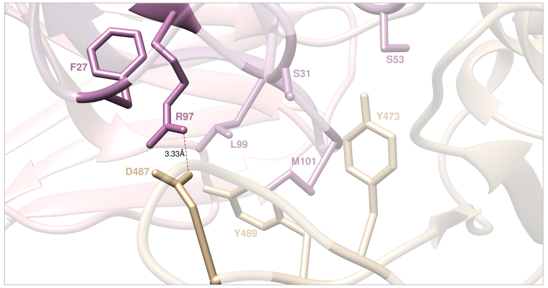


**Figure S17.** Main interactions involving the S-RBD$_{CoV-2}$ D487 at the interface with the LY-Cov016 antibody as obtained from the corresponding equilibrated MD simulations. The wild-type N487 is presented and discussed in the main text (Figure 10B). Colors and other explanations as in Figure S11. For further details see Tables S15 and S25.

**Table S25.** Main intermolecular interactions between the wild-type S-RBD$_{CoV-2}$ residue N487 and all considered mutants at the protein-protein interface detected during MD simulations of the RBD of SARS-CoV-2 (COV-2) in complex with the LY-Cov016 antibody. Acronyms and other explanations as in Table S19.

| **HB** | **COV-2** | **LY-Cov016** | **N487** | **D487** |
|---|---|---|---|---|
| s-s | Y473 | $S_H$31 | ✓(2.83) | ✗(p) |
| **HB** | **COV-2** | **LY-Cov016** | **N487** | **D487** |
| s-s | X487 | $R_H$97 | ✓(2.86) | ✓(SB,3.33) |
| **HB** | **COV-2** | **LY-Cov016** | N487 | D487 |
| s-s | Y489 | $R_H$97 | ✓(3.38) | ✗(p) |
| **CI** | **COV-2** | **LY-Cov016** | **N487** | **D487** |
| p | Y473 | $S_H$53 | ✓ | ✓ |
| vdW/h | X487 | $F_H$27 | ✓ | ✗ |
| vdW/h | Y489 | $L_H$99 | ✓ | ✓ |
| vdW/h | Y489 | $M_H$101 | ✓ | ✓ |

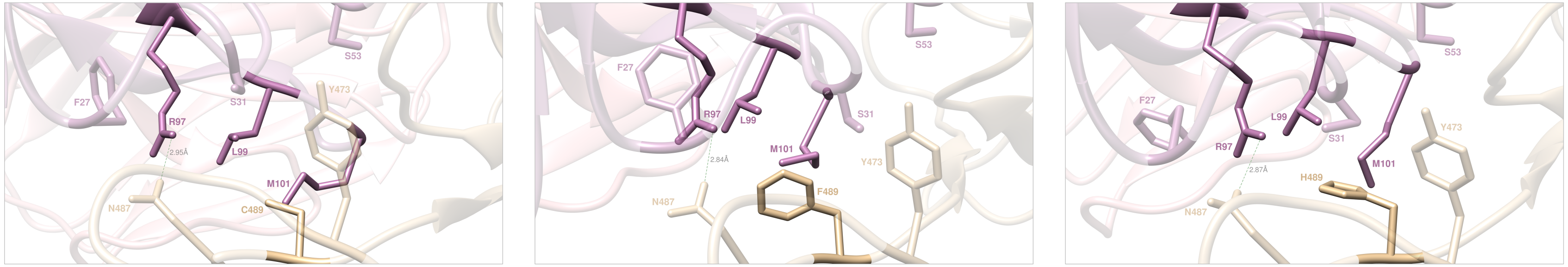


**Figure S18.** Main interactions involving the S-RBD$_{CoV-2}$ C489 (top), F489 (middle), and H489 (right) at the interface with the LY-Cov016 antibody as obtained from the corresponding equilibrated MD simulations. The wild type Y489 and the S489 mutant complexes are presented and discussed in the main text (Figures 10B and 14B). Colors and other explanations as in Figure S11. For further details see Tables S15 and S26.

**Table S26.** Main intermolecular and intramolecular interactions between the wild-type S-RBD$_{CoV-2}$ residue Y489 and all considered mutants* at the protein-protein interface detected during MD simulations of RBD of SARS-CoV-2 (COV-2) in complex with the LY-Cov016 antibody. Acronyms and other explanations as in Table S19. *Mutant S489 is discussed in detail in main text.

| **HB** | **COV-2** | **LY-Cov016** | **Y489** | **C489** | **F489** | **H489** | **S489** |
|---|---|---|---|---|---|---|---|
| s-s | Y473 | $S_H$31 | ✓(2.83) | ✗(p) | ✗(p) | ✗(p) | ✗(p) |
| **HB** | **COV-2** | **LY-Cov016** | **Y489** | **C489** | **F489** | **H489** | **S489** |
| s-s | N487 | $R_H$97 | ✓(2.86) | ✓(2.95) | ✓(2.84) | ✓(2.87) | ✓(2.99) |
| **HB** | **COV-2** | **LY-Cov016** | **Y489** | **C489** | **F489** | **H489** | **S489** |
| s-s | X489 | $R_H$97 | ✓(3.38) | ✗ | ✗(vdW/h) | ✗(p) | ✗ |
| **CI** | **COV-2** | **LY-Cov016** | **Y489** | **C489** | **F489** | **H489** | **S489** |
| p | Y473 | $S_H$53 | ✓ | ✓ | ✓ | ✓ | ✓ |
| vdW/h | N487 | $F_H$27 | ✓ | ✓ | ✓ | ✓ | ✓ |
| vdW/h | X489 | $L_H$99 | ✓ | ✗ | ✓ | ✓ | ✗ |
| vdW/h | X489 | $M_H$101 | ✓ | ✓ | ✓ | ✓ | ✓ |

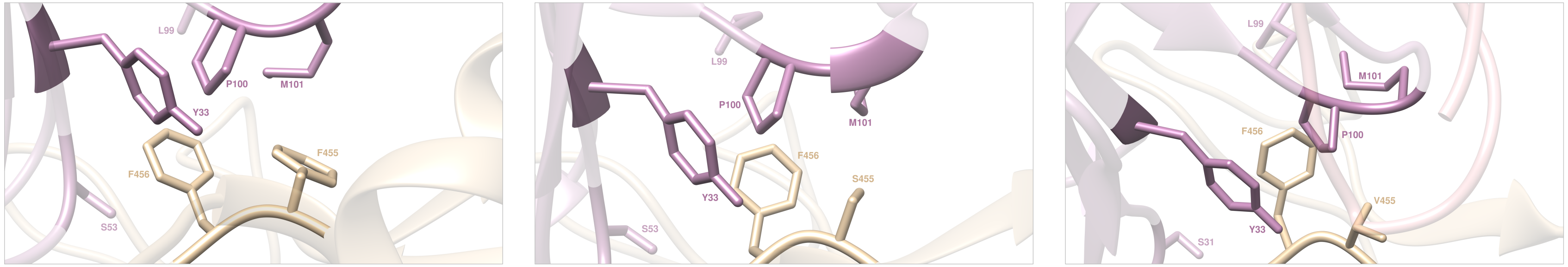


**Figure S19.** Main interactions involving the S-RBD$_{CoV\text{-}2}$ F455 (left), S455 (middle), and V455 (right) at the interface with the LY-Cov016 antibody as obtained from the corresponding equilibrated MD simulations. The wild-type L455 is presented and discussed in the main text (Figure 10C). Colors and other explanations as in Figure S11. For further details see Tables S15 and S27.

**Table S27.** Main intermolecular interactions between the wild-type S-RBD$_{CoV\text{-}2}$ residue L455 and all considered mutants at the protein-protein interface detected during MD simulations of the RBD of SARS-CoV-2 (COV-2) in complex with the LY-Cov016 antibody. Acronyms and other explanations as in Table S19.

| CI | COV-2 | LY-Cov016 | L455 | F455 | S455 | V455 |
|---|---|---|---|---|---|---|
| vdW/h | X455 | $Y_H$33 | ✓ | ✓ | ✓ | ✓ |
| vdW/h | X455 | $P_H$100 | ✓ | ✓ | ✗ | ✓ |
| vdW/h | X455 | $M_H$101 | ✓ | ✓ | ✗ | ✗ |
| **CI** | **COV-2** | **LY-Cov016** | **L455** | **F455** | **S455** | **V455** |
| vdW/h | F456 | $Y_H$33 | ✓ | ✓ | ✓ | ✓ |
| vdW/h | F456 | $S_H$53 | ✓ | ✓ | ✓ | ✓ |
| vdW/h | F456 | $L_H$99 | ✓ | ✓ | ✓ | ✓ |
| vdW/h | F456 | $P_H$100 | ✓ | ✓ | ✓ | ✓ |
| vdW/h | F456 | $M_H$101 | ✓ | ✓ | ✓ | ✓ |

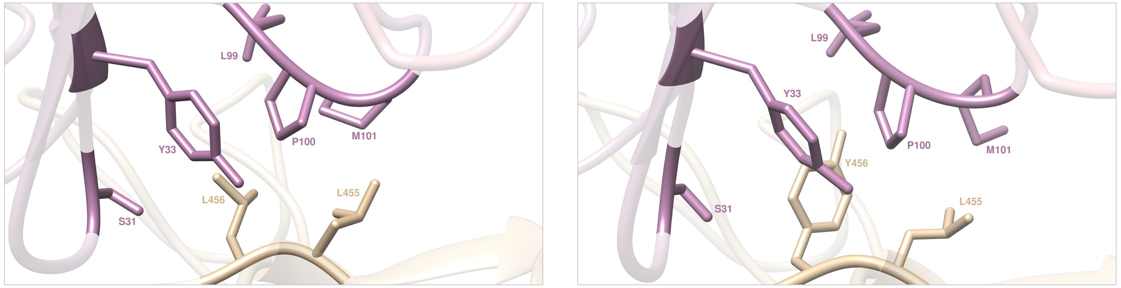


**Figure S20.** Main interactions involving the S-RBD$_{CoV-2}$ L456 (left) and Y456 (right) at the interface with the LY-Cov016 antibody as obtained from the corresponding equilibrated MD simulations. The wild-type F456 is presented and discussed in the main text (Figure 10C). Colors and other explanations as in Figure S11. For further details see Tables S1 and S28.

**Table S28.** Main intermolecular interactions between the wild-type S-RBD$_{CoV-2}$ residue F456 and all considered mutants at the protein-protein interface detected during MD simulations of the RBD of SARS-CoV-2 (COV-2) in complex with the LY-Cov016 antibody. Acronyms and other explanations as in Table S19.

| CI | COV-2 | LY-Cov016 | F456 | L456 | Y456 |
|---|---|---|---|---|---|
| vdW/h | L455 | $Y_H$33 | ✓ | ✓ | ✓ |
| vdW/h | L455 | $P_H$100 | ✓ | ✓ | ✓ |
| vdW/h | L455 | $M_H$101 | ✓ | ✓ | ✓ |
| **CI** | **COV-2** | **LY-Cov016** | **F456** | **L456** | **Y456** |
| vdW/h | X456 | $Y_H$33 | ✓ | ✓ | ✓ |
| vdW/h | X456 | $S_H$53 | ✓ | ✓ | ✓ |
| vdW/h | X456 | $L_H$99 | ✓ | ✗ | ✓ |
| vdW/h | X456 | $P_H$100 | ✓ | ✓ | ✓ |
| vdW/h | X456 | $M_H$101 | ✓ | ✓ | ✓ |

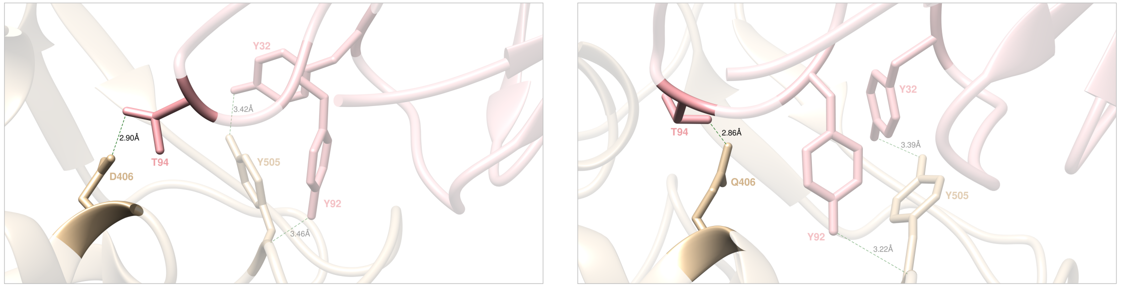


**Figure S21.** Main interactions involving the S-RBD$_{CoV-2}$ D406 (left) and Q456 (right) at the interface with the LY-Cov016 antibody as obtained from the corresponding equilibrated MD simulations. The wild-type E406 is presented and discussed in the main text (Figure 10D). Colors and other explanations as in Figure S11. For further details see Tables S15 and S29.

**Table S29.** Main intermolecular interactions between the wild-type S-RBD$_{CoV-2}$ residue D406 and all considered mutants at the protein-protein interface detected during MD simulations of the RBD of SARS-CoV-2 (COV-2) in complex with the LY-Cov016 antibody. Acronyms and other explanations as in Table S19.

| **HB** | **COV-2** | **LY-Cov016** | **E406** | **D406** | **Q406** |
|---|---|---|---|---|---|
| s-s | X406 | $T_L94$ | ✓(2.84) | ✓(2.90) | ✓(2.86) |
| **HB** | **COV-2** | **LY-Cov016** | **E406** | **D406** | **Q406** |
| s-s | Y505 | $Y_L32$ | ✓(3.01) | ✓(3.42) | ✓(3.39) |
| b-s | Y505 | $Y_L92$ | ✓(3.14) | ✓ (3.46) | ✓(3.22) |

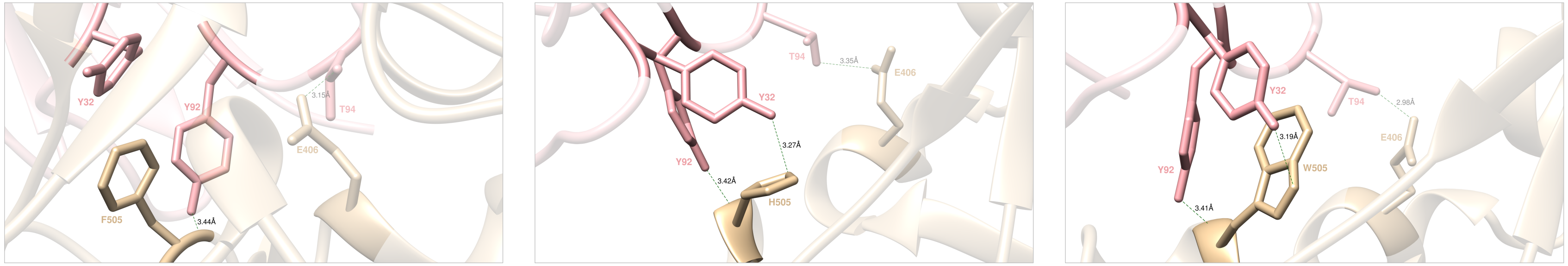


**Figure S22.** Main interactions involving the S-RBD$_{CoV-2}$ F505 (left), H505 (middle), and W505 (right) at the interface with the LY-Cov016 antibody as obtained from the corresponding equilibrated MD simulations. The wild-type Y505 is presented and discussed in the main text (Figure 10D). Colors and other explanations as in Figure S11. For further details see Tables S15 and S30.

**Table S30.** Main intermolecular interactions between the wild-type S-RBD$_{CoV-2}$ residue Y505 and all considered mutants at the protein-protein interface detected during MD simulations of the RBD of SARS-CoV-2 (COV-2) in complex with the LY-Cov016 antibody. Acronyms and other explanations as in Table S19.

| HB | COV-2 | LY-Cov016 | Y505 | F505 | H505 | W505 |
|---|---|---|---|---|---|---|
| s-s | E406 | $T_L$94 | ✓(2.84) | ✓(3.15) | ✓(3.35) | ✓(2.98) |
| **HB** | **COV-2** | **LY-Cov016** | **Y505** | **F505** | **H505** | **W505** |
| s-s | X505 | $Y_L$32 | ✓(3.01) | ✗(π/π) | ✓(3.27) | ✓(3.19) |
| b-s | X505 | $Y_L$92 | ✓(3.14) | ✓(3.44) | ✓(3.42) | ✓(3.41) |
