## Supporting information_part3 for "*In Silico* Molecular-Based Rationale for SARS-CoV-2 Spike Circulating Mutations Able to Escape Bamlanivimab and Etesevimab Monoclonal Antibodies"

The starting structure for the wild type SARS-CoV-2 S-protein receptor binding domain (S-$RBD_{CoV-2}$) in complex with either LY-CoV555 or LY-CoV016 were obtained from the RCSB Protein Data Bank[1] (PDB ID 7KMG[2] and 7C01[3], respectively). The physiological protonation state for all residues in each system was obtained by the H++ server[4] (http://biophysics.cs.vt.edu/H++).

The *tleap* software provided within AMBER20[5] was used for the parametrization of each system, assigning the ff14SB[6] and GLYCAM06j-1[7] forcefields to the protein and glycan structures. The complexes were next solvated in a box of TIP3PB[8] water molecules spanning at least 1.5 nm from each solute atoms. An appropriate number of $Na^+$ and $Cl^-$ atoms were added to neutralize the system and mimic the physiological salt concentration of 0.15 M. The following simulation scheme was applied for each simulated system. While applying a weak restraint (10 kcal/mol) on the proteins' backbone atoms, the simulation box was firstly energy minimized (this and the following minimization steps were composed of 3000 steps of steepest descent followed by 3000 steps of conjugated gradient algorithms), than heated to 150 K in 10 ps of canonical ensemble (NVT) molecular dynamics (MD), followed by another 50 ps MD simulation in the isothermal/isobaric ensemble (NPT, P = 1 atm, maintained by the Berendsen barostat[9]) to reach the target temperature of 300 K. The restraints were then gradually removed in 5 steps (-2 kcal/mol per step) of energy minimization. The MD simulation was next carried out without restraints for further 10 ns in NPT conditions (*phase 1*); after this time interval, the MD data production run was further continued up to 1 µs, during which pressure was maintained using the Monte Carlo barostat implemented in AMBER (*phase 2*). Along the entire MD trajectory, electrostatic interactions were computed by means of the particle mesh Ewald[10] (PME) algorithm, temperature was regulated by the Langevin method[11] (collision frequency of 3 $ps^{-1}$). The SHAKE algorithm[12] was applied to allow a 2 fs integration time step. All calculations were run with the *pmemd* module of AMBER20 running on the supercomputer Marconi100 (CINECA, Bologna, Italy) and on our CPU/GPU hybrid cluster. All images were produced by the UCSF Chimera software[13], VMD software[14] and on Prism 8 GraphPad Prism version 8.0.0 for Mac (GraphPad Software, San Diego, California USA, www.graphpad.com).

After the first 5 ns of the *phase 2* MD trajectory, 5 ns MD data were selected to calculate enthalpy and entropy contributions. Configurational sampling in this part of the simulation was preformed accordingly, with a time step of 10 fs; thus, a total of 500 000 snapshots, sufficient for the interaction entropy (IE) calculations[15-17], were extracted from the relevant MD trajectory for the calculation of the protein/protein residue- specific interactions. The free energy was calculated for each molecular species in the framework of the MM/PBSA ansatz[18], and the protein/antibody (AB) binding free energy was computed as the difference:

$$\Delta G = G_{S-RBD_{COV2}/AB} - \left(G_{S-RBD_{COV2}} + G_{AB}\right) = \Delta E_{vdW} + \Delta E_{ELE} + \Delta G_{SOL} - T\Delta S = \Delta H - T\Delta S$$

Here $\Delta E_{vdW}$ and $\Delta E_{ELE}$ represent van der Waals and electrostatic molecular mechanics energies, and $\Delta G_{SOL}$ includes the solvation free energy. The internal dielectric constant was set to the values of 2, 3 and 9 for nonpolar, polar, and charged residues, respectively[15, 19]. Lastly, the entropic contribution ($T\Delta S$) was explicitly computed from the MD simulation by using the Interaction Entropy (IE) method [15-17].

The role of the protein/protein interface key residues was studied by performing computational alanine scanning (CAS) experiments[20]. Accordingly, the absolute binding free energy of each mutant receptor - in which each key residue was replaced by alanine by truncating the mutated residue at the $C_\gamma$ atom, and replacing it with a hydrogen - was calculated with the MM/PBSA method. Accordingly, the difference in the binding free energy between the wild-type (WT) protein and its alanine mutant (ALA) counterpart, $\Delta\Delta G_{CAS}$, is given by:

$$\Delta\Delta G_{CAS} = \Delta G_{WT} - \Delta G_{ALA}$$

Thus, the CAS methodology allows for the estimation of the contribution of a given residue with respect to the overall protein−protein binding free energy; indeed, according to the equation, a negative value of $\Delta\Delta G_{CAS}$ indicated a favorable contribution for the wild type residue in that position and vice versa.

The role of the protein/AB interface key residues mutations was calculated with the MM/PBSA method. Accordingly, the difference in the binding free energy between the wild-type (WT) protein and mutant counterpart, $\Delta\Delta G$, is given by:

$$\Delta\Delta G = \Delta G_{WT} - \Delta G_{MUTANT}$$

Thus, the adopted methodology allows for the estimation of the contribution of a given residue with respect to the overall protein/AB binding free energy; indeed, according to the equation, a negative value of $\Delta\Delta G$ indicated a favorable contribution for the wild type residue in that position and vice versa.

At the structural level, the stability of the main protein/protein interface intermolecular and intramolecular interactions detected during the MD simulation time interval adopted for the energetic analysis was assessed along the entire duration of the MD run.
